# Calcium-permeable AMPA receptors govern PV neuron feature selectivity

**DOI:** 10.1101/2023.07.20.549908

**Authors:** Ingie Hong, Juhyun Kim, Thomas Hainmueller, Dong Won Kim, Richard C. Johnson, Soo Hyun Park, Nathachit Limjunyawong, Zhuonan Yang, David Cheon, Taeyoung Hwang, Amit Agarwal, Thibault Cholvin, Fenna M. Krienen, Steven A. McCarroll, Xinzhong Dong, David A. Leopold, Seth Blackshaw, Dwight E. Bergles, Marlene Bartos, Solange P. Brown, Richard L. Huganir

## Abstract

The brain helps us survive by forming internal representations of the external world^1,2^. Excitatory cortical neurons are often precisely tuned to specific external stimuli^3,4^. However, inhibitory neurons, such as parvalbumin-positive (PV) interneurons, are generally less selective^5^. PV interneurons differ from excitatory cells in their neurotransmitter receptor subtypes, including AMPA receptors^6,7^. While excitatory neurons express calcium-impermeable AMPA receptors containing the GluA2 subunit, PV interneurons express receptors that lack the GluA2 subunit and are calcium-permeable (CP-AMPARs). Here we demonstrate a causal relationship between CP-AMPAR expression and the low feature selectivity of PV interneurons. We find a low expression stoichiometry of GluA2 mRNA relative to other subunits in PV interneurons which is conserved across ferrets, rodents, marmosets, and humans, causing abundant CP-AMPAR expression. Replacing CP-AMPARs in PV interneurons with calcium-impermeable AMPARs increased their orientation selectivity in the visual cortex. Sparse CP-AMPAR manipulations demonstrated that this increase was cell-autonomous and could occur well beyond development. Interestingly, excitatory-PV interneuron connectivity rates and unitary synaptic strength were unaltered by CP-AMPAR removal, suggesting that the selectivity of PV interneurons can be altered without drastically changing connectivity. In GluA2 knockout mice, where all AMPARs are calcium-permeable, excitatory neurons showed significantly reduced orientation selectivity, suggesting that CP-AMPARs are sufficient to drive lower selectivity regardless of cell type. Remarkably, hippocampal PV interneurons, which usually exhibit low spatial tuning, became more spatially selective after removing CP-AMPARs, indicating that CP-AMPARs suppress the feature selectivity of PV interneurons independent of modality. These results reveal a novel role of CP-AMPARs in maintaining a low-selectivity sensory representation in PV interneurons and suggest a conserved molecular mechanism that distinguishes the unique synaptic computations of inhibitory and excitatory neurons.

## Introduction

Genes define a neuron’s response to synaptic input and thus program its biophysical computations^6,8^. For instance, neurotransmitter receptor profiles can decide the influx of Ca^2+^ to dendrites, triggering neuronal changes that enable information storage^1,9,10^. Interestingly, gene expression varies widely across neuron types, prompting distinct responses to the same sensory input and specialized roles within a given network^11,12^. A key gene expression difference among the cardinal neuron types lies in their synaptic receptor composition^13,14^. Despite significant advances in understanding the role of synaptic genes’ plasticity, how these genes affect computations in the native brain is vastly underexplored.

Neurons in the neocortex exquisitely compute and represent features of the outside world through sparse, de-correlated activity^2^. This capability is particularly well-characterized in the hippocampus, where place cells fire strongly to specific locations^3^, and in the primary visual cortex, where neurons are highly tuned to oriented edges or movement directions in the visual receptive field^4^. In the visual cortex, a neuron’s response selectivity for orientation, spatial frequency, color, or speed is conferred through organized synaptic inputs arising from relay neurons in the thalamus and excitatory/inhibitory neurons from within the cortex^4,15^. However, the genes, receptors, and plasticity programs giving rise to this finely tuned circuit organization are largely unknown. A critical clue to this question arises from the natural diversity of feature tuning expressed by different cell types^16^. Excitatory glutamatergic neurons display sharp orientation and direction selectivity, whereas parvalbumin-expressing (PV) GABAergic interneurons show low orientation selectivity^17-21^ (but see refs. ^22,23^). This distinction appears general across various modalities, with PV interneurons typically displaying less selectivity. For instance, in the hippocampus, PV cells adjacent to excitatory ‘place’ cells show much lower spatial selectivity^5^.

PV basket cells are exquisitely adapted to fast spiking and provide robust and rapid feedback inhibition to nearby neurons^5^. They possess a distinct glutamate receptor profile, with small N-methyl-D-aspartate receptor (NMDAR) currents but large inwardly rectifying calcium-permeable AMPA (α-amino-3-hydroxy-5-methyl-4-isoxazolepropionic acid) receptor (CP-AMPAR) currents^7,24-28^. AMPARs are tetrameric glutamate receptors that serve the majority of fast Hong et al., 2023 (preprint) excitatory synaptic transmission in the brain. CP-AMPARs lack the voltage-dependent block of NMDARs by Mg^2+^, allowing them to induce distinct Ca^2+^ dynamics and forms of plasticity at synapses^29-33^. This characteristic suggests that CP-AMPARs in PV interneurons have the potential to continually shape and adjust the relative strengths of inputs carrying different types of information in a manner distinct from nearby excitatory neurons. Here we explore the consequences of high CP-AMPAR levels and reveal they play a striking role in maintaining the low orientation selectivity of PV interneurons.

### Conserved AMPAR subunit mRNA stoichiometry

The calcium permeability of an AMPAR arises from two distinct routes, both involving GluA2/Gria2: the lack of a GluA2 subunit (GluA2-lacking AMPAR) or the lack of RNA editing in the Gria2 gene site encoding a critical pore amino acid of GluA2 (unedited GluA2)^25,34^. The lack of GluA2, in turn, could be due to transcriptional or translational regulation. We first investigated the mechanism that drives high CP-AMPAR expression in PV interneurons and identified transcriptional regulation conserved across multiple mammalian species.

Analysis of SmartSeq-based high-coverage single-cell RNA-seq data from the mouse cortex^14^ revealed that RNA editing at the GluA2 Q/R site was uniformly complete (>95%) in all neuronal cell types (Extended Data Fig. 1), suggesting that unedited GluA2 does not significantly contribute to CP-AMPAR expression in PV interneurons.

Validated antibodies against the major forebrain glutamate receptor subunits GluA1 and GluA2 (Extended Data Fig. 2) showed PV interneurons expressing GluA2 at roughly 60% of the levels of neighboring CaMKIIα+ excitatory neurons in both mice (Fig. 1a,b) and marmosets (Fig. 1c,d). Conversely, GluA1 was expressed ∼1.7 fold higher in PV interneurons, likely contributing to a lower GluA2:GluA1 ratio and abundant GluA2-lacking CP-AM-PARs (Extended Data Fig. 3).

**Figure 1.**
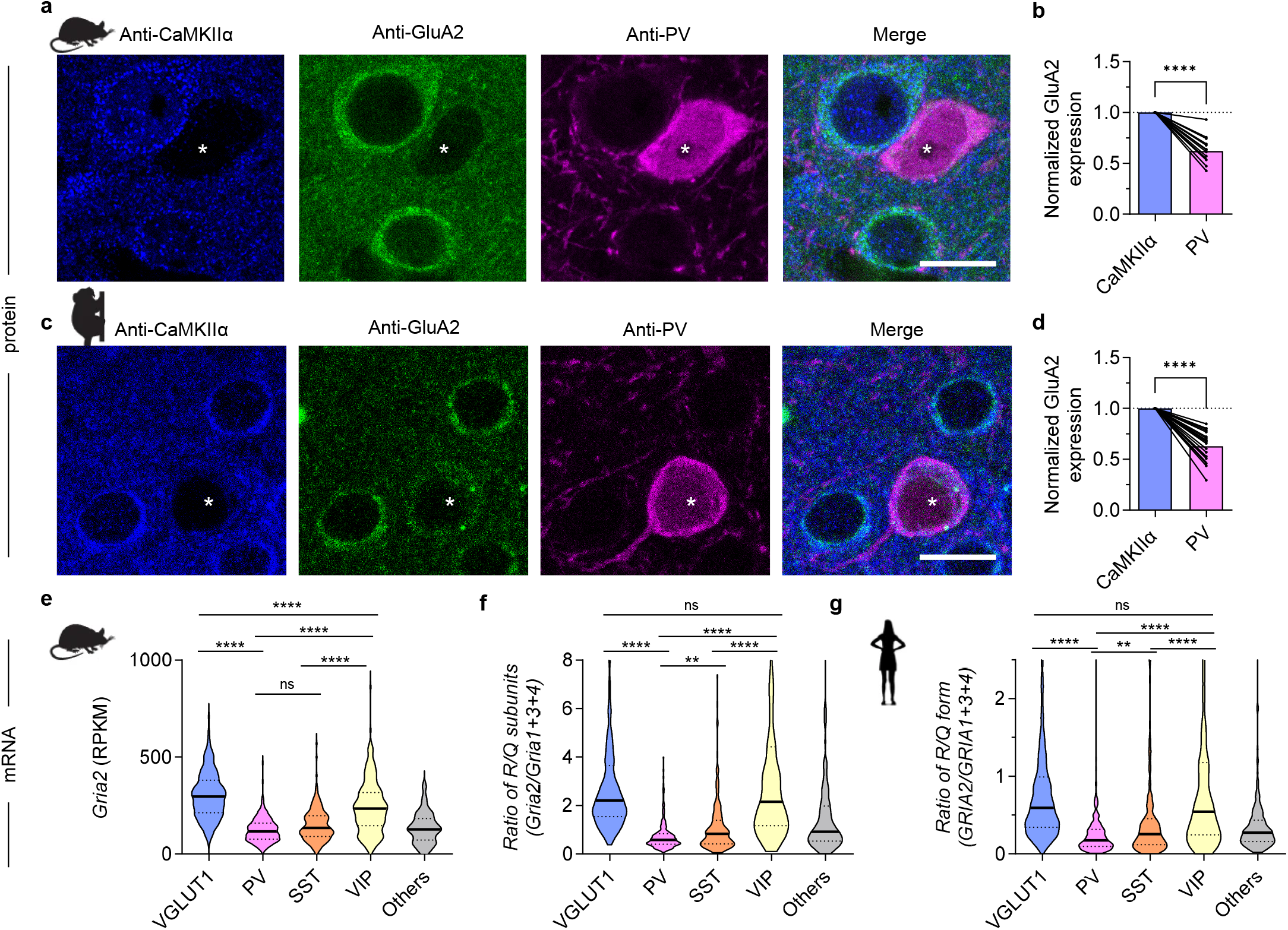
Selective low expression of GluA2 and *Gria2* in PV and SST interneurons in mice, marmosets, and humans. **a**, Immunohistochemical staining of PV and GluA2 in visual cortex layer 2/3. PV interneurons (asterisks) show markedly lower GluA2 expression compared to CaMKIIα excitatory counterparts. Layer 2/3 of visual cortex, scale bars, 10 μm. **b**, Quantification of relative GluA2 expression as a ratio of PV/CaMKIIα neurons (mean ± SEM) shows that PV interneurons express significantly less GluA2 (PV: 0.62 ± 0.03-fold vs CaMKIIα; n = 15 neuron pairs from 3 slices, 3 mice; P < 0.0001, 1-sample t-test). **c**, GluA2 expression in PV interneurons and CaMKIIα excitatory neurons in the marmoset cortex. Scale bars, 10 μm. **d**, Marmoset PV interneurons express significantly less GluA2 compared to nearby CaMKIIα neurons (PV: 0.63 ± 0.03fold vs CaMKIIα; n = 22 pairs from 7 slices, 3 marmosets, P < 0.0001, one sample t-test). Bars and error bars denote mean ± SEM. **e**, Analysis of Smart-seq single-cell RNA-seq data14 from the visual cortex of p56 mice shows distinctly lower expression of *Gria2* mRNA in PV and SST interneurons (n = 756/270/178/185/118 neurons from VGLUT1/PV/SST/VIP/Other cell types, respectively, H(4) = 610.9, P < 0.0001, KW one-way ANOVA; P < 0.0001 for all VGLUT1 post-hoc comparisons, Dunn’s multiple comparison correction). A fraction of outlier cells were omitted for visualization. Conventional marker protein names are adapted to denote cardinal neuronal cell classes (VGLUT1 neurons and CaMKIIα neurons both refer to forebrain excitatory neurons). Post-hoc comparisons with the ‘others’ group are omitted for brevity. **f**,**g**, This low expression of *Gria2* contributes to the lower ratio of calcium impermeable/calcium permeable AMPAR subunits (R/Q subunit ratio) both in mice (**f**) and in humans35 (**g**). In both (**f**) and (**g**), a KW one-way ANOVA test reveals a significant difference (mice: H(4) = 593.6, P < 0.0001; humans: H(4) = 491.9, P < 0.0001), and post-hoc comparisons demonstrate significant differences between all non-’others’ pairs except VGLUT vs. VIP (panel **g** shows human data from n = 2151/235/193/282/181 neurons from VGLUT1/PV/SST/VIP/Other cell types, respectively). Post-hoc comparisons with the ‘others’ group are omitted for brevity. Thick center lines and dotted lines in violin plots represent median and 25-75% interquartile range, respectively.

Single-cell data on mRNA expression^14^ for the genes encoding AMPAR subunits closely matched the protein level data, with PV interneurons expressing GluA2 mRNA at about 40% of its express level in neighboring excitatory neurons (VGLUT1+, Fig. 1e), suggesting transcriptional regulation of CP-AMPARs. SST interneurons are GABAergic, with the same developmental origin (MGE, medial ganglionic eminence). These interneurons also expressed lower GluA2. However, VIP+ interneurons, which originate separately from the CGE (caudal ganglionic eminence), displayed GluA2 mRNA levels similar to excitatory neurons (Fig. 1e). Motivated by the tightly correlated mRNA expression of AMPAR subunits (Extended Data Fig. 4), we calculated the ratio of GluA2:GluA1+3+4, which reflects the relative levels of calcium impermeable:permeable (R:Q) subunits and found a similar ratio profile (Fig. 1f). Strikingly, this GluA2:GluA1+3+4 mRNA ratio was always lowest in PV interneurons across ferret, mouse, marmoset, macaque, and human cortex datasets^35,36^ (Fig. 1g and Extended Data Fig. 5), suggesting evolutionary pressure toward lower R/Q ratios specifically in these neurons. These results show that PV inhibitory neurons across these species express GluA2 at a tightly regulated low expression stoichiometry, likely through a transcriptional mechanism strongest in PV interneurons^7^.

### Manipulation of CP-AMPARs in PV neurons

To test the functional significance of low GluA2 and high CP-AMPAR expression, we used a recently generated transgenic mouse^37^ to express additional GluA2 with an eGFP tag selectively in PV interneurons a Cre-dependent fashion (Extended Data Fig. 6). We crossed this transgenic mouse line with PV-Cre knock-in mice^38^, generating PV-Cre;lsl-eGFP-GluA2 mice (Fig. 2a). The cross robustly expressed GluA2 at the cell soma and along the dendrites of PV interneurons (Extended Data Fig. 7c and Fig. 2b) at high concordance with PV immunofluorescence (Extended Data Fig. 8). The transgenic expression of GluA2 led to PV interneurons with levels of GluA2 comparable to excitatory neurons, roughly twice the level of PV-Cre;lsl-eGFP control mice. This increased expression occurred both at the mRNA level (221.6% ± 26.9%; Extended Data Fig. 18a) and protein level (217.2 ± 11.2%; Fig. 2b and Extended Data Fig. 7a-c), revealed by FACS-assisted PV interneuron RNA-seq (Methods) and immunohistochemistry, respectively. Notably, GluA1 protein (but not mRNA) expression was lower in PV interneurons expressing exogenous GluA2 (Fig. 2c and Extended Data Fig. 7d-f), similar to excitatory neurons, suggesting a posttranscriptional homeostatic or displacing effect. Transgenic Hong et al., 2023 (preprint) expression of GluA2 in PV cells did not yield significant changes in PV or SST interneuron density in the visual cortex (Extended Data Fig. 9).

**Figure 2.**
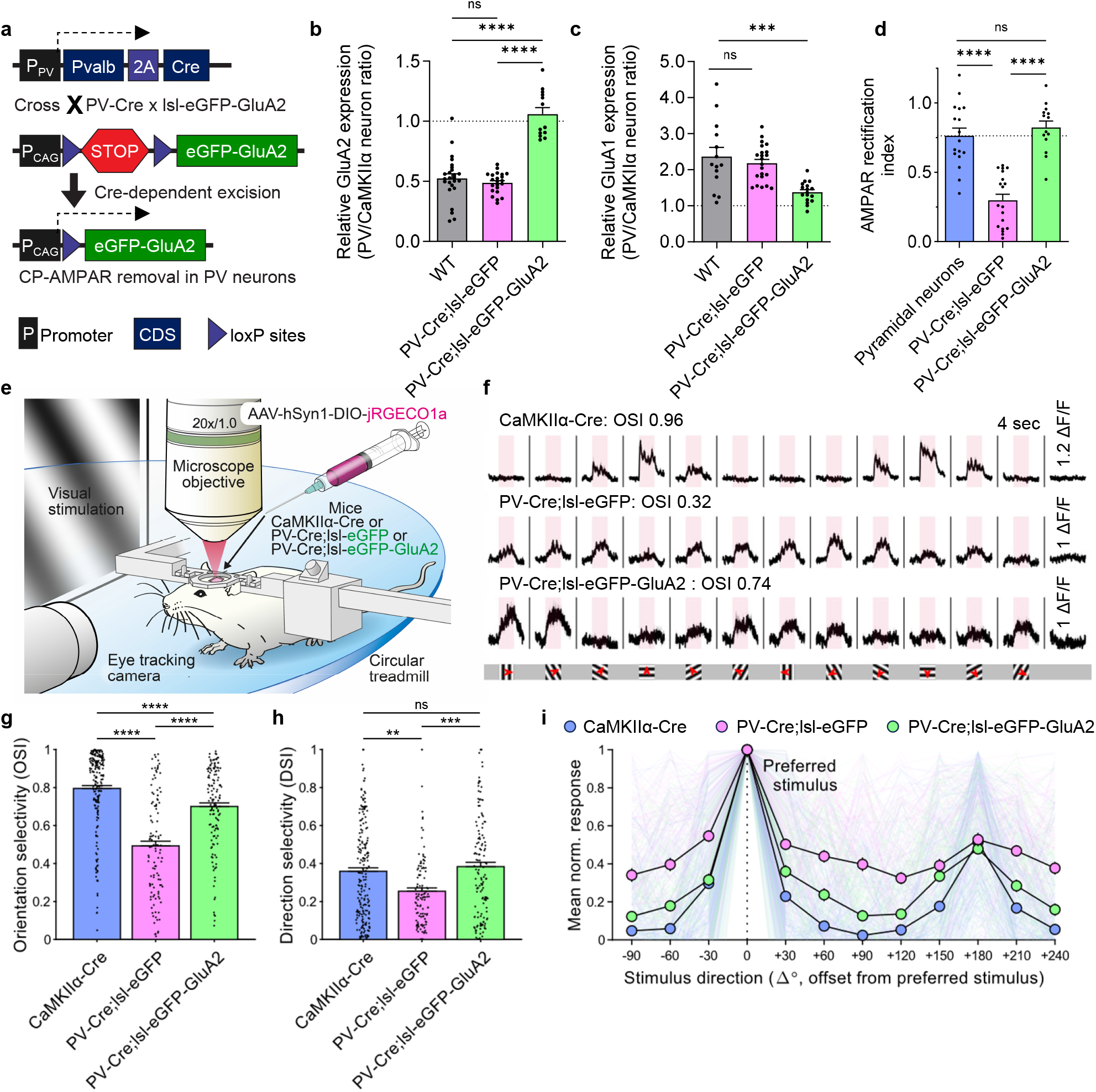
GluA2 expression in PV interneurons alters orientation selectivity in layer 2/3 of mouse visual cortex. **a**, Strategy for removing CP-AMPARs selectively in PV interneurons. **b**, Quantification of relative GluA2 protein expression as a ratio of PV/CaMKIIα neurons (n = 25/22/13 pairs from 4/4/4 slices, 3/3/3 mice, P < 0.0001, KW one-way ANOVA; P < 0.0001 for all PV-Cre;lsl-eGFP-GluA2 post-hoc comparisons, Dunn’s multiple comparison correction). **c**, Relative GluA1 protein expression (n = 14/22/17 pairs from 3/3/3 slices, 3/3/3 mice, P < 0.0001, one-way ANOVA; P < 0.001 for all PV-Cre;lsl-eGFP-GluA2 post-hoc comparisons, Tukey’s multiple comparison correction). Bars and error bars denote mean ± SEM. **d**, The low AMPAR rectification index in PV control neurons (PV-Cre;lsl-eGFP, 0.298 ± 0.044) is increased in PV-Cre;lsl-eGFP-GluA2 mice (0.823 ± 0.047) to levels comparable with pyramidal neurons (0.763 ± 0.056) recorded for comparison (n = 17/19/14 cells from 4/3/2 mice, P < 0.0001, one-way ANOVA test; P < 0.0001 for all post-hoc comparisons with PV-Cre;lsl-eGFP, Tukey’s multiple comparison correction). This indicates the removal of calcium-permeable AMPARs by eGFP-GluA2 expression. **e**, Pre-injected mice were head-fixed and visually stimulated during 2P imaging of V1 to reveal differences in tuning. **f**, Representative soma activity traces of CaMKIIα, PV-Cre;lsl-eGFP, and PV-Cre;lsl-eGFP-GluA2 neurons. Pink rectangles denote the 4s visual stimulation period and 1.2/1.0/1.0 ΔF/F for each group. Grey shading corresponds to SEM. Whole screen drifting grating stimulation with 12 different orientations were used to assess orientation selectivity. Red arrows mark the drifting direction. **g**,**h**, Quantification of orientation and direction selectivity. CaMKIIα neurons in the CaMKIIα-Cre mice displayed higher orientation selectivity compared to PV interneurons in PV-Cre;lsl-eGFP mice, and the PV-Cre;lsl-eGFP-GluA2 group showed higher OSI than PV-Cre;lsl-eGFP controls (n = 202/114/137 neurons from 4/4/5 mice, H(2) = 99.10, P < 0.0001, KW one-way ANOVA; P < 0.0001 for all post-hoc comparisons, Dunn’s multiple comparison correction). The CaMKIIα-Cre group displayed higher direction selectivity compared to the PV-Cre;lsl-eGFP group, and the PV-Cre;lsl-eGFP-GluA2 group showed higher DSI than PV-Cre;lsl-eGFP controls (H(2) = 15.91, P = 0.0004, KW one-way ANOVA; P = 0.0029 for CaMKIIα-Cre vs. PV-Cre;lsl-eGFP, P = 0.0006 for PV-Cre;lsl-eGFP vs. PV-Cre;lsl-eGFP-GluA2). **i**, Normalized average response profile of all positively responding neurons from each group, aligned to their preferred stimulus direction (0°). Note that a prominent peak is also present at +180°, due to the orientation selective nature of V1 neurons. Responses are plotted as mean ± SEM.

Electrophysiological recordings revealed the signatureinward rectification of CP-AMPARs typical of PV interneurons was absent in PV-Cre;lsl-eGFP-GluA2 mice (Fig. 2d; Extended Data Fig. 10), showing synaptic incorporation of calcium-impermeable AMPARs. These results validate the PV-Cre;lsl-eGFP-GluA2 mouse model to test the functional role of CP-AMPARs in PV interneurons.

### CP-AMPARs suppress PV selectivity

To test the role of CP-AMPARs in PV interneurons on sensory representation in awake mice, we assessed the orientation preference of layer 2/3 (L2/3) neurons in the primary visual cortex (V1) with *in vivo* two-photon (2P) calcium imaging (Fig. 2e). We injected PV-Cre;lsl-eGFP-GluA2 mice, PV-Cre;lsl-eGFP mice as controls, and CaMKIIα-Cre mice for comparison, with a Cre-dependent jRGECO1a AAV targeting L2/3 of the visual cortex. Cranial windows were implanted on these mice and retinotopic mapping was performed to map the monocular primary visual cortex for 2P imaging (Extended Data Fig. 11a-e). We imaged soma Ca^2+^ activity as a proxy for action potential activity during drifting grating presentation, focusing on the impact of neuronal firing rather than dendritic calcium dynamics^39,40^. A portion of PV interneurons was visually responsive (Extended Data Fig. 11f), displaying robust activity toward 4-sec presentations of full-field drifting gratings but not to blank isoluminant grey screen control trials (Fig. 2f). Consistent with previous observations^17-21^, the orientation selectivity of L2/3 PV interneurons was lower than excitatory neurons (with considerable variability in both populations^4,22,23^). Selectivity was significantly enhanced when GluA2 expression matched the level in excitatory neurons (Fig. 2g; H_(2)_ = 99.10, P < 0.0001, KW 1-way ANOVA; P < 0.0001 for CaMKIIα-Cre vs. PV-Cre;lsl-eGFP and PV-Cre;lsl-eGFP vs. PV-Cre;lsl-eGFP-GluA2, Dunn’s multiple comparison correction). Despite distinct circuit underpinnings^4,15,16,19,41-43^, direction selectivity was similarly enhanced, suggesting a general increase in selectivity as a result of CP-AMPAR removal (Fig. 2h; H_(2)_ = 99.10, P < 0.0001, KW 1-way ANOVA; P = 0.0029 for CaMKIIα-Cre vs. PV-Cre;lsl-eGFP and P = 0.0006 for PV-Cre;lsl-eGFP vs. PV-Cre;lsl-eGFP-GluA2). However, the average response amplitude was not significantly changed (Extended Data Fig.11g). The average tuning curve demonstrates how relative non-preferred stimuli responses are reduced in PV interneurons without CP-AMPARs, yielding increased orientation and direction selectivity (Fig. 2i and Extended Data Fig. 11h-j).

### Cell-autonomous effect of CP-AMPAR removal

These results suggested that CP-AMPARs help lower the visual feature selectivity of PV interneurons. We asked whether this effect arose from the systemic expression of GluA2 in PV interneurons or a cell-autonomous effect. To test the cell-autonomous effect of CP-AMPAR removal, we developed an adeno-associated virus (AAV) vector to sparsely express SEP (super-ecliptic pHluorin)-tagged GluA2 in a Cre-dependent fashion (see Methods). Using this AAV-DIO-SEP-GluA2 virus, we observed robust expression of SEP-GluA2 in cultured neurons and *in vivo* (Extended Data Fig. 12a, b). As expected, rectification measurements demonstrated robust removal of CP-AMPARs in PV interneurons compared with the control AAV-DIO-eGFP (Extended Data Fig. 12c-f). To further control for increased GluA2 expression, we used an AAV expressing the calcium-permeable form of GluA2 (AAV-DIO-SEP-GluA2Q). This AAV similarly supplements GluA2 and should increase the portion of CP-AMPARs. Indeed, this virus increased the rectification in PV interneurons (Extended Data Fig. 12c, g).

Sparsely expressing GluA2 in PV interneurons increased orientation selectivity, suggesting the effect of CP-AMPAR removal is cell-autonomous (Fig. 3a-c and Extended Data Fig. 13; H_(2)_ = 11.84, P = 0.0027, KW 1-way ANOVA; P = 0.0028 for eGFP vs. SEP-GluA2 OSI, Dunn’s multiple comparison correction). Interestingly, this effect was observed even in a >8-month-old mouse (P < 0.0001, Mann-Whitney U-test), consistent with an ongoing role of CP-AMPARs in suppressing orientation selectivity. Note that the calcium-permeable form of GluA2 (SEP-GluA2Q) did not increase PV interneuron orientation selectivity (P = 0.9723 for eGFP vs. SEP-GluA2Q), suggesting a specific role of the calcium permeability of the channel pore. SEP-GluA2Q expression did not lead to a decrease in orientation selectivity either, suggesting saturation or a floor effect.

**Figure 3.**
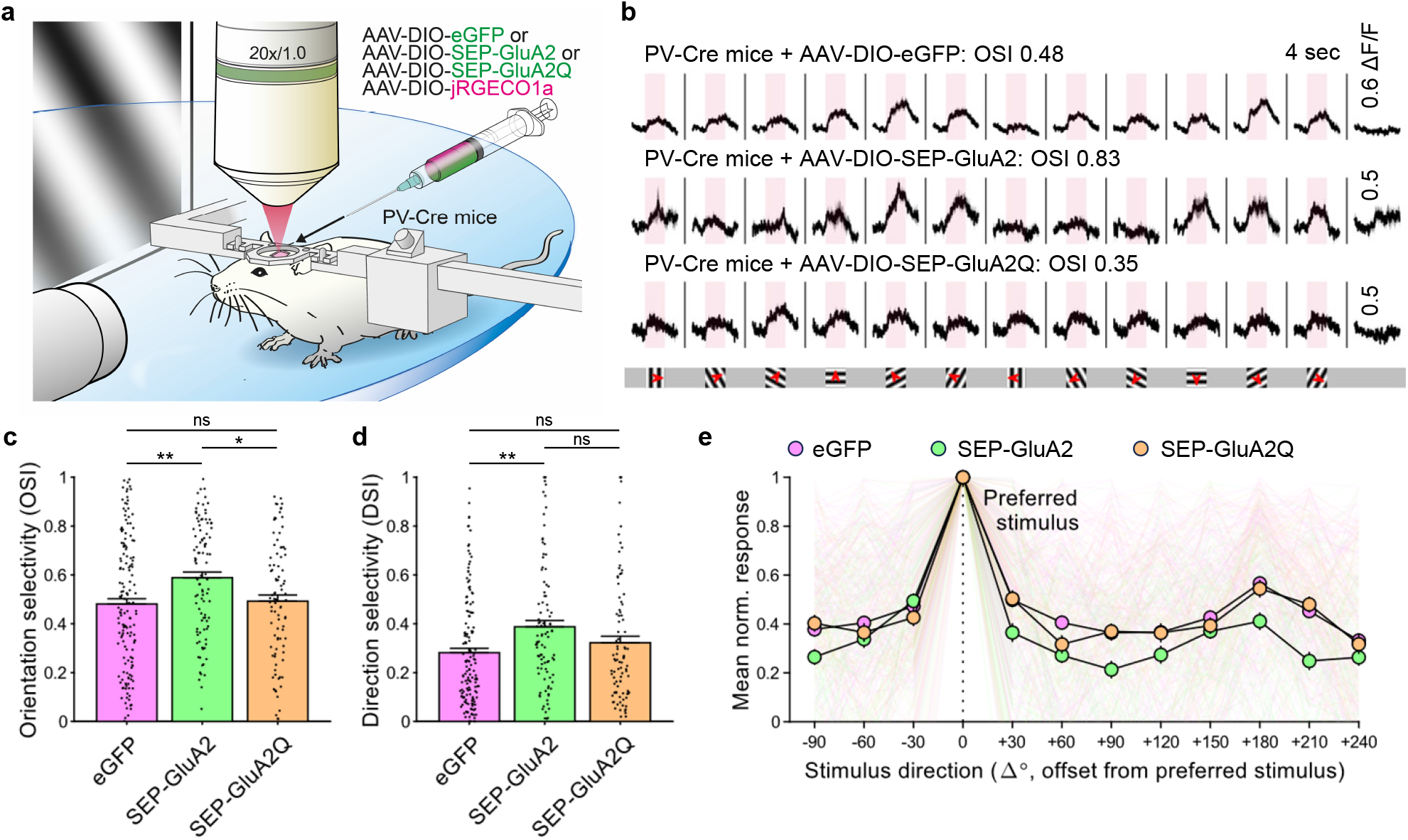
Sparse GluA2 expression in layer 2/3 PV interneurons increases their orientation and direction selectivity. **a**, Pre-injected mice were head-fixed and visually stimulated during 2P imaging of V1 to reveal differences in tuning. **b**, Representative traces of neurons infected with AAV-DIO-eGFP and AAV-DIO-SEP-GluA2. Pink regions denote the 4s visual stimulation period and 0.6/0.5/0.5 ΔF/F for each group. Whole screen drifting grating stimulation with 12 different orientations was used to assess orientation selectivity. Red arrows mark the drifting direction. **c**, The SEP-GluA2 group displayed higher orientation selectivity compared to eGFP controls, whereas the SEP-GluA2Q group did not show increased OSI (n = 154/100/91 neurons from 4/4/3 mice, H(2) = 11.84, P = 0.0027, KW one-way ANOVA; P = 0.0028 for eGFP vs. SEP-GluA2, P = 0.9723 for eGFP vs. SEP-GluA2Q, Dunn’s multiple comparison correction). **d**, Similarly, the SEP-GluA2 group displayed higher direction selectivity compared to the eGFP control group, whereas the SEP-GluA2Q group failed to show higher DSI (P = 0.0016, H(2) = 12.84, KW one-way ANOVA; P = 0.0010 for eGFP vs. SEP-GluA2, P = 0.5558 for eGFP vs. SEP-GluA2Q). **e**, Normalized average response profile of all positively responding neurons, aligned to their preferred stimulation (0°). Responses are plotted as mean ± SEM.

Sparse SEP-GluA2 expression increased direction selectivity, but SEP-GluA2Q did not (Fig. 3d; P = 0.0016, H_(2)_ = Hong et al., 2023 (preprint) 12.84, KW 1-way ANOVA; P = 0.0010 for eGFP vs. SEP-GluA2, P = 0.5558 for eGFP vs. SEP-GluA2Q), suggesting that CP-AMPARs are necessary for PV interneurons’ low selectivity to various visual features. The proportion of visually responsive neurons and average response amplitude were not significantly different (Extended Data Fig. 14a,b). The average tuning curve confirms reduced responses to non-preferred stimuli in SEP-GluA2-expressing PV interneurons (Fig. 3e).

### CP-AMPARs shape excitability, but not connectivity

We next investigated the role of circuit changes in orientation selectivity increase in PV interneurons. Excitatory neurons have much sparser input connectivity with local neurons compared to PV interneurons which may underlie their sharper orientation selectivity^43-46^. Thus, one hypothesis is that PV interneurons’ dense local excitatory input connections become sparser following removal of CP-AMPARs to resemble the selective inputs of excitatory neurons. Alternatively, the nominal connectivity rate may be unchanged, leaving more specific mechanisms to account for selectivity change. We used paired whole-cell patch-clamp recordings in acute brain slices to test the connectivity of L2/3 PV interneurons and nearby excitatory neurons. We found no significant changes in the excitatory input nor in reciprocal inhibitory output connection probability (Extended Data Fig. 14). In connected pairs, the excitatory-to-PV interneuron unitary EPSP amplitudes were not significantly smaller (Extended Data Fig. 14f), despite the loss of high-conductance CP-AM-PARs, suggesting a feedback or homeostatic mechanism preserving synaptic strength. These results reject a large change in excitatory input connectivity as a mechanism for increasing orientation selectivity but do not exclude the possibility that presynaptic input reorganization could lead to higher selectivity.

Input reorganization could arise from altered synaptic plasticity in PV interneurons due to the removal of CP-AM-PARs, which mediate an anti-Hebbian form of LTP in hippocampal PV interneurons through their ability to allow calcium influx at polarized potentials^30^. Synaptic plasticity has been explored much less in cortical PV interneurons and generally presents as LTD or smaller LTP compared to hippocampal interneurons and excitatory neurons^47,48^. Consistently, several anti-Hebbian LTP induction protocols failed to result in potentiation in visual cortex PV interneurons, instead leading to depression (Extended Data Fig. 15). This LTD was exaggerated in PV-Cre;lsl-eGFP-GluA2 mice compared to control mice (P = 0.03968, unpaired *t*-test), suggesting that CP-AMPARs regulate the expression of non-Hebbian LTD in PV interneurons.

Surprisingly, PV interneurons displayed drastically higher intrinsic excitability after CP-AMPAR removal, with a substantial increase in current-injected spike frequency, input resistance, and action potential (AP) half-width, along with a decrease in rheobase and afterhyperpolarization (AHP; Extended Data Fig. 16). The short AP half-width, low input resistance, and large AHP are all canonical features of PV interneurons^5,49^. This suggests that removing CP-AM-PARs led to a shift toward excitatory neuron-like intrinsic excitability characteristics. The resting membrane potential (RMP) was also higher in PV-Cre;lsl-eGFP-GluA2 mice when measured without synaptic glutamate and GABA receptor blockers (Extended Data Fig. 16j), suggesting a change in the balance of tonically active excitatory and inhibitory inputs (extrinsic synaptic excitability).

PV interneuron activation typically requires the coincident activation of multiple excitatory synaptic inputs^50,51^. However, the lower rheobase, higher RMP, and higher input resistance in PV-Cre;lsl-eGFP-GluA2 mice suggest some strong synapses may reach the activation threshold unilaterally, which can increase selectivity^52^. Together these results show intact connectivity but altered synaptic plasticity and intrinsic excitability in PV interneurons after removing CP-AMPARs, which may bias their recruitment to a few strong inputs, endowing them with increased tuning selectivity.

### Transcriptional response to CP-AMPAR removal

To investigate the novel link between CP-AMPARs and excitability, we assessed global PV interneuron transcriptome changes with FICSR-seq (Fixation-Capture Single Cell RNA Recovery-seq, see Methods) on forebrain PV interneurons (Extended Data Fig. 17). FACS-assisted PV interneuron bulk RNA-seq of PV-Cre;lsl-eGFP-GluA2 mice and PV-Cre;lsl-eGFP controls showed no expression changes in 278 out of 279 genes comprising the major classes of ion channels and excitatory/inhibitory synapse proteins (Extended Data Figs. Hong et al., 2023 (preprint) 18-20). This lack of expression changes suggests a post-transcriptional regulation of intrinsic and extrinsic (synaptic) excitability. The exception was GluA2 mRNA, which was expressed ∼2 fold compared to control PV interneurons (P_adj_ = 4.63×10^−9^, Benjamini-Hochberg correction; Extended Data Fig. 18), in agreement with protein measurements (Fig. 2b). GluA1, although downregulated at protein level (Fig. 2c), was unchanged at the mRNA level. These transcriptomic results suggest that the substantial changes in PV interneuron excitability after CP-AMPAR removal are not supported by changes in gene expression but likely reflect post-transcriptional regulation.

### CP-AMPARs blunt excitatory selectivity

We found that removing CP-AMPARs from PV interneurons renders them more selective. Conversely, we wondered whether introducing CP-AMPARs to excitatory neurons would reduce their orientation selectivity. To test this, we assessed visual representation in GluA2 homozygous knockout mice, where even excitatory neurons express abundant amounts of CP-AMPARs. Earlier studies established altered synaptic plasticity in these mice^53,54^, but the impact on sensory representation has not been reported.

Using a dual virus approach (Fig. 4a), we measured the visual responses of excitatory neurons (which typically have low CP-AMPAR levels) in GluA2 knockout mice (-/-) and littermate controls (+/+). Excitatory neurons in the GluA2 knockout mice displayed substantially lower orientation selectivity (Fig. 4b,c,e and Extended Data Fig. 21; P < 0.0001, Mann-Whitney U-test) and direction selectivity (Fig. 4d, e; P < 0.0001, Mann-Whitney U-test). The proportion of visually responsive neurons and average response amplitude were not significantly different (Extended Data Fig. 21b,c). Non-preferred stimuli responses are broadly reduced in excitatory neurons in the GluA2 knockout mice, leading to decreased orientation and direction selectivity (Fig. 4e and Extended Data Fig. 21d,e). These results suggest that CP-AM-PAR expression is sufficient to reduce selectivity regardless of neuron type.

**Figure 4.**
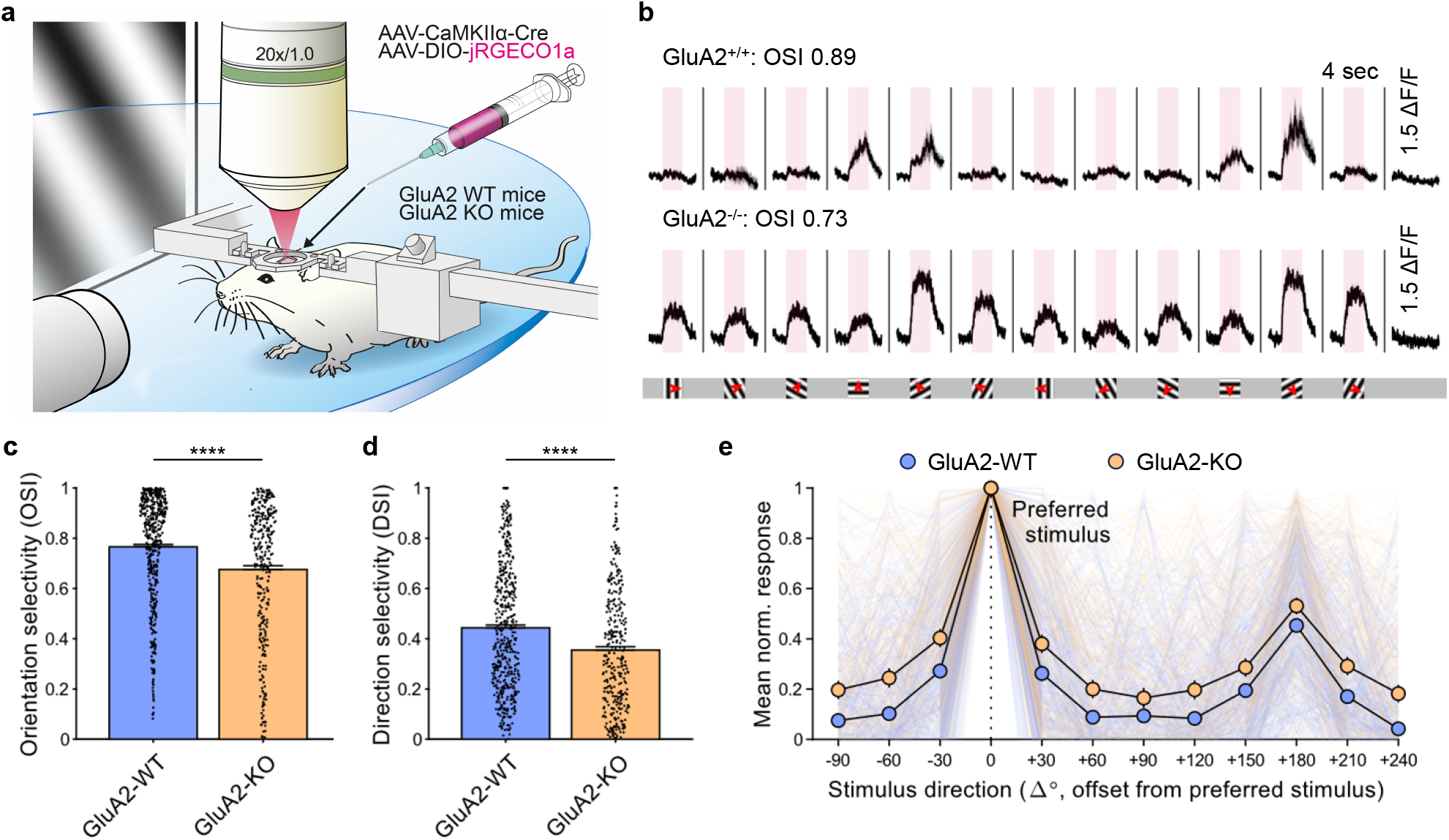
GluA2 homozygous knockout leads to decrease of selectivity in excitatory neurons. **a**, Pre-injected GluA2 knockout (KO; -/-) and littermate wild-type (WT; +/+) mice were head-fixed and visually stimulated during 2P imaging of V1 to reveal differences in tuning. **b**, Representative traces of GluA2-WT and GluA2 KO excitatory neurons. Pink regions denote the 4s visual stimulation period and 1.5 ΔF/F for both groups. Whole screen drifting grating stimulation with 12 different orientations was used to assess orientation selectivity. Red arrows mark the drifting direction. **c**, Quantification of orientation selectivity shows a significantly lower OSI in GluA2 knockouts compared to littermate wildtype (WT) controls (n = 504/340 from 3/3 mice, P < 0.0001, Mann-Whitney U-test). **d**, GluA2-KO group displays lower direction selectivity as well (P < 0.0001, Mann-Whitney U-test). **e**, Normalized average response profile of all positively responding neurons, aligned to their preferred stimulation (0°). Responses are plotted as mean ± SEM.

### Spatial selectivity of CA1 PV interneurons

We asked whether CP-AMPARs regulated PV interneuron selectivity beyond the visual cortex. PV interneurons in the hippocampus display lower spatial selectivity than their neighboring pyramidal cells^3,5^, but the molecular mechanisms underlying this lower selectivity are unknown.

Using a virtual-navigation task in head-fixed animals^55^ (Fig. 5a and Methods), we imaged hundreds of PV interneurons in the CA1 of PV-Cre mice transfected with Cre-dependent AAV expressing SEP-GluA2 or eGFP as a control (Fig. 5b) while mice were running on a 4-m long virtual linear track. We previously measured reliable place fields of excitatory neurons with this experimental setup^55^, indicating that the hippocampus forms a robust internal representation of the virtual environment.

**Figure 5.**
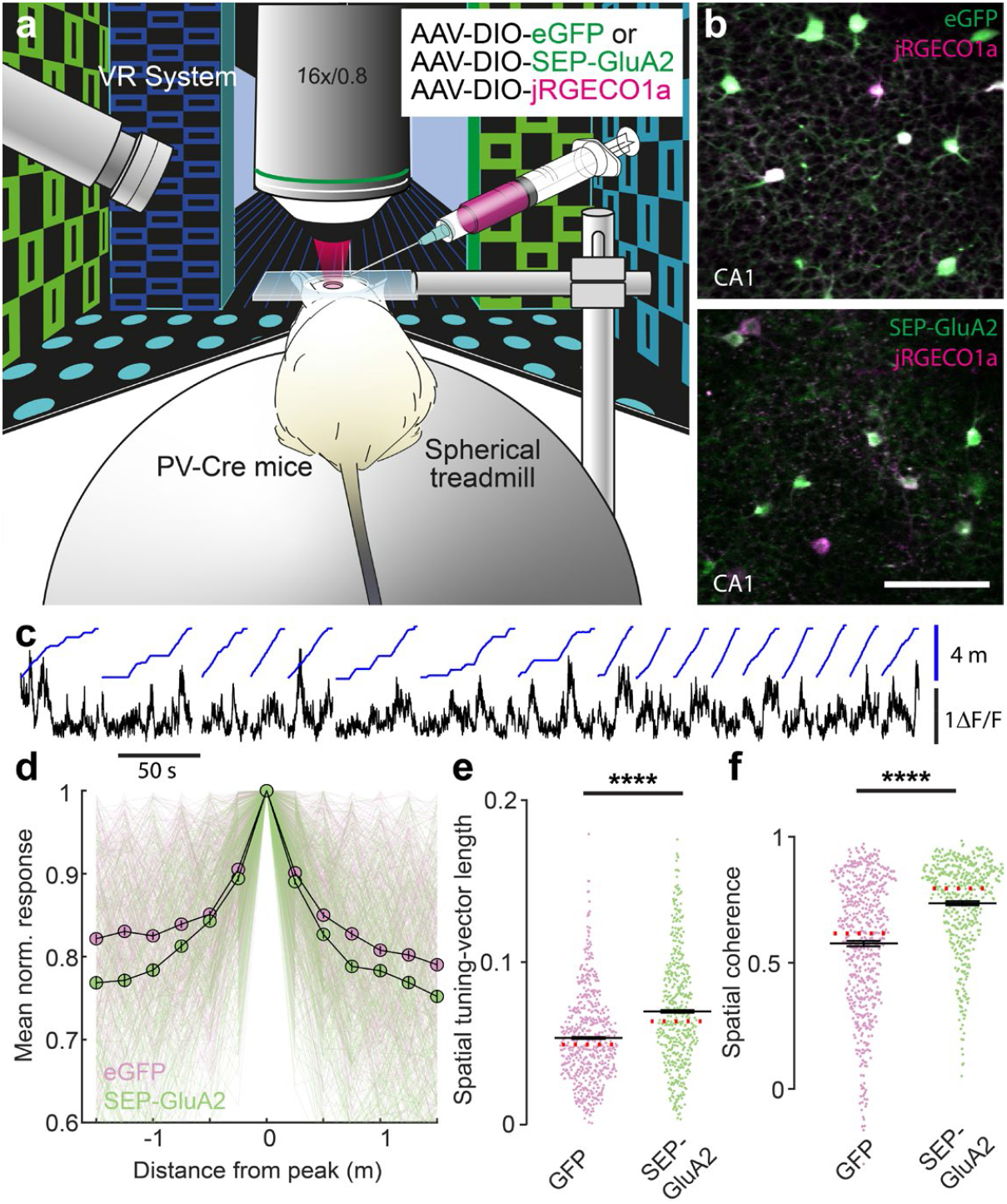
Increased spatial tuning of hippocampal PV interneurons after expression of GluA2. **a**, Experimental schematic of the virtual reality (VR) system. **b**, Time average of fluorescence acquired in vivo for jRGECO1a (magenta) and SEP-GluA2 (*left*) or eGFP (*right*) in green, respectively. Scale bar, 100 μm. **c**, Ca^2+^ activity traces (*black*) and mouse position in virtual reality linear track (*blue*) over time. **d**, Normalized average spatial response profile of hippocampal CA1 PV interneurons expressing SEP-GluA2 (green) or eGFP (magenta) aligned to the location of their peak activation. Responses are plotted as mean ± SEM. Thin lines denote individual cells. **e**, Spatial tuning-vector length (see also Extended Data Fig. 22c) of PV interneurons transfected with SEP-GluA2 was significantly higher than GFP controls (n = 583/476 cells from n = 4/4 mice, P = 1.472×10^−14^, Wilcoxon rank-sum test). **f**, Spatial coherence was also higher in the SEP-GluA2 group (P = 1.532×10^−26^, Wilcoxon rank-sum test). Black lines in (**e-f**) denote mean ± SEM, and the red dotted line denotes the median. Dots denote values for individual cells.

Strikingly, we observed that the activity of SEP-GluA2-expressing PV interneurons (Fig. 5c) was more sharply tuned than GFP-expressing PV neurons to a preferred location (Fig. 5d). This was reflected in higher spatial tuning-vector length (Fig. 5e and Extended Data Fig. 22a, b), higher spatial coherence (Fig. 5f; local smoothness of the spatial tuning curve), higher spatial information (Extended Data Fig. 22e), larger within-session stability of spatial tuning curves, and lower variability of spatial responses between trials (Extended Data Fig. 22b,f,e). In summary, these data suggest that GluA2-lacking CP-AMPARs lower the selectivity of PV interneurons regardless of modality and play a broad role in sensory representation beyond the neocortex.

## Discussion

Our results show that CP-AMPARs are both necessary for low orientation selectivity in PV interneurons and sufficient to induce lower selectivity in excitatory neurons (which typically have few CP-AMPARs). Postsynaptic AMPA receptors are well-understood for their role in synaptic transmission and defining synaptic strength^9,10,56^. These new results suggest that their biophysical properties can control neuronal response tuning, expanding their active role in computation. Whereas CP-AMPARs have been studied extensively in excitatory neuron synapses, our results attribute a novel role for CP-AMPARs in sensory representation to the forebrain CP-AMPARs, which overwhelmingly reside in inhibitory neurons.

Our findings have broad implications for understanding inhibitory architecture. Inhibitory PV interneurons provide rapid feedback inhibition to local excitatory neurons. This lateral inhibition constrains the timing and extent of their firing while reducing informational redundancy^5,12^. While the selectivity of PV interneuron activity compared with excitatory neurons^57-59^ and the tuning bias of their outputs on local excitatory neurons^41,44,45,60-62^ have been under debate, we show here that the lower selectivity of these PV interneurons is biophysically implemented with a well-conserved molecular mechanism, including transcriptional *Gria2* downregulation.

Whether other mammalian and non-mammalian organisms share such molecular or computational architecture is a question for future studies. It is fascinating to note that even ‘PV-like’ GABAergic neurons in evolutionarily distant lizards, which lack *Pvalb* expression, also display low *Gria2* expression^63^. Conversely, the importance of high calcium-impermeable AMPAR expression in excitatory neurons has been highlighted recently by the discovery of human heterozygous *de novo GRIA2* mutations through wholegenome sequencing efforts^64^. Mutations that lead to *GRIA2* loss of function or which remove the calcium-blocking pore residue of *GRIA2* are invariably associated with intellectual disability and autistic behaviors, suggesting that the tight control of AMPAR calcium permeability is essential for human cognition.

In a given brain area, neurons display varying levels of selectivity to their preferred stimuli set. This selectivity can be stratified along the line of neuronal cell types^12,16,65^. This stratification suggests gene expression can significantly impact a given neuron’s selectivity and sensory representation. Because synaptic inputs are summed in space and time to trigger neuronal activity, the selectivity of a neuron can be dictated by (a) the functional bias of the synaptic input pool^41^, (b) the organization of the input synapses along the dendritic structure, and (c) the intrinsic excitability of each neuron.

Previous studies suggested that PV have low selectivity because they receive high-density excitatory input from cells with diverse tuning features and low overall functional bias^20,43-45^. Our paired recordings show that gross input connectivity rates are intact in mice when PV neuron CP-AMPAR levels are lowered, demonstrating that PV interneuron orientation selectivity can increase without significantly changing connection rates. However, these recordings do not address whether the functional bias of input connections or the clustering of such synapses throughout the dendritic tree is altered by the lack of CP-AMPARs. Thus, anti-Hebbian plasticity may have a role in the dendritic organization of functionally tuned synapses. Meanwhile, our results show that the intrinsic excitability of PV interneurons is tightly coupled to the AMPA receptor profile, suggesting interleaved and coordinated mechanisms that define the computation of a given neuron. It is possible that blocking a key Ca^2+^ input source in dendrites by removing CP-AM-PARs leads to a homeostatic response in PV interneurons to upregulate excitability^66^. Intrinsic excitability is uniquely adapted in PV interneurons^5^, and whether this tightly regulated feature of PV interneurons is causally involved in selectivity remains an essential question for future studies.

What do these results tell us about biological and *in silico* intelligent circuits? Hebbian plasticity in neuronal networks is predicted to increase the correlation between neuronal activity and degrade total information content^1^. Anti-Hebbian plasticity is a possible mechanism to counteract this, reducing redundancy and keeping the representation more independent. However, researchers have traditionally thought Anti-Hebbian plasticity was implemented at the output GABAergic synapses of the inhibitory network^1,67^. By contrast, our work shows that CP-AMPARs at the *input* of the inhibitory network lower GABAergic selectivity, allowing PV interneurons to broadly inhibit correlated activity through lateral inhibition. This characteristic adds to the flexibility of the network and may contribute to the canonical normalization computation that PV interneurons are thought to carry out^68^.

We have described a mechanism that commonly governs PV interneuron selectivity across multiple modalities, from orientation/direction selectivity in the visual cortex to spatial selectivity in the hippocampus. By no means have we exhaustively assessed the selectivity of these neurons in other domains, such as color, ocular dominance, and speed^4,15,19,21,41^. Selectivity to some visual features emerges before visual experience at eye-opening and can be driven by genetically-determined circuit formation^15^. Future studies will show how experience-dependent synaptic regulation through CP-AMPARs interacts with genetically-determined mechanisms to fine-tune sensory representation. As we look beyond PV interneurons, we anticipate the discovery of known and yet-to-be-discovered mechanisms in other cell types that interact together, enabling internal representations to support intelligent behavior.

## Methods

### Mice and marmosets

All procedures were approved by the Johns Hopkins Animal Care and Use Committee and conducted per the guidelines of the National Institutes of Health and the Society for Neuroscience. Hippocampal imaging experiments were carried out according to German national and institutional guidelines and approved by the ‘Tierversuchskommission’ of the Regierungspräsidium Freiburg (license no. G16/037). Marmoset post-mortem tissue was obtained from terminal experiments approved by NIH Institutional Animal Care and Use Committees. We used these mouse lines: PV-Cre^38^ (JAX #008069), lsl-eGFP^69^ (JAX #010701), lsl-eGFP-GluA2 (Extended Data Fig. 6), GluA2 KO^53^ (JAX #002913), GluA1 KO^70^ (JAX #024422). We generated the ROSA26-lsl-eGFP-GluA2 mouse line by electroporating mouse ES cells with an engineered construct containing ROSA26-CAG-loxP-STOPloxP-eGFP-Gria2-WPRE (adapted from targeting vector used to generate Ai14 mice^71^) and homologous recombination (Extended Data Fig. 6). We acquired PV-Cre;lsl-eGFP-GluA2 (and PV-Cre;lsl-eGFP) mice from a cross with PV-Cre mice, born at Mendelian ratios. GluA2^-/-^ pups displayed lower body weight compared with wild-type littermates. They displayed occasional mortality, mitigated by separating the littermates from the parents to reduce litter sizes^53^. All lines were maintained on a mixed background composed primarily of C57BL/6J, and mice of both sexes were used for experiments. We maintained all animals on a 12 hr light/dark cycle.

### Constructs

We used Q/R and R/G RNA-edited flip-isoform short c-tail rat GluA2 cDNA sequences for mutant animal generation and viruses unless otherwise stated. SEP-GluA2 and GFP-GluA2 fusion constructs were generated by N-terminal insertion of SEP or GFP at four amino acids after the signal peptide padded with linker sequences, as in previously published constructs^72^. We generated the FUW-Cre construct by replacing the eGFP in FUGW with the Cre recombinase gene.

pAAV.Syn.Flex.NES-jRGECO1a.WPRE.SV40^73^ was a gift from Douglas Kim & GENIE Project (Addgene plasmid #100853). The loxP/lox2272 sequences in the Flex cassette were inverted or exchanged with lox511/loxFAS to mitigate compatibility with other DIO AAVs. pAAV-CW3SL-EGFP^74^ was a gift from Bong-Kiun Kaang (Addgene plasmid #61463).

To deliver large genes, such as the SEP-GluA2 fusion gene with the high tropism and low cytotoxicity provided by adeno-associated virus (AAV) vectors, we heavily optimized vector components to allow larger transgene size. Using the short hSyn1 promoter (469bp), abbreviated linker sequences and DIO sequences, and an optimized WPRE+polyA signal (CW3SL, 425bp)^74^, we generated a pan-neuronal Cre-dependent AAV expression vector with a minimal backbone (1350bp ITR to ITR without cargo) and large cargo capacity size (∼3.65kb; based on an earlier estimation of 5kb AAV genome size limit^75^; 3.85kb where Cre-dependency is not required). The loxP/lox2272 sites were spaced by a minimal 64bp (5’ end-to-5’ end) to set the second recombination event distance (128bp) above 118bp, at which inefficient recombination has been reported, but at an exact multiple of the helical repeat length (10.6 bp). This repeat length allowed better-aligned loxP sites upon DNA looping, thereby maximizing the efficiency of Cre-mediated excision^76^.

As proof-of-principle, this study showed that SEP-GluA2 (3378bp), a large fusion protein previously only expressed through electroporation or lentiviral transfection, can be robustly expressed with this vector both *in vitro* and *in vivo* (Extended Data Fig. 12). The DIO-SEP-GluA2Q vector harbored a GluA2 cDNA unedited at the Q/R editing site (R607Q)^77^. GluA2 Q/R RNA editing occurs at the pre-mRNA stage and requires a hairpin structure in the adjacent intron, which is absent in this vector. This structure bypasses RNA editing and expression of a calcium-permeable GluA2Q subunit. The DIO-eGFP control virus was similarly generated, replacing SEP-GluA2 with eGFP, for use as a control.

AAV was produced by HHMI-Janelia Viral Tools using a PEI triple transfection protocol into AAV293T cells (an ITR-containing plasmid, 2/9 capsid helper from UPenn Vector Core, and the E1-deleted pHelper plasmid from Agilent). The cells were grown under serum-free conditions (three 150mm culture dishes at ∼3×10^7^ cells/dish for each 100 µl batch), purified by two rounds of CsCl density gradient centrifugation, and exchanged into storage buffer (1xPBS, 5% Sorbitol, 350mM NaCl). Virus titers (GC/ml) were determined by qPCR targeting the AAV ITRs.

### Stereotaxic cranial surgeries

We used stereotaxic surgery to inject viruses and implant 4 mm square cranial windows over the left primary visual cortex (V1). Mice of mixed sex (>6-week-old) were given Carprofen (5mg/kg) or Buprenorphine (sustained release; 0.5-1.0 mg/kg) and Dexamethasone (4mg/kg) for analgesia and were anesthetized using Avertin or isoflurane (1.5-2.5%). We made a craniotomy with a #11 scalpel blade centered at 2.5 mm lateral and 3.4 mm posterior to bregma.

For AAV injections, viruses were diluted with sterile phosphate-buffered saline (PBS) to 1∼5×10^13^ GC/ml. We injected the solution at 5-10 sites spanning the posterior central area of the craniotomy (corresponding to V1), with ∼100 nl injections at each site at 250 μm below the dura surface. Injections were made using a beveled glass pipette and a custom mineral oil-based injection system over 2–4 min. We left the pipette in place for another 2–3 min to allow diffusion and prevent backflow.

We placed a 4 mm square glass coverslip over the craniotomy and attached a stainless-steel head bar to the skull during surgery to allow rigid head-fixation during imaging. We allowed mice to recover for 1-2 weeks before imaging and handled them extensively to alleviate experiment-related stress.

For hippocampal experiments, virus injections and cortical excavation/window implantation were done in separate surgeries. We made a small craniotomy over the hippocampus and injected 500 nl of AAV into CA1 (A/P: -2.0 mm; M/L 2.0 mm; D/V -1.4 mm). In the same surgical session, we implanted mice with a stainless-steel head plate (25 × 10 × 0.8 mm with an 8 mm central aperture) horizontally. We allowed mice to recover from surgery for at least 5 days before training sessions. We continued postoperative analgesic treatment with Carprofen (5 mg/kg body weight) for 3 days after surgery.

Cortical excavation and hippocampal imaging window implantation were performed >10 days after the initial virus injection, per published protocols^55^. We made a craniotomy (diameter 3 mm) centered at A/P -1.5 mm and M/L -1.5 mm. Parts of the somatosensory cortex and posterior parietal association cortex were gently aspirated while irrigating with chilled saline. We continued aspiration until the external capsule was exposed. We then gently peeled away the outer part of the external capsule using fine forceps, leaving the inner capsule and the hippocampus undamaged. The imaging window implant consisted of a 3 mm diameter coverslip (CS-3R, Warner Instruments) glued to the bottom of a stainless-steel cannula (3 mm diameter 1.2-1.5 mm height). The window was gradually lowered into the craniotomy using forceps until the glass was in contact with the external capsule. The implant was then affixed to the skull using cyanoacrylate. We allowed mice to recover from window implantation for 2-3 days.

### Awake in vivo 2-photon fluorescence imaging

We performed retinotopic mapping^78,79^ to verify the location of V1 using optimized protocols and software (https://github.com/ingiehong/retinotopy). We conducted awake in vivo two-photon imaging with a custom-built, resonant/galvo two-photon laser-scanning microscope (Sutter Instrument) controlled by ScanImage (Vidrio Technologies) and light-proofed to allow imaging in ambient light during visual stimulation. The designs for the head-fixed imaging platform and lightproofing apparatus are available online (https://github.com/ingiehong/StackGPS). We imaged neurons in L2/3 of monocular V1 expressing eGFP/SEP and jRGECO1a using a 20×/1.0 NA water-immersion objective (Zeiss) and a Ti:Sapphire laser (Coherent Chameleon Ultra; Spectra-Physics Insight X3) tuned at 930 nm or 1040 nm, respectively, with 20∼100 mW of power delivered to the back-aperture of the objective.

We corrected the lateral motion of acquired image sequences using a rigid motion correction algorithm (NoRMCorre^80^). Neuronal somata with calcium transients were segmented using a constrained non-negative matrix factorization (CNMF) algorithm^81^. The source-separated GCaMP/jRGECO1a signal from each neuron was used to estimate various visual response properties of L2/3 neurons.

### Visual stimulation

Visual stimuli were presented on a gamma-corrected 27” LED monitor placed 22 cm in front of the center of the eye contralateral to the hemisphere in which imaging was performed. The visual stimuli consisted of full-screen drifting gratings (4 sec duration, sinusoidal, 0.05 cycles/deg, 1 Hz, 100% contrast) following a 4 sec iso-luminant grey screen. 6 orientation gratings spaced at 30° were presented drifting in both directions orthogonal to the gratings (total 12 directions) in a pseudo-randomized order to characterize sensory tuning, using Psychtoolbox-3^82^ and FocusStack/Stimserver^83^. We used the average response during the 4 sec stimuli across 9-11 presentations to calculate visual responsiveness and orientation/direction selectivity. Visually responsive neurons were defined as cells with significant stimulus-related fluorescence changes (ANOVA across blank and twelve direction periods, P < 0.05)^84^.

The orientation/direction tuning curve was constructed by measuring the mean ΔF/F, averaged over the stimulus period for each grating drifting direction θ, denoted as *R*(θ). The orientation selectivity index (OSI) was calculated for visually responsive units^21,84,85^ with slight modifications on prior definitions^85^ to avoid values outside the intended interval ([0 1]) and to accommodate occasional *bona fide* negative responses^86-88^. The preferred drifting direction (θ*oppo*) of the cell was determined as the stimuli that evoked the greatest responses, *R*(θ*pref*) and *RR*(θ_*oppo*_), as a sum where θ_*oppo*_ = θ_*oppop*+180°_, *R*(θ_*pref*_) > *R*(θ_*oppo*_). The orientation selectivity index (OSI) was defined as:

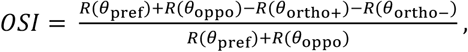

All response values were subtracted by the most negative *R*(θ) when negative responses were present (*R*_*corrected*_), effectively ensuring the relative dynamic range of responses were reflected in the index where they would otherwise distort the index (leading to values outside [0 1]), or be clipped (when negative values were discarded). Formally,

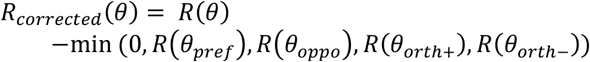

Empirically, this modified index correlates tightly with the OSI calculated with the prior definition^85^ of OI/OSI, is bounded by [0 1], and accommodates tuning curves that are partially or entirely negative. Notably, the trends and results of statistical comparisons in this work did not change with the choice of index definition. Direction selectivity (DSI), global orientation-selectivity index (gOSI), and global direction selectivity (gDSI) were defined as:

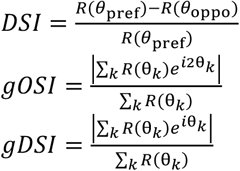

gOSI and gDSI gave the same conclusions as OSI/DSI (data not shown). Note that *R*_*corrected*_(θ) can also be used in gOSI and gDSI, with the same benefits.

### Head-fixed navigation and hippocampal imaging

Mice implanted with hippocampal imaging windows were subjected to a custom head-fixed virtual reality environment described earlier^55^. It consisted of a spherical treadmill monitored by an optical sensor that translated motion on the treadmill into forward motion through the virtual environment. We adjusted the forward gain so that 4 m of distance traveled along the circumference of the treadmill equaled one full traversal along a simulated linear track displayed on monitors surrounding the mouse. The track consisted of textured walls, floors, and other 3D-rendered objects at the track’s sides as visual cues. To motivate consistent behavior, we administered soy-milk rewards (4 µl) when the animal traversed certain locations that were spread at fixed distances along the track, and animals were trained for 5-10 days until they displayed consistent running behavior before commencing imaging experiments.

Imaging was performed using a resonant/galvo highspeed laser scanning two-photon microscope (Neurolabware) with a frame rate of 30 Hz for bidirectional scanning and a power of 5-20 mW measured at the objective front aperture. The microscope had an electrically tunable, fast z-focusing lens (Optotune, Edmund optics) to switch between z-planes within less than a millisecond. Images were acquired through a 16x objective (Nikon, 0.8 N.A., 3 mm WD). eGFP and jRGECO1a were excited at 930 nm or 1040 nm, respectively, with a femtosecond-pulsed two-photon laser (Mai Tai DeepSee®, Spectra-Physics). We scanned three imaging planes (∼25 µm z-spacing between planes) in rapid alternation so that each plane was sampled at 10 Hz. The planes spanned 300-500 µm in x/y direction and were placed so as many labeled neurons as possible were depicted. We attached the animal’s head plate to the bottom of an opaque imaging chamber before each experiment to block ambient light from the photodetectors. We fixed the chamber in the behavioral apparatus with the animal. A ring of black foam rubber between the imaging chamber and the microscope objective blocked any remaining stray light.

### Spatial tuning analysis

We motion-corrected all imaging data line-by-line using the SIMA software package^89^ with a 2D Hidden Markov Model or the software package ‘Suite2P’^90^. If no decent motion correction could be achieved, we discarded the data. To segment interneuron somata, regions of interest (ROIs) were drawn manually using ImageJ (NIH) or automatically by applying the ‘Suite2P’ software package^90^. In the case of automated ROI settings, the experimenter subsequently inspected individual ROIs. The average jRGECO1a signal over time was then obtained from each ROI for all runs. We restricted our analysis to mouse running periods with a minimum speed of 5 cm*s^-1^. To obtain baseline-normalized ΔF/F calcium traces, we examined the fluorescence value distribution of the jRGECO1a signal and subtracted and divided the entire trace by the 8th percentile value of this distribution^91^. Rarely, individual datapoints ended up below zero after baseline subtraction, and we set these negative values to zero for further calculations.

To compute spatial vector tuning, we plotted the mean activity (ΔF/F) of each spatial bin at its respective angle from the start position on the circular track into a polar coordinate system (Fig. 5e and Extended Data Fig. 22c). We then computed the circular mean of this distribution to obtain the cell’s mean tuning vector length and angle. Spatial coherence (Fig. 5f) was determined as the correlation (Pearson’s R) between the mean fluorescence value in each 5 cm bin on the track and its two nearest neighbors, measuring the local smoothness of the spatial tuning curve^92^. To calculate spatial information (SI; Extended Data Fig. 22e), we computed the average calcium activity (mean ΔF/F) for each 5 cm wide bin along the linear track to approximate the neurons’ average firing rate in that location. SI was then calculated for each cell as 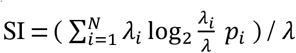, where *λ*_*i*_ and *p*_*i*_ are the average calcium activity and fraction of time spent in the i-th bin, respectively, *λ* is the overall calcium activity averaged over the entire linear track, and N is the number of bins on the track. Given the distribution of the underlying values, we plotted the log10 of SI values and compared them statistically (Extended Data Fig. 22e).

To assess the stability of a cell’s spatial representation within a session, we divided the track into 5 cm bins and calculated the mean ΔF/F value for each bin while the animal was moving on the track with a speed > 5 cm/s to get activity maps for each individual cell. This mapping was done separately for the first- and second half of the recording session. We then computed the within-session stability as the cross-correlation between the mean activity maps of the first and second half of the session (Extended Data Fig. 22b,f). We also computed population vector correlations as a function of position in the first and second half of the recording (Extended Data Fig. 22g) to visualize the local similarity of population activity across time. Before computing these correlations, we re-normalized each neuron’s map by subtracting the mean over space and dividing by the standard deviation (z-scoring) to mitigate the potential effects of mean rate differences between cells on assessing local population vector similarity.

### Quantification of Gria2 mRNA A-to-I editing rates

We mapped the raw sequencing reads from a mouse brain single-cell RNA-seq dataset (n = 1679)^14^ to the mouse reference genome (GRCm38) with a gene annotation, GENCODE vM16^93^ using STAR^94^. The uniquely-mapped reads whose sequencing qualities (Phred score) were greater than 20 were counted for the “QR” and “RG” RNA editing sites in Gria2. We filtered out samples if the proportions of the sequencing read with “A” or “G” alleles together accounted for less than 95%, to avoid potential sequencing errors. We defined the RNA editing rate for a given site as a ratio of the number of sequencing reads showing “G” relative to the number of reads with either “A” or “G.”

### FACS-assisted RNA-seq of PV interneurons

To assess transcriptional changes specifically in PV interneurons after removing CP-AMPARs with RNA-seq, we sorted dissociated cortical PV interneurons by their GFP fluorescence with FACS. Dissociation of adult mouse brain neurons leads to a rapid decimation of viable PV interneurons^95-97^, potentially biasing downstream analyses to a select subpopulation of PV interneurons. Various proposed methods to mitigate PV interneuron loss failed to recover them at native cell frequencies in adult mice^98^. Several fixation-based FACS approaches have been proposed to target immune cells and neurons, but crosslinking leads to lower RNA yield for RNA-seq.

We developed and used a brain-slice optimized workflow, fixation-capture single-cell RNA recovery-seq (FICSRseq), which recovers PV interneurons vulnerable to dissociation at native cell frequencies. We cut brain slices from adult mice (113.1 ± 11.6 days old) in NMDG cutting solution + Trehalose^95^ and diced them into small pieces <1 mm^3^. Extracellular proteins were digested with pronase (2 mg/ml; 8 U/µl) at 34-37°C, after which the slice pieces were fixed in 4% PFA in PBS (with 0.1 U/ml RNase inhibitor, Promega) for 15 mins and dissociated into single cells through careful trituration. We filtered the single cells through a 40 μm filter, labeled them with the cell-permeable nuclear dye DRAQ5 (1:1,000 dilution) to identify nuclei-containing cells, and then subjected them to FACS. DRAQ5+/GFP+ or DRAQ5+/GFP-cells were sorted, and over 20K cells were collected per mouse cortex to provide extensive coverage of low-expressing PV interneuron transcripts.

We treated the fixed cells with Proteinase K before RNA extraction (RecoverAll Total Nucleic Acid Isolation Kit for FFPE, Thermo Fisher Scientific) to liberate RNA from protein-protein and protein-nucleic acid crosslinks generated by fixation. We prepared cDNA libraries from GFP+ and GFP-samples (NEBNext Ultra RNA Library Prep Kit for Illumina, NEB) from RNA enriched with mRNA through bead-based polyA selection. cDNA libraries were barcoded and sequenced together on an Illumina Hiseq 2500 sequencer, generating 150-bp paired-end reads. We processed RNA-seq reads with bcbio-nextgen v1.2.3 (https://doi.org/10.5281/zenodo.3564938), aligning to GRCm38 with the STAR aligner^94^ and quantifying counts per gene with Sailfish^99^ using the Ensembl annotation. We used DESeq2^100^ to analyze differential expression.

### Brain slice preparation and whole-cell patchclamp recordings

We anesthetized mice of either sex (P32-P62 for studies of synaptic properties, P69-P77 for studies of intrinsic properties) using isoflurane. We rapidly removed their brains in an ice-cold sucrose solution containing the following (in mM): 76 NaCl, 25 NaHCO_3_, 25 glucose, 75 sucrose, 2.5 KCl, 1.25 NaH_2_PO_4_, 0.5 CaCl_2_, 7 MgSO_4_, pH 7.3, 315 mOsm. We hemisected the brain along the midline and mounted one or both hemispheres on a 30° ramp. We then sectioned acute parasagittal slices of the visual cortex, 300 μm thick, in the same ice-cold sucrose-cutting solution using a vibratome (VT1200s, Leica). Slices were incubated in warm (32–35°C) sucrose solution for 30 min and then transferred to warm (32–35°C) artificial cerebrospinal fluid (aCSF) composed of the following (in mM): 125 NaCl, 26 NaHCO_3_, 2.5 KCl, 1.25 NaH_2_PO_4_, 1 MgSO_4_, 20 D-(+)-glucose, 2 CaCl_2_, 0.4 ascorbic acid, 2 pyruvic acid, 4 L-lactic acid, pH 7.3, 315 mOsm. Slices were then allowed to cool to room temperature. For rectification measurements, we cut coronal slices with an NMDG-based cutting solution and incubated them >15 mins. Then we transferred them to aCSF (see ‘Analysis of AMPAR rectification’ section). All solutions were continuously equilibrated with 95% O_2_/5% CO_2_.

We transferred slices to a submersion chamber on an upright microscope (Zeiss AxioExaminer; 40X objective, 1.0 N.A.) and continuously superfused (2-4 ml/min) them with warm (∼32-34°C), oxygenated aCSF. We visualized neurons with a CCD camera (Sensicam QE, Cooke) using either infrared differential interference contrast (IR-DIC) microscopy or epifluorescence. The visual cortex was identified based on the relative position of the cortex and hippocampus and the anatomical borderline between the visual cortex and retrosplenial dysgranular cortex (RSD). We selected slices in which the apical dendrites of infragranular pyramidal neurons ran parallel to the plane of the slice up through L2/3 in the area targeted for recording. PV interneurons were targeted for recording based on eGFP or SEP-GluA2 expression along with unlabeled L2/3 pyramidal neurons. We filled patch pipettes (2-4 MΩ) pulled (P-97, Sutter Instrument) from borosilicate capillary glass (Sutter Instrument) with an internal solution containing (in mM): 2.7 KCl, 120 KMeSO_3_, 9 HEPES, 0.18 EGTA, 4 ATP magnesium salt, 0.3 GTP sodium salt, 20 phosphocreatine disodium salt, and adjusted to pH 7.3, 295 mOsm. For recordings of PV interneurons, the internal solution included 0.25% w/v biocytin. Whole-cell patch-clamp recordings were obtained using Multiclamp 700B amplifiers (Molecular Devices) and digitized using an Instrutech ITC-18 (HEKA) and software written in Igor Pro (Wavemetrics). All signals were low-pass filtered at 10 kHz and sampled at 20-100 kHz. Neurons with an access resistance >30 MΩ or a resting membrane potential greater than -60 mV were not used for further recordings or analysis. The access resistance was not compensated in current clamp, and recordings were not corrected for the liquid junction potential.

### Analysis of intrinsic excitability, synaptic connectivity, and synaptic plasticity

We measured the resting membrane potential (RMP) shortly after establishing the whole-cell current-clamp recording configuration. A 1second hyperpolarizing current (−100 pA) pulse was used to calculate the input resistance of recorded neurons. To assess the spiking behavior of the cell, we injected 1-second depolarizing current steps into the recorded neurons. We measured the current–spike frequency relationship with a range of depolarizing current steps presented in pseudorandom order (1-s long, 40-pA increments, 5-s interstimulus intervals). Each current intensity was tested three times. For each current intensity, we counted the total number of action potentials exceeding an amplitude of 0 mV generated during each current step, then averaged the values across the three trials. We determined the rheobase by first probing the neuron’s response with 1-s-long depolarizing steps (5-s interstimulus intervals) to define a small range of current steps that bounded the rheobase. We then tested the neuron response within this range using 1-s-long depolarizing steps with 1-pA increments. We measured action potential properties from single spikes evoked by rheobase current injections. To compare the current–spike frequency relationship and rheobase between cells from the same baseline, we held cell membrane potentials at -70 mV when injecting depolarizing current steps. We performed all electrophysiological recordings assessing the intrinsic properties of PV interneurons in the presence of the following blockers of glutamate and GABA receptors: 5 µM NBQX (AMPA receptor antagonist), 5 µM (RS)-3-(2carboxypiperazin-4-yl)-propyl-1-phosphonic acid (NMDA receptor antagonist), and 10 µM 6-imino-3-(4-methoxyphenyl)-1(6H)-pyridazinebutanoic acid hydrobromide (SR95531;GABA_A_ receptor antagonist; all from Tocris Bioscience).

To determine the properties of unitary synaptic connections among neurons, we generated two action potentials in the presynaptic neuron by injecting short, depolarizing current steps (3-ms pulse duration, 20 Hz, 10-s inter-trial interval). We held pyramidal neurons and PV interneurons at approximately -55 mV and -70 mV during synaptic connectivity tests to detect IPSPs and EPSPs, respectively. We assessed synaptic connectivity (EPSP or IPSP) with an average of 10-50 trials. A synaptic connection was detected if the average trace’s first response amplitude was >3 times the root mean squared (RMS) of the average trace during baseline conditions and verified visually. We calculated the paired-pulse ratio (PPR) by dividing the amplitude of the second postsynaptic potential by the first.

We subjected a subset of connected Pyr→PV pairs, all of which exhibited an average EPSP amplitude > 0.3 mV at baseline, to an anti-Hebbian (AH) protocol. After recording 50 traces (6 Hz) as a baseline, we induced synaptic plasticity by pairing 400 presynaptic action potentials delivered at 5 Hz with continuous hyperpolarization of the postsynaptic PV interneuron to –90 mV ^30,101^. After induction, EPSPs were recorded under the same conditions as the baseline measurement (50 traces in response to presynaptic action potentials, 6 Hz).

### Analysis of AMPAR rectification

To measure AMPAR rectification^102,103^, we cut coronal brain slices in ice-cold cutting solution containing (in mM) 96 NMDG, 2.5 KCl, 1.25 NaH_2_PO_4_, 25 NaHCO_3_, 25 D-(+)-glucose, 10 MgSO_4_, 0.5 CaCl_2_, 96 HCl, 20 HEPES, 12 N-acetylcysteine, 5 sodium L-ascorbate, and oxygenated with carbogen gas (95% O_2_ and 5% CO_2_). The 300 µm-thick slices were kept in aCSF (125 NaCl, 2.5 KCl, 2 MgCl_2_, 2 CaCl_2_, 1.0 NaH_2_PO_4_, 26.2 NaHCO_3,_ and 11 glucose, oxygenated with carbogen gas at 23–25 °C until they were transferred for recording to a submerged chamber superfused with aCSF (1-3 ml/min) supplemented with ∼50 µM picrotoxin and 100 μM APV to isolate AMPAR-mediated excitatory synaptic transmission.

We made targeted whole-cell recordings of eGFP/SEP-GluA2-positive L2/3 PV interneurons using pipettes of 3-5 M*Ω* resistance. The intracellular solution contained (in mM): 115 CsMeSO_4_, 0.4 EGTA, 5.0 TEA-Cl, 1 QX314, 2.8 NaCl, 20 HEPES, 3.0 ATP magnesium salt, 0.5 GTP sodium salt, 10 phosphocreatine disodium salt, 0.1 spermine and was adjusted to pH 7.2, 285–290 mOsm. When we achieved whole-cell mode, we allowed > 5 min for dialysis of the intracellular solution before collecting data. We held cells at -70 mV holding potential and recorded them at room temperature. We left the junction potential (∼11 mV) uncorrected. Signals were measured with a MultiClamp 700B amplifier, digitized using a Digidata 1440A digitizer (Molecular Devices) at 20 kHz, and acquired with pClamp 10 software (Molecular Devices). We recorded AMPAR currents at 11 membrane potentials to construct a current-voltage (I-V) plot (V_h_ = -60 to +60 mV, except for a subset of pyramidal neurons recorded for comparison up to +50 mV). We calculated the rectification index as a weighted ratio of negative (−60 mV) and positive (+60 mV) currents. We compensated for the junction potential (11mV): Rectification index (RI) = (I_-60mV_/-71)/(I_+60mV_/49). An AMPAR rectification index of 1 represented perfect linearity, whereas values <1 indicate inward rectification. We estimated the reversal potential (E_rev_) by cubic polynomial regression that fitted the linear, rectifying, and double-rectifying AMPAR I-V curves well.

### Immunohistochemistry

We deeply anesthetized mice with isoflurane, then transcardially perfused them with phosphate-buffered saline (PBS) and 4% paraformaldehyde (PFA). We removed and postfixed the brain in 4% PFA/PBS for >2 hours. We sectioned the brain coronally into 25 μm slices using a vibratome (VT-1000, Leica). We acquired marmoset brains post-mortem from terminal experiments and sliced them into 40 µm sections. Free-floating sections underwent antigen retrieval using L.A.B. solution (Polysciences) when necessary and were blocked and permeabilized in 3% BSA with 0.3% Triton X-100 in PBS for 1 h at room temperature. We incubated sections with primary antibodies overnight at 4°C, washed them with PBS three times for 5 mins, and then incubated them with secondary antibodies for 2 hours at room temperature. After another round of washes, we mounted the slices on glass slides in PermaFluor mounting medium (Thermo Fisher Scientific) and imaged them using a laser scanning confocal microscope (Zeiss LSM880).

The following primary antibodies were used: rabbit anti-parvalbumin (1:2000, PV25, Swant), goat anti-parvalbumin (1:1000, PVG-213, Swant), rat anti-somatostatin (1:200, MAB354, Chemicon), mouse anti-CaMKIIα (1:1000, sc-32288, Santa Cruz), rabbit anti-GluA1 (1:1000, JH4294, generated in-house), mouse anti-GluA2 (1:5000; clone 15F1, generous gift from E. Gouaux), chicken anti-GFP (1:1000, GFP-1020, Aves), and rabbit anti-dsRed2 (1:1000, Clontech). The following secondary antibodies were used: Alexa Fluor 405 donkey anti-goat (1:1000, ab175665, Abcam), Dylight 405 goat anti-mouse IgG2a (1:1000, 115-477-186 Jackson ImmunoResearch), Alexa Fluor 488 goat anti-mouse IgG2a (1:1000, A-21131, Thermo Fisher Scientific), Alexa Fluor 488 goat anti-chicken (1:1000, A-11039, Thermo Fisher Scientific), Alexa Fluor 546 goat anti-rabbit (1:1000, A-11035, Thermo Fisher Scientific), Alexa Fluor 568 goat anti-mouse IgG1 (1:1000, A-21124, Thermo Fisher Scientific), Alexa Fluor 568 goat anti-rabbit (1:500, Thermo Fisher Scientific), Texas Red donkey anti-goat (1:1000, SAB3700332, Millipore Sigma), Alexa Fluor 647 goat anti-rabbit (1:1000, A-21245, Thermo Fisher Scientific), Alexa Fluor 647 goat anti-mouse IgG2a (1:1000, A-21241, Thermo Fisher Scientific), Alexa Fluor 647 donkey anti-goat (1:1000, A-21447, Thermo Fisher Scientific), and Alexa Fluor 647 goat anti-rat (1:500, A-21247, Thermo Fisher Scientific).

### Statistical Analysis

We performed statistical tests in MATLAB (Mathworks), Prism (Graphpad), or R. Data distributions were tested for normality with the Shapiro–Wilk test. We used parametric tests if the data were normally distributed and non-parametric otherwise, as detailed in the text describing each comparison. For parametric tests, we used unpaired/paired t-tests and 1-way/2-way ANOVA tests with Tukey’s post-hoc multiple comparison correction. For data that did not follow normal or log-normal distributions, we used the following statistical tests where appropriate: Mann–Whitney U-test (Wilcoxon rank-sum test), Kruskal-Wallis 1-way ANOVA with Dunn’s post-hoc multiple comparison correction (all two-sided). For categorical data, we used Fisher’s test or χ2 with/without Yates correction according to degrees of freedom and sample size. We reported center and spread values as mean ± SEM (Standard Error of the Mean) or median ± IQR (interquartile range) unless otherwise stated. We used no statistical methods to plan sample sizes but used sample sizes similar to those frequently used in the field. The text or figure legends include the number of animals and cells. We did not use randomization; data collection and analysis were not performed blind to the conditions of the experiments unless otherwise stated. P-values < 0.05 were considered to be statistically significant. When we used a statistical test, the P-value is noted either in the manuscript text or depicted in figures and legends as: *P < 0.05, **P < 0.01, ***P < 0.001, ****P < 0.0001, n.s., not significant, P *≥* 0.05.

## Data Availability

Sequencing data included in this manuscript will be available at NCBI GSE, under the accession number GSEXXXXXX. Other data of this study is available from the corresponding author upon reasonable request.

We used the following previously published datasets:

Tasic B et al., 2016. Adult mouse cortical cell taxonomy by single cell transcriptomics. NCBI Gene Expression Omnibus. GSE71585

Lake B et al., 2016. Neuronal subtypes and diversity revealed by single-nucleus RNA sequencing of the human brain. dbGaP Study Accession phs000833.v3.p1

Krienen FM et al., 2020. Innovations in primate interneuron repertoire. NCBI Gene Expression Omnibus. GSE151761

## Code availability

Computer codes used to acquire data and analyze results of the study are available at https://github.com/ingiehong/retinotopy, https://github.com/ingiehong/StackGPS, and from the corresponding author upon reasonable request.

## Acknowledgments

We would like to thank Alexei Bygrave, Richard Roth, Elena Lopez-Ortega, Han Tan, Austin Graves, Ashley Irving, Sarah Richardson, Michele L. Pucak, Michael Ayad, Sam Das, Joshua Kufera, and Johnson Moran for technical assistance and scientific discussions. We thank Lisa Hamm for administrative support, Shahriar Sheikhbahaei for assistance in processing marmoset tissue, Tsai-wen Chen, Hod Dana, and Steve Van Hooser for assistance in acquiring and analyzing visual responses, Catherine Bone for scientific illustrations, Anne N. Connor for editing the manuscript, Janelia AAV core for packaging services, Don B. Arnold and Yang Li for sharing tools. This work was supported by R37NS036715 (R.L.H.), KBRI Basic Research Program (23-BR-02-02 to J.K.), and U01DA056556 (I.H. and R.L.H.).

## Author contributions

Generation of 2P imaging data: I.H., T.Ha., T.C., Generation of electrophysiology data: I.H., J.K., Generation of RNA-seq data: I.H., D.W.K., R.C.J., N.L., T.Hw., F.M.K. Generation and validation of mice: I.H., Z.Y., D.C., A.A., Generation of marmoset data: I.H., S.H.P., D.A.L., Data analysis: I.H., J.K., T.Ha., D.W.K., R.C.J., S.H.P., N.L., Z.Y., D.C., T.Hw., A.A., T.C., F.M.K., S.A.M., X.D., D.A.L., S.B., D.E.B., M.B., S.P.B., R.L.H. Data interpretation: I.H., J.K., T.Ha., D.W.K., S.H.P., S.B., D.E.B., M.B., S.P.B., R.L.H., Writing manuscript: I.H., J.K., T.Ha., D.A.L., D.E.B., M.B., S.P.B., R.L.H.

## References

1 Destexhe, A. & Marder, E. Plasticity in single neuron and circuit computations. Nature 431, 789–795 (2004). https://doi.org:10.1038/na-ture03011

2 Olshausen, B. A. & Field, D. J. Sparse coding of sensory inputs. Curr Opin Neurobiol 14, 481–487 (2004). https://doi.org:10.1016/j.conb.2004.07.007

3 Moser, M. B., Rowland, D. C. & Moser, E. I. Place cells, grid cells, and memory. Cold Spring Harb Perspect Biol 7, a021808 (2015). https://doi.org:10.1101/cshperspect.a021808

4 Niell, C. M. Cell types, circuits, and receptive fields in the mouse visual cortex. Annu Rev Neurosci 38, 413–431 (2015). https://doi.org:10.1146/annurev-neuro-071714-033807

5 Hu, H., Gan, J. & Jonas, P. Interneurons. Fast-spiking, parvalbumin(+) GABAergic interneurons: from cellular design to microcircuit function. Science 345, 1255263 (2014). https://doi.org:10.1126/sci-ence.1255263

6 Huang, Z. J. & Paul, A. The diversity of GABAergic neurons and neural communication elements. Nat Rev Neurosci 20, 563–572 (2019). https://doi.org:10.1038/s41583-019-0195-4

7 Matta, J. A. et al. Developmental origin dictates interneuron AMPA and NMDA receptor subunit composition and plasticity. Nat Neurosci 16, 1032–1041 (2013). https://doi.org:10.1038/nn.3459

8 Petersen, C. C. & Crochet, S. Synaptic computation and sensory processing in neocortical layer 2/3. Neuron 78, 28–48 (2013). https://doi.org:10.1016/j.neuron.2013.03.020

9 Huganir, R. L. & Nicoll, R. A. AMPARs and synaptic plasticity: the last 25 years. Neuron 80, 704–717 (2013). https://doi.org:10.1016/j.neu-ron.2013.10.025

10 Malinow, R. & Malenka, R. C. AMPA receptor trafficking and synaptic plasticity. Annu Rev Neurosci 25, 103–126 (2002). https://doi.org:10.1146/annurev.neuro.25.112701.142758

11 Zeng, H. & Sanes, J. R. Neuronal cell-type classification: challenges, opportunities and the path forward. Nat Rev Neurosci 18, 530–546 (2017). https://doi.org:10.1038/nrn.2017.85

12 Fishell, G. & Kepecs, A. Interneuron Types as Attractors and Controllers. Annu Rev Neurosci 43, 1–30 (2020). https://doi.org:10.1146/an-nurev-neuro-070918-050421

13 Paul, A. et al. Transcriptional Architecture of Synaptic Communication Delineates GABAergic Neuron Identity. Cell 171, 522–539 e520 (2017). https://doi.org:10.1016/j.cell.2017.08.032

14 Tasic, B. et al. Adult mouse cortical cell taxonomy revealed by single cell transcriptomics. Nat Neurosci 19, 335–346 (2016). https://doi.org:10.1038/nn.4216

15 Seabrook, T. A., Burbridge, T. J., Crair, M. C. & Huberman, A. D. Architecture, Function, and Assembly of the Mouse Visual System. Annu Rev Neurosci 40, 499–538 (2017). https://doi.org:10.1146/an-nurev-neuro-071714-033842

16 Isaacson, J. S. & Scanziani, M. How inhibition shapes cortical activity. Neuron 72, 231–243 (2011). https://doi.org:10.1016/j.neu-ron.2011.09.027

17 Sohya, K., Kameyama, K., Yanagawa, Y., Obata, K. & Tsumoto, T. GA-BAergic neurons are less selective to stimulus orientation than excitatory neurons in layer II/III of visual cortex, as revealed by in vivo functional Ca2+ imaging in transgenic mice. J Neurosci 27, 2145–2149 (2007). https://doi.org:10.1523/JNEUROSCI.4641-06.2007

18 Nowak, L. G., Sanchez-Vives, M. V. & McCormick, D. A. Lack of orientation and direction selectivity in a subgroup of fast-spiking inhibitory interneurons: cellular and synaptic mechanisms and comparison with other electrophysiological cell types. Cereb Cortex 18, 1058–1078 (2008). https://doi.org:10.1093/cercor/bhm137

19 Liu, B. H. et al. Visual receptive field structure of cortical inhibitory neurons revealed by two-photon imaging guided recording. J Neurosci 29, 10520–10532 (2009). https://doi.org:10.1523/JNEUROSCI.1915-09.2009

20 Kerlin, A. M., Andermann, M. L., Berezovskii, V. K. & Reid, R. C. Broadly tuned response properties of diverse inhibitory neuron subtypes in mouse visual cortex. Neuron 67, 858–871 (2010). https://doi.org:10.1016/j.neuron.2010.08.002

21 Niell, C. M. & Stryker, M. P. Highly selective receptive fields in mouse visual cortex. J Neurosci 28, 7520–7536 (2008). https://doi.org:10.1523/JNEUROSCI.0623-08.2008

22 Runyan, C. A. et al. Response features of parvalbumin-expressing interneurons suggest precise roles for subtypes of inhibition in visual cortex. Neuron 67, 847–857 (2010). https://doi.org:10.1016/j.neu-ron.2010.08.006

23 Runyan, C. A. & Sur, M. Response selectivity is correlated to dendritic structure in parvalbumin-expressing inhibitory neurons in visual cortex. J Neurosci 33, 11724–11733 (2013). https://doi.org:10.1523/JNEU-ROSCI.2196-12.2013

24 Jonas, P., Racca, C., Sakmann, B., Seeburg, P. H. & Monyer, H. Differences in Ca2+ permeability of AMPA-type glutamate receptor channels in neocortical neurons caused by differential GluR-B subunit expression. Neuron 12, 1281–1289 (1994). https://doi.org:10.1016/0896-6273(94)90444-8

25 Geiger, J. R. et al. Relative abundance of subunit mRNAs determines gating and Ca2+ permeability of AMPA receptors in principal neurons and interneurons in rat CNS. Neuron 15, 193–204 (1995). https://doi.org:10.1016/0896-6273(95)90076-4

26 Ozawa, S., Iino, M. & Tsuzuki, K. Two types of kainate response in cultured rat hippocampal neurons. J Neurophysiol 66, 2–11 (1991). https://doi.org:10.1152/jn.1991.66.1.2

27 McBain, C. J. & Dingledine, R. Heterogeneity of synaptic glutamate receptors on CA3 stratum radiatum interneurones of rat hippocampus. J Physiol 462, 373–392 (1993). https://doi.org:10.1113/jphys-iol.1993.sp019560

28 Traynelis, S. F. et al. Glutamate receptor ion channels: structure, regulation, and function. Pharmacol Rev 62, 405–496 (2010). https://doi.org:10.1124/pr.109.002451

29 Alle, H., Jonas, P. & Geiger, J. R. PTP and LTP at a hippocampal mossy fiber-interneuron synapse. Proc Natl Acad Sci U S A 98, 14708–14713 (2001). https://doi.org:10.1073/pnas.251610898

30 Lamsa, K. P., Heeroma, J. H., Somogyi, P., Rusakov, D. A. & Kullmann, D. M. Anti-Hebbian long-term potentiation in the hippocampal feedback inhibitory circuit. Science 315, 1262–1266 (2007). https://doi.org:10.1126/science.1137450

31 Gu, J. G., Albuquerque, C., Lee, C. J. & MacDermott, A. B. Synaptic strengthening through activation of Ca2+-permeable AMPA receptors. Nature 381, 793–796 (1996). https://doi.org:10.1038/381793a0

32 Mahanty, N. K. & Sah, P. Calcium-permeable AMPA receptors mediate long-term potentiation in interneurons in the amygdala. Nature 394, 683–687 (1998). https://doi.org:10.1038/29312

33 Isaac, J. T., Ashby, M. C. & McBain, C. J. The role of the GluR2 subunit in AMPA receptor function and synaptic plasticity. Neuron 54, 859–871 (2007). https://doi.org:10.1016/j.neuron.2007.06.001

34 Sommer, B., Kohler, M., Sprengel, R. & Seeburg, P. H. RNA editing in brain controls a determinant of ion flow in glutamate-gated channels. Cell 67, 11–19 (1991). https://doi.org:10.1016/0092-8674(91)90568-j

35 Lake, B. B. et al. Neuronal subtypes and diversity revealed by singlenucleus RNA sequencing of the human brain. Science 352, 1586–1590 (2016). https://doi.org:10.1126/science.aaf1204

36 Krienen, F. M. et al. Innovations present in the primate interneuron repertoire. Nature 586, 262–269 (2020). https://doi.org:10.1038/s41586-020-2781-z

37 Khawaja, R. R. et al. GluA2 overexpression in oligodendrocyte progenitors promotes postinjury oligodendrocyte regeneration. Cell Rep 35, 109147 (2021). https://doi.org:10.1016/j.celrep.2021.109147

38 Hippenmeyer, S. et al. A developmental switch in the response of DRG neurons to ETS transcription factor signaling. PLoS Biol 3, e159 (2005). https://doi.org:10.1371/journal.pbio.0030159

39 Carter, A. G. & Sabatini, B. L. State-dependent calcium signaling in dendritic spines of striatal medium spiny neurons. Neuron 44, 483–493 (2004). https://doi.org:10.1016/j.neuron.2004.10.013

40 Goldberg, J. H., Tamas, G., Aronov, D. & Yuste, R. Calcium microdo-mains in aspiny dendrites. Neuron 40, 807–821 (2003). https://doi.org:10.1016/s0896-6273(03)00714-1

41 Scholl, B., Thomas, C. I., Ryan, M. A., Kamasawa, N. & Fitzpatrick, D. Cortical response selectivity derives from strength in numbers of synapses. Nature 590, 111–114 (2021). https://doi.org:10.1038/s41586-020-03044-3

42 Rossi, L. F., Harris, K. D. & Carandini, M. Spatial connectivity matches direction selectivity in visual cortex. Nature 588, 648–652 (2020). https://doi.org:10.1038/s41586-020-2894-4

43 Scholl, B., Pattadkal, J. J., Dilly, G. A., Priebe, N. J. & Zemelman, B. V. Local Integration Accounts for Weak Selectivity of Mouse Neocortical Parvalbumin Interneurons. Neuron 87, 424–436 (2015). https://doi.org:10.1016/j.neuron.2015.06.030

44 Hofer, S. B. et al. Differential connectivity and response dynamics of excitatory and inhibitory neurons in visual cortex. Nat Neurosci 14, 1045–1052 (2011). https://doi.org:10.1038/nn.2876

45 Bock, D. D. et al. Network anatomy and in vivo physiology of visual cortical neurons. Nature 471, 177–182 (2011). https://doi.org:10.1038/nature09802

46 Packer, A. M. & Yuste, R. Dense, unspecific connectivity of neocortical parvalbumin-positive interneurons: a canonical microcircuit for inhibition? J Neurosci 31, 13260–13271 (2011). https://doi.org:10.1523/JNEUROSCI.3131-11.2011

47 Lu, J. T., Li, C. Y., Zhao, J. P., Poo, M. M. & Zhang, X. H. Spike-timing-dependent plasticity of neocortical excitatory synapses on inhibitory interneurons depends on target cell type. J Neurosci 27, 9711–9720 (2007). https://doi.org:10.1523/JNEUROSCI.2513-07.2007

48 Le Duigou, C., Savary, E., Kullmann, D. M. & Miles, R. Induction of Anti-Hebbian LTP in CA1 Stratum Oriens Interneurons: Interactions between Group I Metabotropic Glutamate Receptors and M1 Muscarinic Receptors. J Neurosci 35, 13542–13554 (2015). https://doi.org:10.1523/JNEUROSCI.0956-15.2015

49 Kawaguchi, Y. & Kubota, Y. GABAergic cell subtypes and their synaptic connections in rat frontal cortex. Cereb Cortex 7, 476–486 (1997). https://doi.org:10.1093/cercor/7.6.476

50 Geiger, J. R., Lubke, J., Roth, A., Frotscher, M. & Jonas, P. Submillisecond AMPA receptor-mediated signaling at a principal neuron-in-terneuron synapse. Neuron 18, 1009–1023 (1997). https://doi.org:10.1016/s0896-6273(00)80339-6

51 Galarreta, M. & Hestrin, S. Spike transmission and synchrony detection in networks of GABAergic interneurons. Science 292, 2295–2299 (2001). https://doi.org:10.1126/science.1061395

52 Goetz, L., Roth, A. & Hausser, M. Active dendrites enable strong but sparse inputs to determine orientation selectivity. Proc Natl Acad Sci U S A 118 (2021). https://doi.org:10.1073/pnas.2017339118

53 Jia, Z. et al. Enhanced LTP in mice deficient in the AMPA receptor GluR2. Neuron 17, 945–956 (1996). https://doi.org:10.1016/s0896-6273(00)80225-1

54 Meng, Y., Zhang, Y. & Jia, Z. Synaptic transmission and plasticity in the absence of AMPA glutamate receptor GluR2 and GluR3. Neuron 39, 163–176 (2003). https://doi.org:10.1016/s0896-6273(03)00368-4

55 Hainmueller, T. & Bartos, M. Parallel emergence of stable and dynamic memory engrams in the hippocampus. Nature 558, 292–296 (2018). https://doi.org:10.1038/s41586-018-0191-2

56 Derkach, V. A., Oh, M. C., Guire, E. S. & Soderling, T. R. Regulatory mechanisms of AMPA receptors in synaptic plasticity. Nat Rev Neurosci 8, 101–113 (2007). https://doi.org:10.1038/nrn2055

57 Najafi, F. et al. Excitatory and Inhibitory Subnetworks Are Equally Selective during Decision-Making and Emerge Simultaneously during Learning. Neuron 105, 165–179 e168 (2020). https://doi.org:10.1016/j.neuron.2019.09.045

58 Moore, A. K. & Wehr, M. Parvalbumin-expressing inhibitory interneurons in auditory cortex are well-tuned for frequency. J Neurosci 33, 13713–13723 (2013). https://doi.org:10.1523/JNEUROSCI.0663-13.2013

59 Li, L. Y. et al. Differential Receptive Field Properties of Parvalbumin and Somatostatin Inhibitory Neurons in Mouse Auditory Cortex. Cereb Cortex 25, 1782–1791 (2015). https://doi.org:10.1093/cer-cor/bht417

60 Znamenskiy, P. et al. Functional selectivity and specific connectivity of inhibitory neurons in primary visual cortex. bioRxiv (2018).

61 Karnani, M. M., Agetsuma, M. & Yuste, R. A blanket of inhibition: functional inferences from dense inhibitory connectivity. Curr Opin Neurobiol 26, 96–102 (2014). https://doi.org:10.1016/j.conb.2013.12.015

62 Yoshimura, Y. & Callaway, E. M. Fine-scale specificity of cortical net-works depends on inhibitory cell type and connectivity. Nat Neurosci 8, 1552–1559 (2005). https://doi.org:10.1038/nn1565

63 Tosches, M. A. et al. Evolution of pallium, hippocampus, and cortical cell types revealed by single-cell transcriptomics in reptiles. Science 360, 881–888 (2018). https://doi.org:10.1126/science.aar4237

64 Salpietro, V. et al. AMPA receptor GluA2 subunit defects are a cause of neurodevelopmental disorders. Nat Commun 10, 3094 (2019). https://doi.org:10.1038/s41467-019-10910-w

65 Bugeon, S. et al. A transcriptomic axis predicts state modulation of cortical interneurons. Nature 607, 330–338 (2022). https://doi.org:10.1038/s41586-022-04915-7

66 Debanne, D., Inglebert, Y. & Russier, M. Plasticity of intrinsic neuronal excitability. Curr Opin Neurobiol 54, 73–82 (2019). https://doi.org:10.1016/j.conb.2018.09.001

67 Pehlevan, C., Hu, T. & Chklovskii, D. B. A Hebbian/Anti-Hebbian Neural Network for Linear Subspace Learning: A Derivation from Multidimensional Scaling of Streaming Data. Neural Comput 27, 1461–1495 (2015). https://doi.org:10.1162/NECO_a_00745

68 Carandini, M. & Heeger, D. J. Normalization as a canonical neural computation. Nat Rev Neurosci 13, 51–62 (2011). https://doi.org:10.1038/nrn3136

## Methods References

69 Sousa, V. H., Miyoshi, G., Hjerling-Leffler, J., Karayannis, T. & Fishell, G. Characterization of Nkx6-2-derived neocortical interneuron lineages. Cereb Cortex 19 Suppl 1, i1–10 (2009). https://doi.org:10.1093/cercor/bhp038

70 Zamanillo, D. et al. Importance of AMPA receptors for hippocampal synaptic plasticity but not for spatial learning. Science 284, 1805–1811 (1999). https://doi.org:10.1126/science.284.5421.1805

71 Madisen, L. et al. A robust and high-throughput Cre reporting and characterization system for the whole mouse brain. Nat Neurosci 13, 133–140 (2010). https://doi.org:10.1038/nn.2467

72 Sekine-Aizawa, Y. & Huganir, R. L. Imaging of receptor trafficking by using alpha-bungarotoxin-binding-site-tagged receptors. Proc Natl Acad Sci U S A 101, 17114–17119 (2004). https://doi.org:10.1073/pnas.0407563101

73 Dana, H. et al. Sensitive red protein calcium indicators for imaging neural activity. Elife 5 (2016). https://doi.org:10.7554/eLife.12727

74 Choi, J. H. et al. Optimization of AAV expression cassettes to improve packaging capacity and transgene expression in neurons. Mol Brain 7, 17 (2014). https://doi.org:10.1186/1756-6606-7-17

75 Dong, J. Y., Fan, P. D. & Frizzell, R. A. Quantitative analysis of the packaging capacity of recombinant adeno-associated virus. Hum Gene Ther 7, 2101–2112 (1996). https://doi.org:10.1089/hum.1996.7.17-2101

76 Hoess, R., Wierzbicki, A. & Abremski, K. Formation of small circular DNA molecules via an in vitro site-specific recombination system. Gene 40, 325–329 (1985). https://doi.org:10.1016/0378-1119(85)90056-3

77 Araki, Y., Lin, D. T. & Huganir, R. L. Plasma membrane insertion of the AMPA receptor GluA2 subunit is regulated by NSF binding and Q/R editing of the ion pore. Proc Natl Acad Sci U S A 107, 11080–11085 (2010). https://doi.org:10.1073/pnas.1006584107

78 Wekselblatt, J. B., Flister, E. D., Piscopo, D. M. & Niell, C. M. Largescale imaging of cortical dynamics during sensory perception and behavior. J Neurophysiol 115, 2852–2866 (2016). https://doi.org:10.1152/jn.01056.2015

79 Juavinett, A. L., Nauhaus, I., Garrett, M. E., Zhuang, J. & Callaway, E. M. Automated identification of mouse visual areas with intrinsic signal imaging. Nat Protoc 12, 32–43 (2017). https://doi.org:10.1038/nprot.2016.158

80 Pnevmatikakis, E. A. & Giovannucci, A. NoRMCorre: An online algorithm for piecewise rigid motion correction of calcium imaging data. J Neurosci Methods 291, 83–94 (2017). https://doi.org:10.1016/j.jneumeth.2017.07.031

81 Giovannucci, A. et al. CaImAn an open source tool for scalable calcium imaging data analysis. Elife 8 (2019). https://doi.org:10.7554/eLife.38173

82 Kleiner, M. et al. What’s new in psychtoolbox-3. Perception 36, 1–16 (2007).

83 Muir, D. R. & Kampa, B. M. FocusStack and StimServer: a new open source MATLAB toolchain for visual stimulation and analysis of two-photon calcium neuronal imaging data. Front Neuroinform 8, 85 (2014). https://doi.org:10.3389/fninf.2014.00085

84 Chen, T. W. et al. Ultrasensitive fluorescent proteins for imaging neuronal activity. Nature 499, 295–300 (2013). https://doi.org:10.1038/na-ture12354

85 Mazurek, M., Kager, M. & Van Hooser, S. D. Robust quantification of orientation selectivity and direction selectivity. Front Neural Circuits 8, 92 (2014). https://doi.org:10.3389/fncir.2014.00092

86 De Valois, R. L., Yund, E. W. & Hepler, N. The orientation and direction selectivity of cells in macaque visual cortex. Vision Res 22, 531–544 (1982). https://doi.org:10.1016/0042-6989(82)90112-2

87 Li, Y., Van Hooser, S. D., Mazurek, M., White, L. E. & Fitzpatrick, D. Experience with moving visual stimuli drives the early development of cortical direction selectivity. Nature 456, 952–956 (2008). https://doi.org:10.1038/nature07417

88 Sclar, G. & Freeman, R. D. Orientation selectivity in the cat’s striate cortex is invariant with stimulus contrast. Exp Brain Res 46, 457–461 (1982). https://doi.org:10.1007/BF00238641

89 Kaifosh, P., Zaremba, J. D., Danielson, N. B. & Losonczy, A. SIMA: Python software for analysis of dynamic fluorescence imaging data. Front Neuroinform 8, 80 (2014). https://doi.org:10.3389/fninf.2014.00080

90 Pachitariu, M. et al. Suite2p: beyond 10,000 neurons with standard two-photon microscopy. BioRxiv, 061507 (2016).

91 Sheffield, M. E. J., Adoff, M. D. & Dombeck, D. A. Increased Prevalence of Calcium Transients across the Dendritic Arbor during Place Field Formation. Neuron 96, 490–504 e495 (2017). https://doi.org:10.1016/j.neuron.2017.09.029

92 Kubie, J. L., Muller, R. U. & Bostock, E. Spatial firing properties of hippocampal theta cells. J Neurosci 10, 1110–1123 (1990). https://doi.org:10.1523/JNEUROSCI.10-04-01110.1990

93 Frankish, A. et al. GENCODE reference annotation for the human and mouse genomes. Nucleic Acids Res 47, D766–D773 (2019). https://doi.org:10.1093/nar/gky955

94 Dobin, A. et al. STAR: ultrafast universal RNA-seq aligner. Bioinformatics 29, 15–21 (2013). https://doi.org:10.1093/bioinformatics/bts635

95 Tasic, B. et al. Shared and distinct transcriptomic cell types across neocortical areas. Nature 563, 72–78 (2018). https://doi.org:10.1038/s41586-018-0654-5

96 Joseph, D. J. et al. Protocol for isolating young adult parvalbumin interneurons from the mouse brain for extraction of high-quality RNA. STAR Protoc 2, 100714 (2021). https://doi.org:10.1016/j.xpro.2021.100714

97 Zeisel, A. et al. Brain structure. Cell types in the mouse cortex and hippocampus revealed by single-cell RNA-seq. Science 347, 1138–1142 (2015). https://doi.org:10.1126/science.aaa1934

98 Saxena, A. et al. Trehalose-enhanced isolation of neuronal sub-types from adult mouse brain. Biotechniques 52, 381–385 (2012). https://doi.org:10.2144/0000113878

99 Patro, R., Mount, S. M. & Kingsford, C. Sailfish enables alignment-free isoform quantification from RNA-seq reads using lightweight algorithms. Nat Biotechnol 32, 462–464 (2014). https://doi.org:10.1038/nbt.2862

100 Love, M. I., Huber, W. & Anders, S. Moderated estimation of fold change and dispersion for RNA-seq data with DESeq2. Genome Biol 15, 550 (2014). https://doi.org:10.1186/s13059-014-0550-8

101 Le Roux, N., Cabezas, C., Bohm, U. L. & Poncer, J. C. Input-specific learning rules at excitatory synapses onto hippocampal parvalbumin-expressing interneurons. J Physiol 591, 1809–1822 (2013). https://doi.org:10.1113/jphysiol.2012.245852

102 Hong, I. et al. AMPA receptor exchange underlies transient memory destabilization on retrieval. Proc Natl Acad Sci U S A 110, 8218–8223 (2013). https://doi.org:10.1073/pnas.1305235110

103 Clem, R. L. & Huganir, R. L. Calcium-permeable AMPA receptor dynamics mediate fear memory erasure. Science 330, 1108–1112 (2010). https://doi.org:10.1126/science.1195298

104 Rodriguez, C. I. et al. High-efficiency deleter mice show that FLPe is an alternative to Cre-loxP. Nat Genet 25, 139–140 (2000). https://doi.org:10.1038/75973

105 Hollmann, M., Hartley, M. & Heinemann, S. Ca2+ permeability of KA-AMPA--gated glutamate receptor channels depends on subunit composition. Science 252, 851–853 (1991). https://doi.org:10.1126/sci-ence.1709304

106 Verdoorn, T. A., Burnashev, N., Monyer, H., Seeburg, P. H. & Sakmann, B. Structural determinants of ion flow through recombinant glutamate receptor channels. Science 252, 1715–1718 (1991). https://doi.org:10.1126/science.1710829

