## Extended Data Figures for "Calcium-permeable AMPA receptors govern PV neuron feature selectivity"

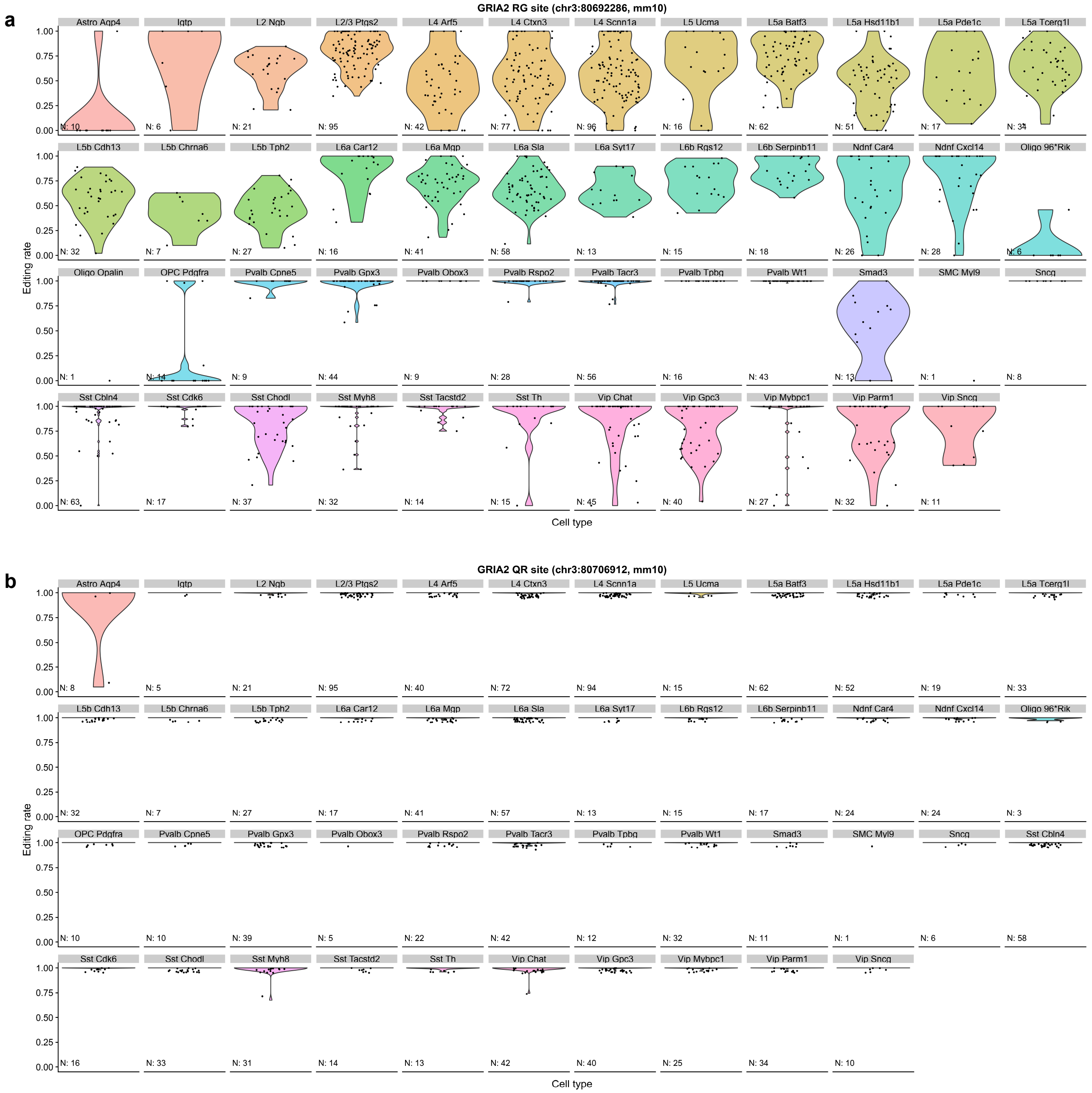


**Extended Data Figure 1 | RNA editing at the *Gria2* Q/R site is largely complete across many cortical cell types. a,** A-to-I RNA editing rates at the *Gria2* R/G site. Editing rates [G/(A+G)] were stratified according to the cell types defined in Tasic et al.[9]. Each dot represents the editing rate in a single cell. The number of samples (cells) is noted in each panel. **b,** A-to-I RNA editing rates at the *Gria2* Q/R editing site. Due to high concentration of data near 1 (complete RNA editing) in this panel, many violin symbols were not visible, and single data points were jittered by 0.05 along the y-axis to aid visualization.

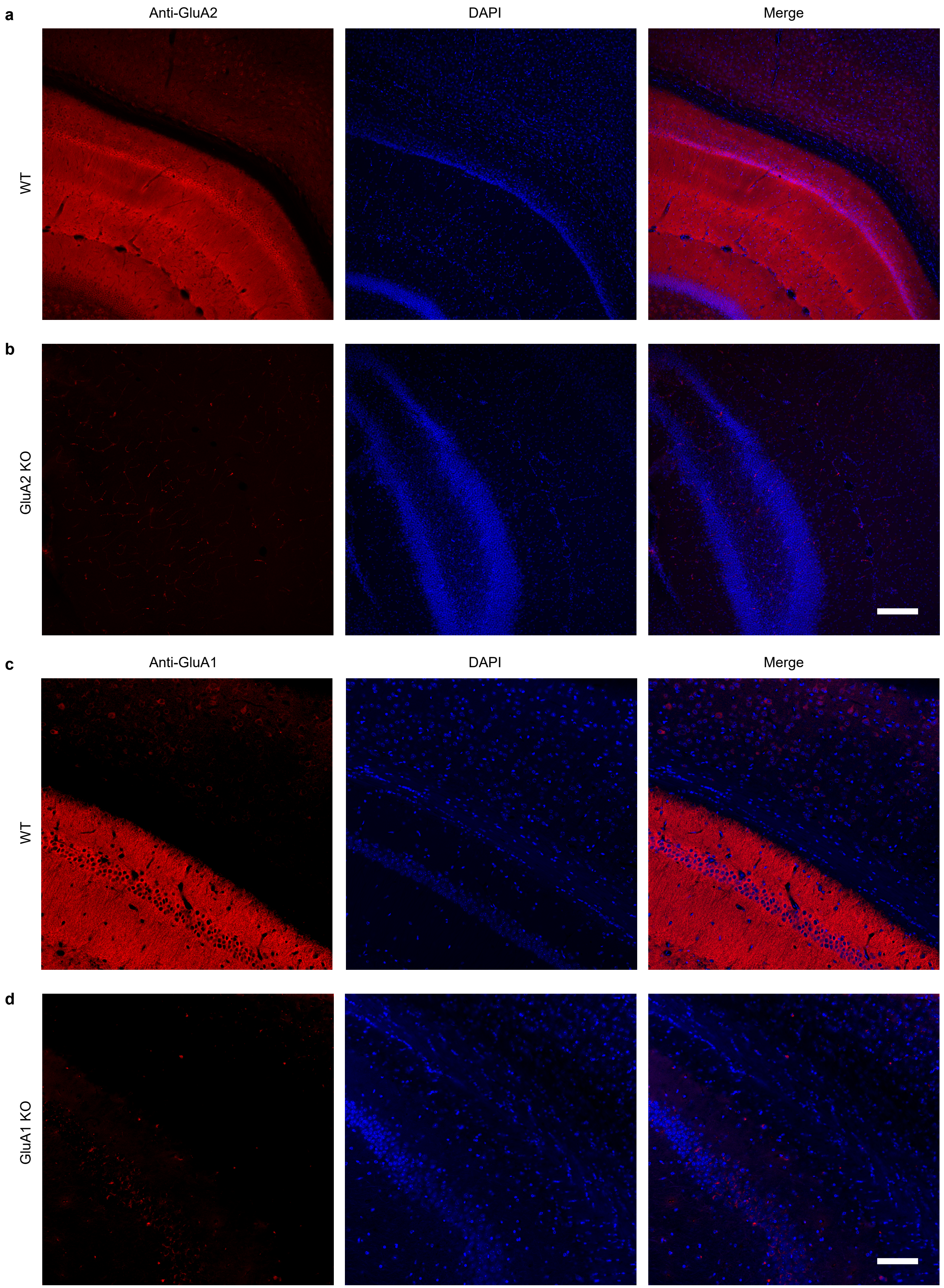


**Extended Data Figure 2 | Immunohistochemical validation of anti-GluA2 and anti-GluA1 antibodies. a-d,** Immunohistochemical staining of GluA2 in GluA2$^{-/-}$ knockout mice (KO, **b**), wild-type littermates (WT, **a**), staining of GluA1 in GluA1$^{-/-}$ knockout mice (KO, **d**), and wild-type littermates (WT, **c**). Images of hippocampus and visual cortex were taken at 20x magnification. Scale bars, 200 µm.

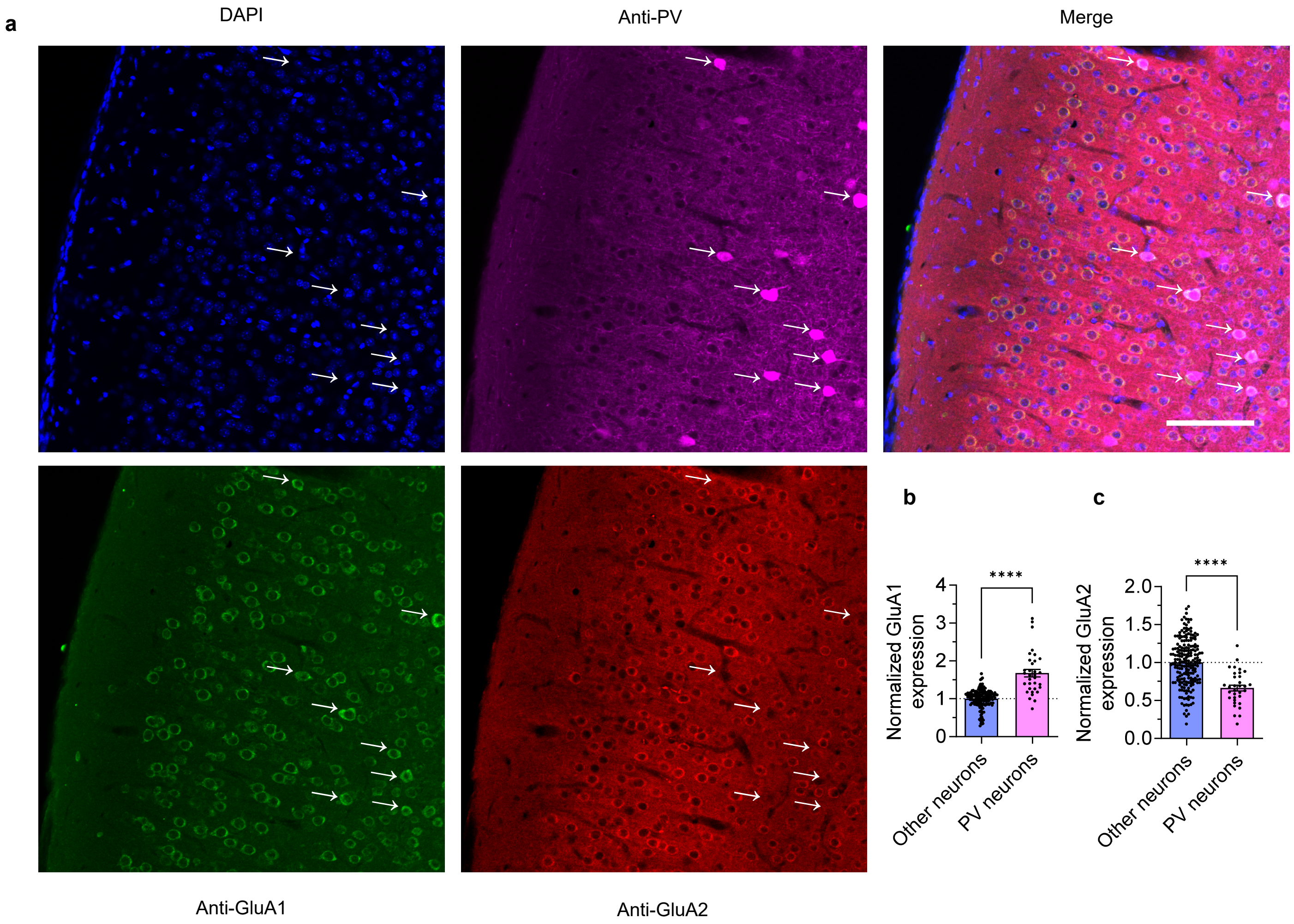


**Extended Data Figure 3 | High expression of GluA1 and low expression of GluA2 protein in PV interneurons. a,** Immunohistochemical staining of PV, GluA1, and GluA2 in layer 2/3 of visual cortex. PV interneurons (arrows) show markedly lower GluA2 expression, and higher GluA1 expression compared to all other neurons (all GluA1+ or GluA2+ and DAPI+ cells). Scale bar, 100 μm. **b,** Quantification of GluA1 expression (mean ± SEM) shows that PV interneurons express significantly more GluA1 (other neurons: 1.00 ± 0.02, n = 203 neurons/3 slices/3 mice; PV interneurons: 1.68 ± 0.10, n = 33 neurons; P < 0.0001, unpaired t-test). **c,** Quantification of GluA2 expression (mean ± SEM) shows that PV interneurons express significantly less GluA2 (other neurons: 1.00 ± 0.02; PV interneurons: 0.66 ± 0.04; P < 0.0001, unpaired t-test).

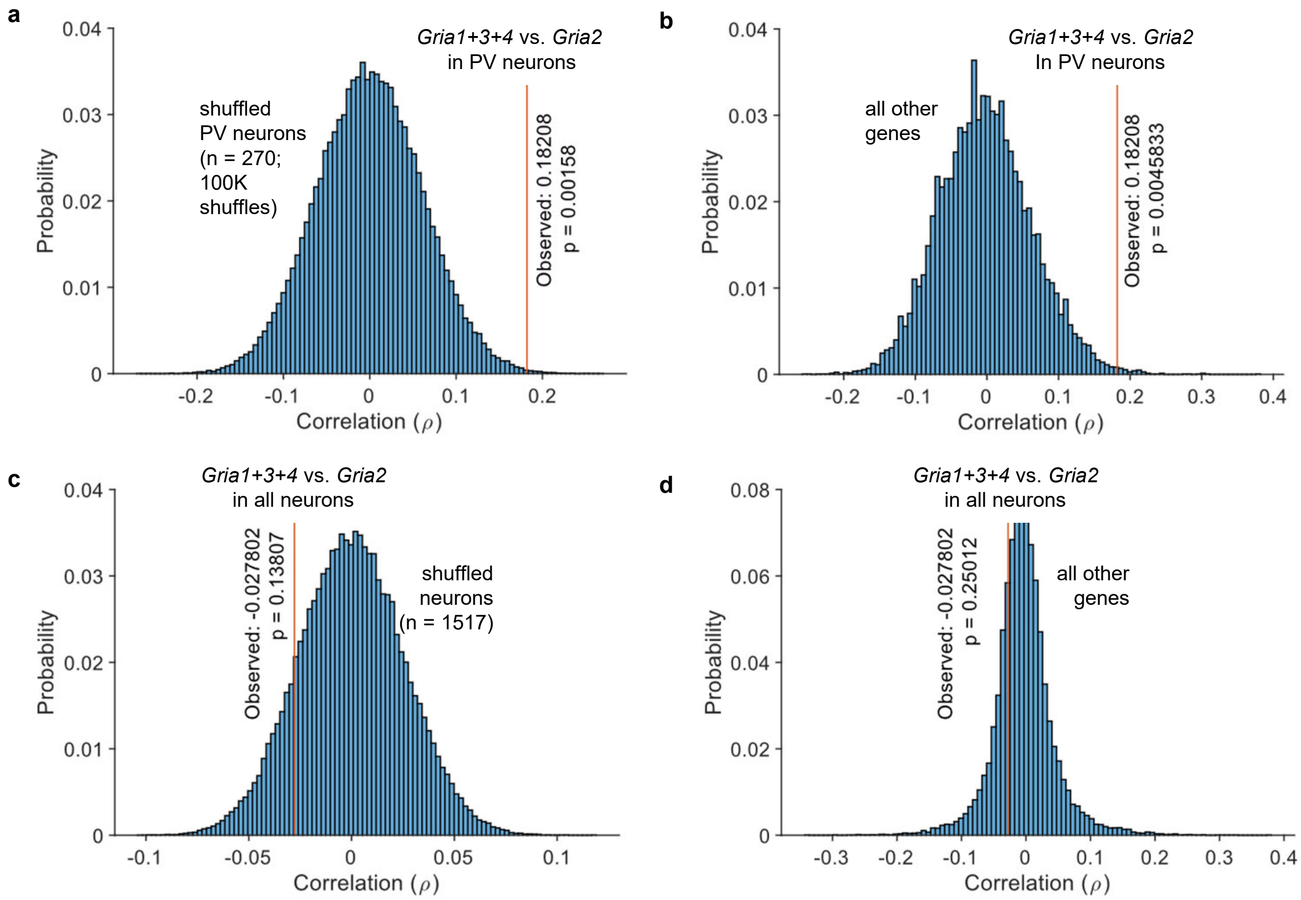


**Extended Data Figure 4 | Potential co-regulation of *Gria1-4* mRNA expression in PV interneurons. a,** Single-cell mRNA expression[9] of *Gria2* in PV neurons showed a strong correlation (ρ = 0.18208, n = 270 cells) with the sum of *Gria1, Gria3, Gria4* mRNA expression, which was highly significant compared to a bootstrap randomized distribution (100,000 shuffles across PV neurons, P = 0.0019). The Monte Carlo P-value was determined by comparing the observed correlation statistic to the simulated distribution. **b,** The correlation between *Gria2* expression and the sum of *Gria1, Gria3, Gria4* expression in PV neurons was also highly significant compared to the distribution of correlations with all other genes (top 0.45 percentile, P = 0.0046). **c,d,** This correlation was not present in the entire neuron population (n = 1517 cells, ρ = -0.027802; P = 0.1381 in comparison to shuffled neuron data; P = 0.25012 in comparison to the correlation of *Gria1+2+3* to all other genes beyond *Gria2*). These results suggest a tight co-regulation of *Gria2 vs. Gria1+3+4* mRNA expression ratio unique to PV interneurons.

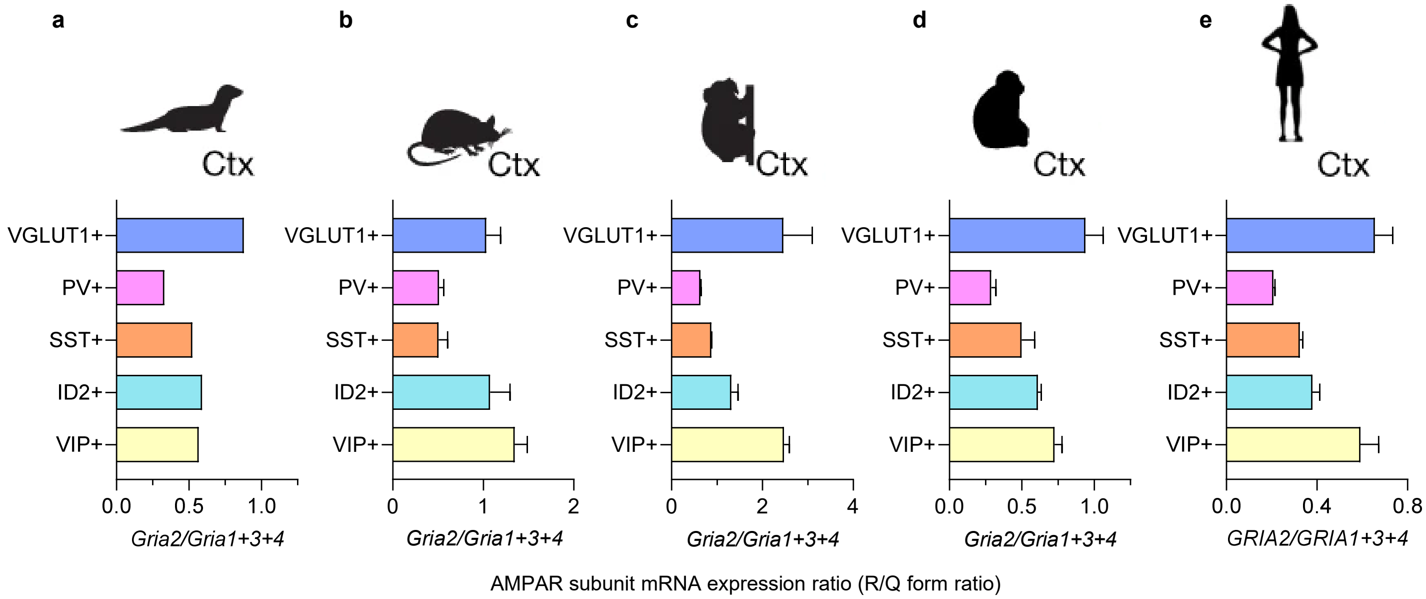


**Extended Data Figure 5 | Conserved low expression of Gria2 mRNA in PV/SST interneurons across mammalian species. a-e,** Analysis of Drop-seq single-cell RNA-seq data[36] from the cortex of (**a**) ferrets, (**b**) mice, (**c**) marmosets, (**d**) macaques, and (**e**) humans shows a conserved lower ratio of calcium impermeable/calcium permeable AMPAR subunits (R/Q form ratio) in PV and SST interneurons. VGLUT1 cells correspond to cortical excitatory neurons (CaMKIIα), and ID2 correspond to neurogliaform cells. Bars and error bars denote mean ± SD of Drop-seq samples.

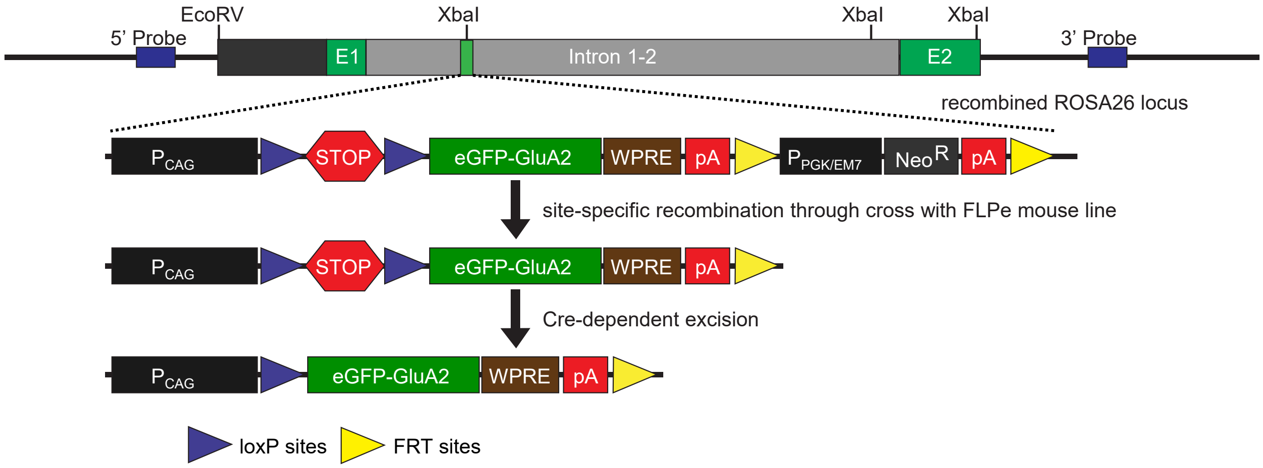


**Extended Data Figure 6 | Development of a Rosa26 knock-in mouse to conditionally express eGFP-tagged GluA2 in a Cre-dependent manner.** To enable robust expression of eGFP-GluA2, we used a strong ubiquitous CMV-βactin hybrid (CAG) promoter (consisting of three gene regulatory elements: 5′ cytomegalovirus early enhancer element, chicken β-actin promoter and rabbit b-globin intron) and added a woodchuck hepatitis virus posttranscriptional regulatory element (WPRE) at the 3′ end of the eGFP-GluA2 coding sequence. The WPRE sequence allows rapid exit of mRNA from the nucleus and increases the mRNA stability in the cytosol. For inducible expression of eGFP-GluA2, a "stopper" cassette consisting of loxP-flanked 3X SV40 polyA (loxP-STOP-loxP, "lsl") was placed upstream of the coding sequence, preventing expression until cyclic recombinase (Cre)-dependent excision. A Neomycin resistance cassette ($Neo^R$) flanked by Flippase Recognition Target sequences (FRT) was present in the targeting vector to allow for selection.

To prevent gene-silencing effects and ensure consistent and long-term expression of these transgenes in all cell types, the CAG-driven inducible eGFP-GluA2 transgenic constructs were targeted to the ubiquitously expressed *Rosa26* locus. For homologous recombination in mouse embryonic stem (ES) cells, the gene-targeting vector was assembled into a ROSA26 targeting plasmid containing a 1.2 kb 5′ homology arm, 4.3 kb 3′ homology arm, and PGK-DTA (Diphtheria toxin fragment A, downstream of 3′ homology arm) for negative selection. ES cells, derived from a SV129 mouse strain, were electroporated with the AsiSI-linearized targeting vectors. A nested PCR screening strategy along the 5′ homology arm was used to identify ES cell clones harboring the correct genomic targeting event. After verification of homologous recombination by Southern blot analysis and confirmation of the karyotypes, correctly targeted ES cell clones were used to generate chimeric mice by injection into blastocysts derived from SV129 females at the Johns Hopkins University Transgenic Core. Germline transmission was achieved by breeding male chimeric founders to C57BL/6N wild-type female mice. The FRT-NeoR cassette was removed by breeding to a transgenic FLPe mouse line[104].

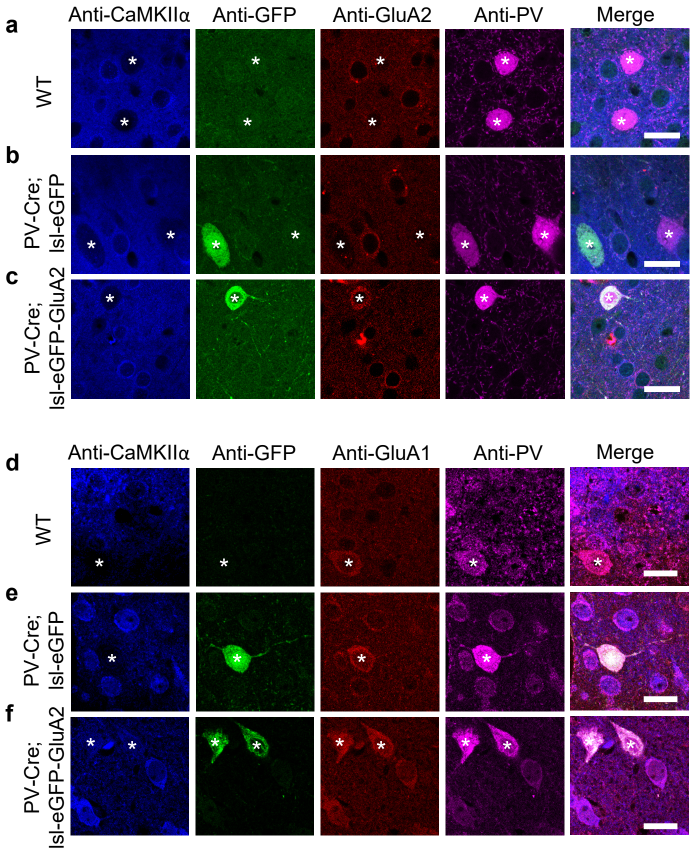


**Extended Data Figure 7 | Transgenic expression of GluA2 in PV interneurons mimics excitatory neuron GluA2 expression levels (related to Fig 2b,c). a-c,** Representative data for Fig. 2b. Immunohistochemical staining of GluA2 expression in PV interneurons and excitatory neurons. PV interneurons are marked by white asterisks and display negative CaMKIIα staining. Images were acquired in layer 2/3 of visual cortex. Scale bars, 15 µm. **d-f,** Representative data for Fig. 2c. Immunohistochemical staining of GluA1 expression in PV interneurons and excitatory neurons. Scale bars, 15 µm.

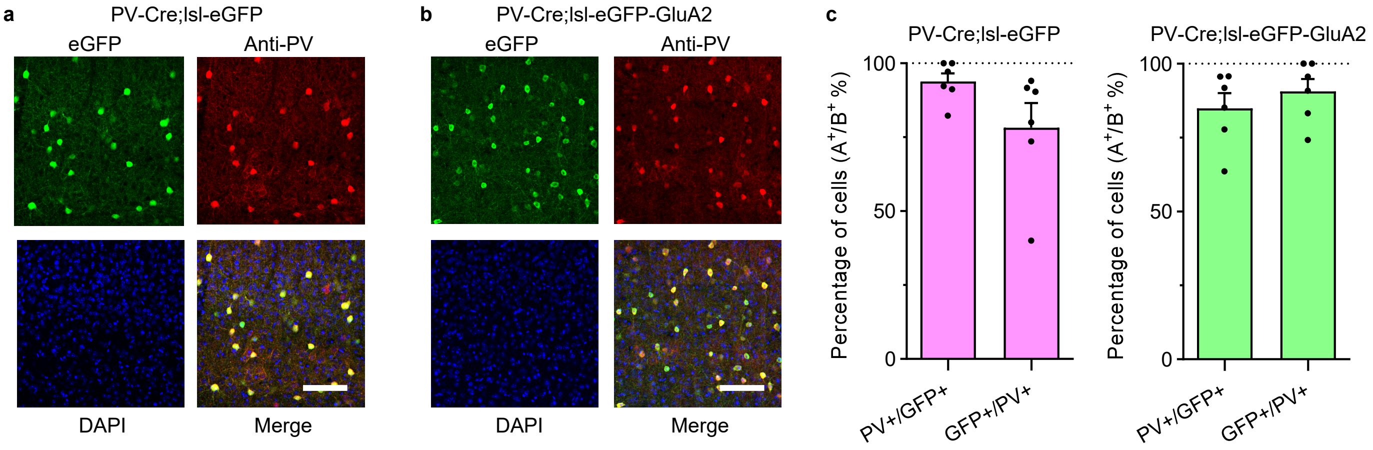


**Extended Data Figure 8 | Concordance of conditionally expressed eGFP and eGFP-GluA2 with PV immunostaining.**
**a,b,** Immunohistochemical staining of PV interneurons in mouse visual cortex. Scale bars, 100 µm. **c,** Quantification of conditional expression concordance in PV interneurons. In both PV-Cre;lsl-eGFP and PV-Cre; lsl-eGFP-GluA2 mouse lines, the ratio of PV+ cells among GFP+ cells and GFP+ cells among PV+ cells was high (n = 6/6 slices for each genotype; PV-Cre;lsl-eGFP mice: PV+/GFP+ = 93.8 ± 2.7%, GFP+/PV+ = 78.3 ± 8.3%; PV-Cre;lsl-eGFP-GluA2 mice: PV+/GFP+ = 85.0 ± 5.1%, GFP+/PV+ = 90.7 ± 4.0%). Bars and error bars denote mean ± SEM.

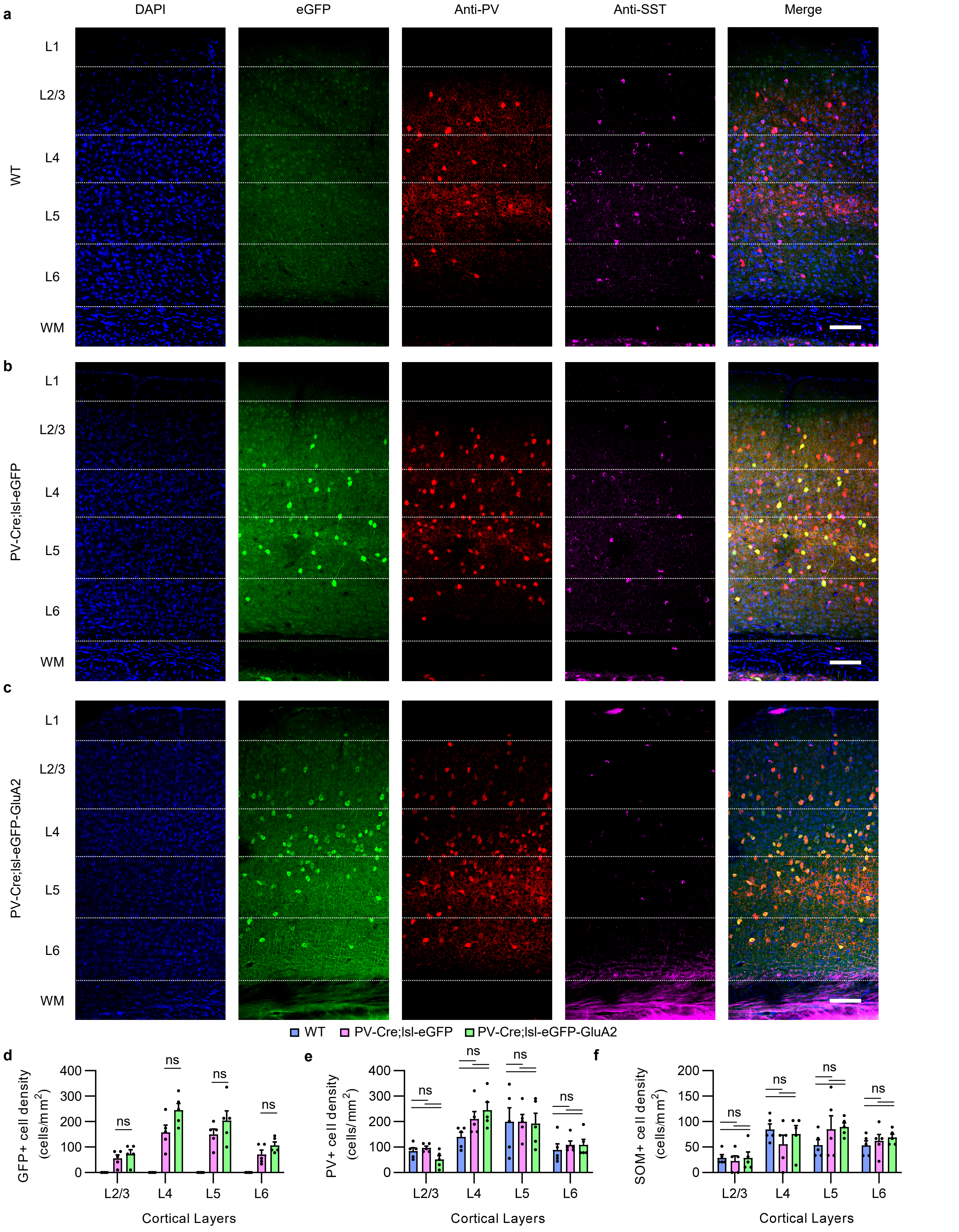


**Extended Data Figure 9 | Transgenic expression of GluA2 in PV interneurons does not alter PV/SST interneuron density. a-c,** Visual cortex immunohistochemical staining of PV/SST interneurons. Layer segmentation was based on marker gene expression staining in V1 of internal and Allen Brain Atlas mouse brain sections. Scale bars, 100 µm. **d-f,** Quantification of GFP+, PV+, and SST+ cell density in cortical layers. PV and SST neuron density did not significantly change in PV-Cre;lsl-eGFP or PV-Cre;lsl-eGFP-GluA2 mice (n = 5/5/5 slices, 3/3/3 mice, 2-way ANOVA, P > 0.05 for all post-hoc comparisons, Šídák's multiple comparison correction). Bars and error bars denote mean ± SEM.

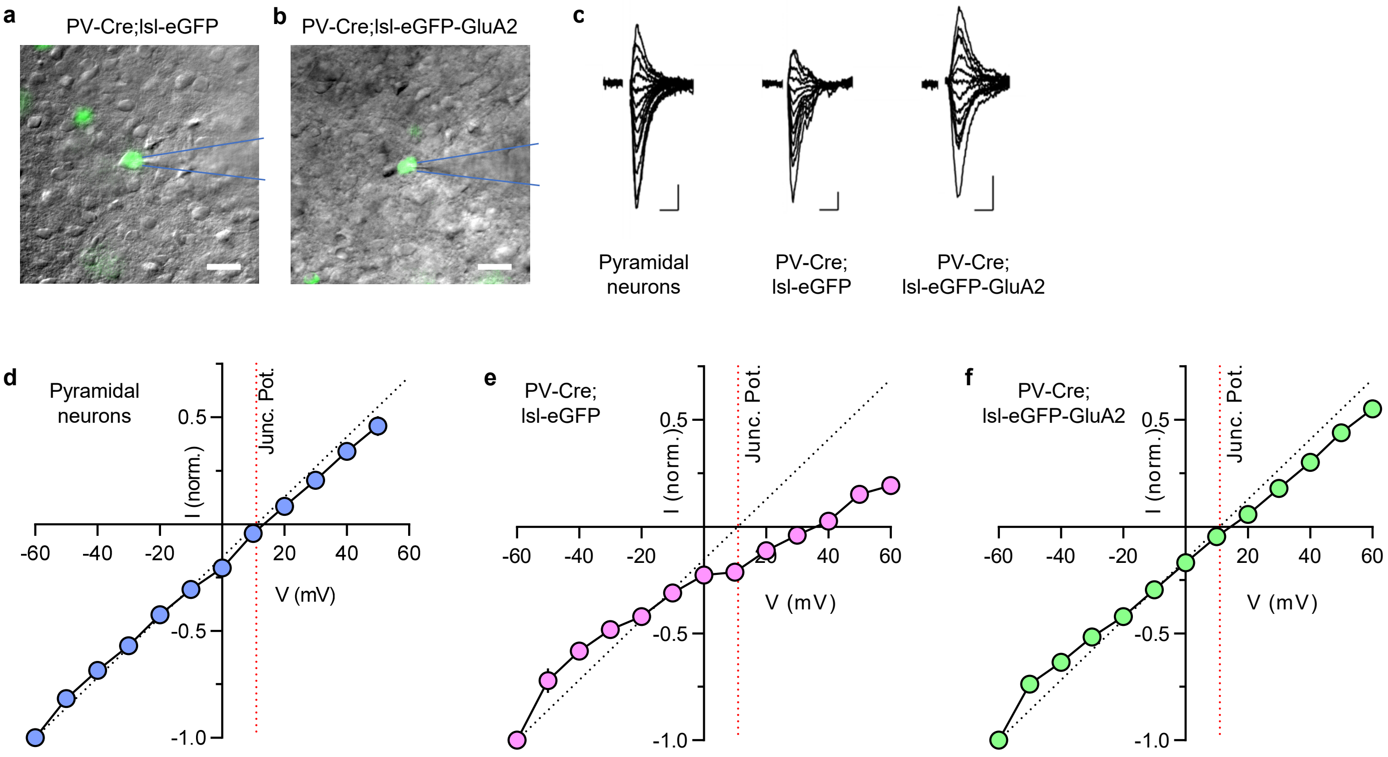


**Extended Data Figure 10 | Conditional transgenic expression of GluA2 in PV interneurons reduces calcium-permeable AMPARs (related to Fig 2d). a-b**, Epi-fluorescence microscopy overlaid upon IR-DIC images of PV-Cre;lsl-eGFP and PV-Cre;lsl-eGFP-GluA2 mice. Scale bars, 20 µm. Layer 2/3 eGFP+ neurons in the visual cortex of PV-Cre;lsl-eGFP and PV-Cre;lsl-eGFP-GluA2 mice were targeted for whole cell patch clamp and rectification measurement. **c**, Example AMPAR-EPSC traces ($V_h$ = −60 mV to 60 mV in 10 mV increments and in the presence of 100 µM AP5) from eGFP or eGFP-GluA2-expressing PV interneurons. Scale bars, 20 pA, 10 ms. **d-f**, Plot of the average I-V relationship for the AMPAR EPSC peak amplitudes of all recorded neurons. Red dotted lines denote the uncorrected junction potential (~11mV). Black dotted lines indicate the expected linear I-V relationship from non-rectifying AMPARs to reveal deviations from linearity. Pyramidal neurons were recorded identically for comparison except for a subset which were measured from $V_h$ = −60 mV to 50 mV. Note that PV control neurons (PV-Cre;lsl-eGFP) display inwardly rectifying AMPAR currents partially reminiscent of the doubly-rectifying CP-AMPAR currents reported previously in heterologous cells[105,106], and additionally display a higher reversal potential, suggesting an alteration of AMPAR channel pore selectivity in cortical PV interneurons.

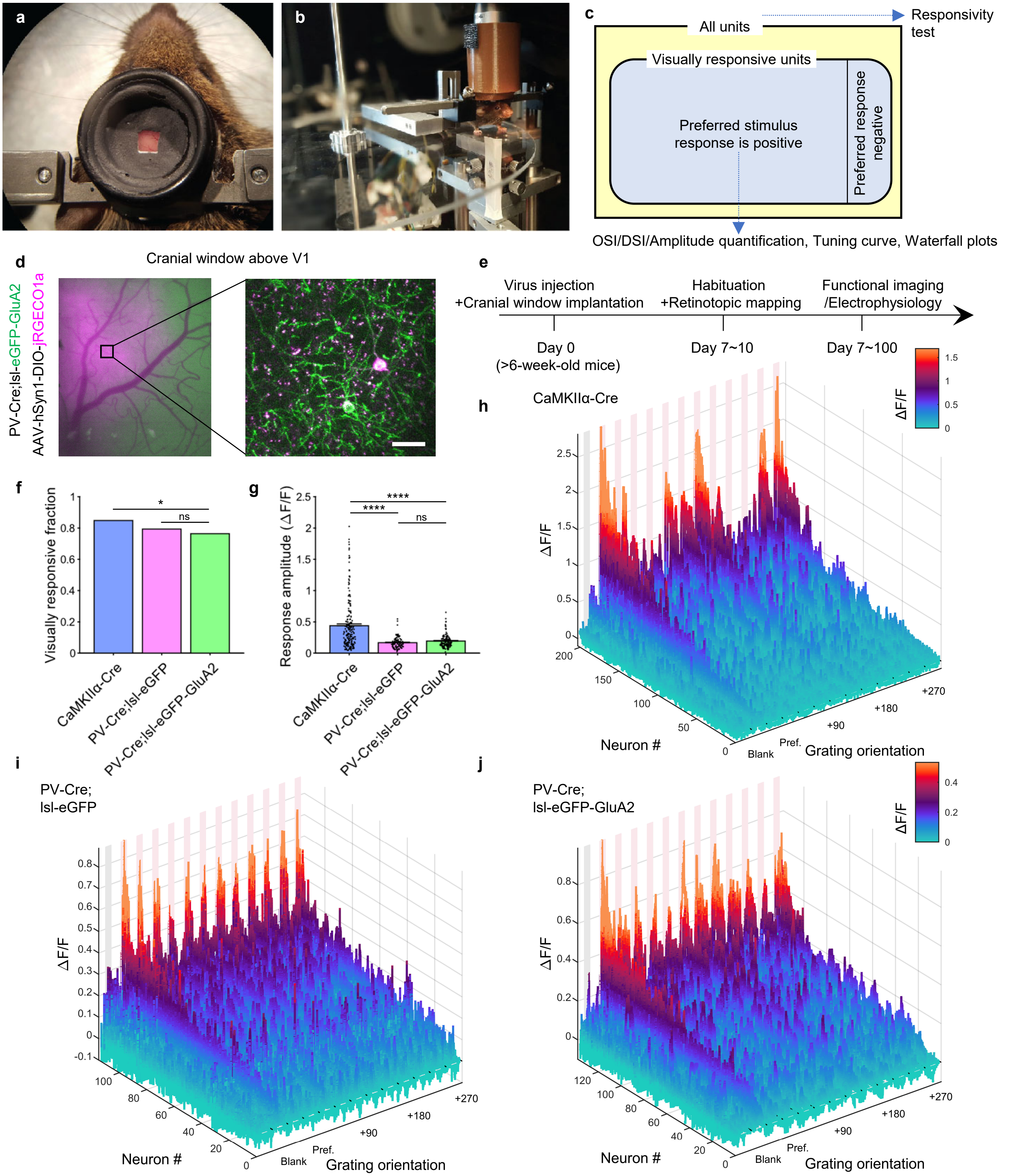


**Extended Data Figure 11 | Awake head-fixed two photon imaging of the visual cortex reveals visual representation changes induced by Cre-dependent eGFP-GluA2 expression in PV interneurons. a**, Reinforced headposts with light-proofing rings to allow visual stimulation during two photon imaging. **b**, Circular treadmill for head-fixed awake imaging. The frame is mounted on a pair of goniometers to allow arbitrary tilt correction. **c**, Venn diagram displaying the hierarchical grouping of visual cortex units based on the responsivity to drifting grating stimuli and valence of largest response. Blue arrows indicate the level at which each analysis or visualization is carried out. **d,** Epifluorescence image of mouse visual cortex through an implanted cranial window (left) and two-photon imaging within the monocular V1 area (right). Scale bar, 20 μm. **e**, Experimental schedule. **f**, Fraction of neurons with statistically significant visual response in CaMKIIα or PV interneurons in each group (0.85, 0.79, 0.76; n = 179/250/395 neurons from 4/4/3 mice, $\chi^2$ = 7.5059, dF = 2, P = 0.0234, Chi-square test). **g**, Average response amplitudes (ΔF/F) of each group. CaMKIIα+ neurons in the CaMKIIα-Cre mice displayed higher amplitude preferred responses compared to PV interneurons in the PV-Cre;lsl-eGFP and PV-Cre;lsl-eGFP-GluA2 group (n = 202/114/137 neurons, $H_{(2)}$ = 66.96, P < 0.0001, KW one-way ANOVA; P < 0.0001 for all post-hoc comparisons with CaMKIIα-Cre, Dunn's multiple comparison correction). **h-j,** Waterfall plots displaying the overall visual response profile of the population of visually responsive neurons with positive preferred stimulus responses in the (**h**) CaMKIIα-Cre, (**i**) PV-Cre;lsl-eGFP, (**j**) PV-Cre;lsl-eGFP-GluA2 groups.

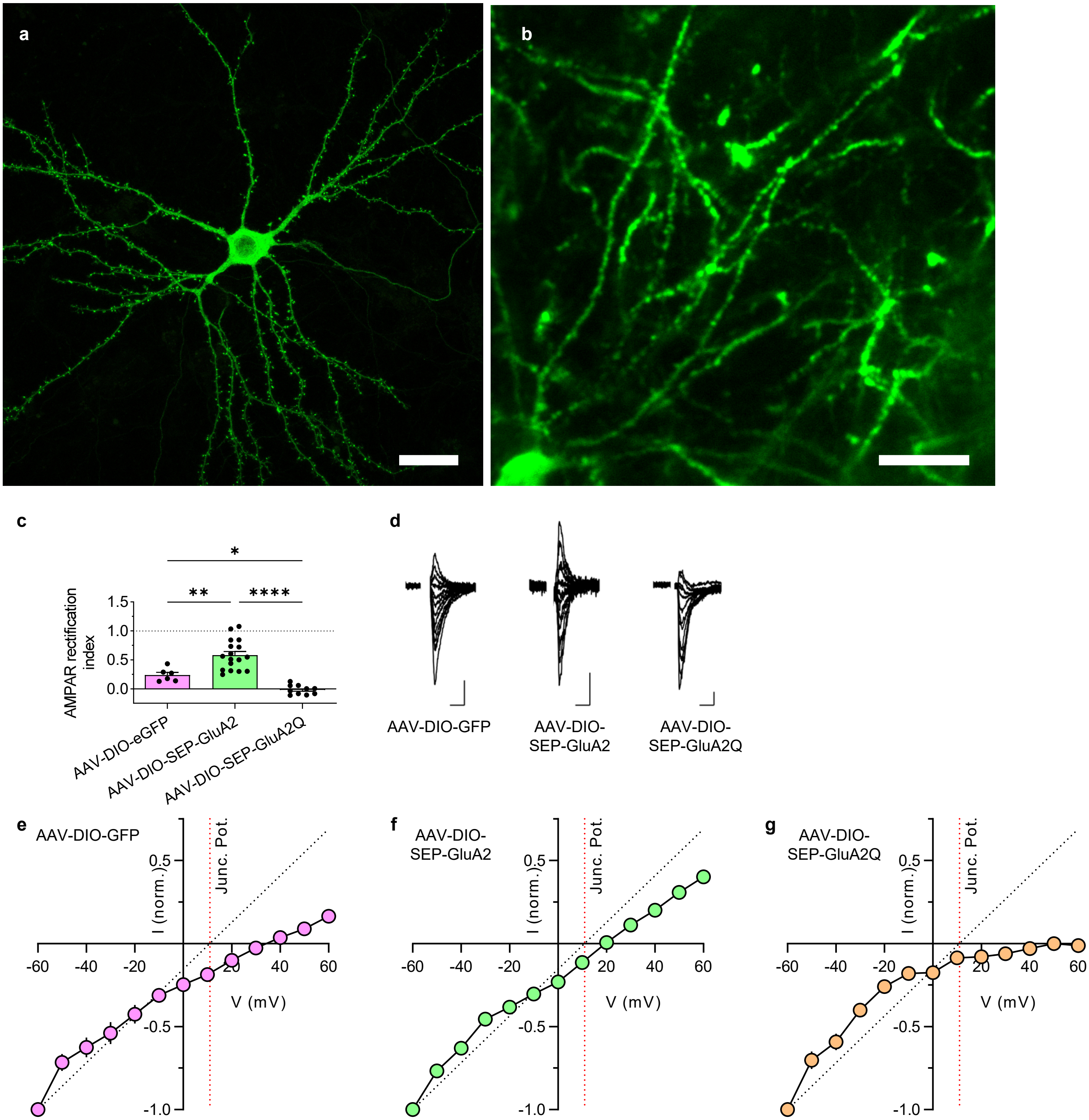


**Extended Data Figure 12 | AAV-mediated Cre-dependent expression of SEP-GluA2 and SEP-GluA2Q in PV interneurons bidirectionally regulates calcium-permeable AMPAR expression. a**, Confocal micrograph of a primary cultured excitatory cortical neuron transfected with FUW-Cre and pAAV-hSyn1-DIO-SEP-GluA2 plasmids. **b**, In vivo two-photon micrograph of a cortical PV interneuron infected with AAV2/9-hSyn1-DIO-SEP-GluA2. Scale bars: 20 µm. **c**, In mice injected with 3 different AAVs, layer 2/3 GFP+ visual cortex neurons were targeted for whole cell patch clamp and AMPAR rectification measurement. Expression of the calcium-impermeable (wild-type) GluA2 subunit (AAV-DIO-SEP-GluA2) relieved AMPAR rectification and increased the rectification index compared to the control group (AAV-DIO-GFP). In comparison, the expression of the calcium-permeable mutant GluA2Q subunit (AAV-DIO-SEP-GluA2Q) resulted in further rectifying AMPAR currents and a lower rectification index (n = 6/17/10 cells from 2/2/3 mice, $P < 0.0001$, one-way ANOVA test; $P = 0.003$ for AAV-DIO-eGFP vs. AAV-DIO-SEP-GluA2, $P = 0.048$ for AAV-DIO-eGFP vs. AAV-DIO-SEP-GluA2Q, $P < 0.0001$ for AAV-DIO-SEP-GluA2 vs. AAV-DIO-SEP-GluA2Q, Tukey's multiple comparison correction). **d**, Example AMPAR-EPSC traces ($V_h$ = −60 mV to 60 mV in 10 mV increments and in the presence of 100 µM AP5) from AAV-expressing PV interneurons. Scale bars, 20 pA, 10 ms. **e-g**, Plot of the average I-V relationship for the AMPAR EPSC peak amplitudes of all recorded neurons. Red dotted lines denote the uncorrected junction potential (~11mV). Black dotted lines indicate the expected linear I-V relationship from non-rectifying AMPARs to reveal deviations from linearity.

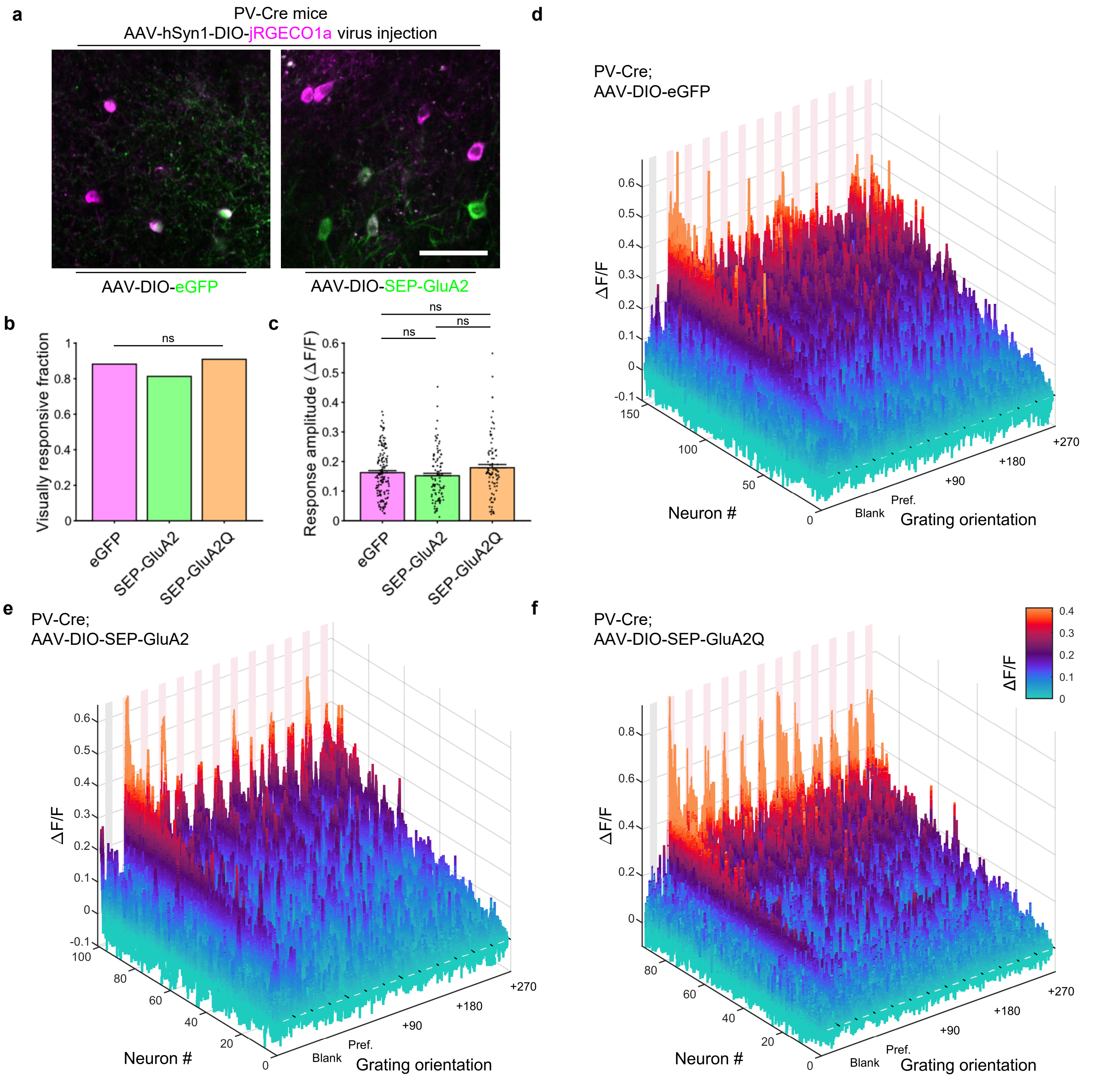


**Extended Data Figure 13 | Visual representation changes induced by sparse viral SEP-GluA2 expression in PV interneurons. a**, Two-photon micrograph of AAV-infected layer 2/3 PV interneurons within the monocular V1 area. Scale bar, 50 µm. **b**, Fraction of neurons with statistically significant visual responses in each group (n = 212/155/110 neurons from 4/4/3 mice, $\chi^2$ = 5.9913, dF = 2, P > 0.0500, Chi-square test). **c**, Average response amplitudes (ΔF/F) of each group's visually responsive neurons were not statistically different (n = 154/100/91 neurons, $H_{(2)}$ = 3.63, P = 0.1628, KW one-way ANOVA). **d-f,** Waterfall plots displaying the overall visual response profile of the population of visually responsive neurons with positive preferred stimulus responses in the (**d**) PV-Cre;AAV-DIO-eGFP, (**e**) PV-Cre;AAV-DIO-SEP-GluA2, (**f**) PV-Cre;AAV-DIO-SEP-GluA2Q groups.

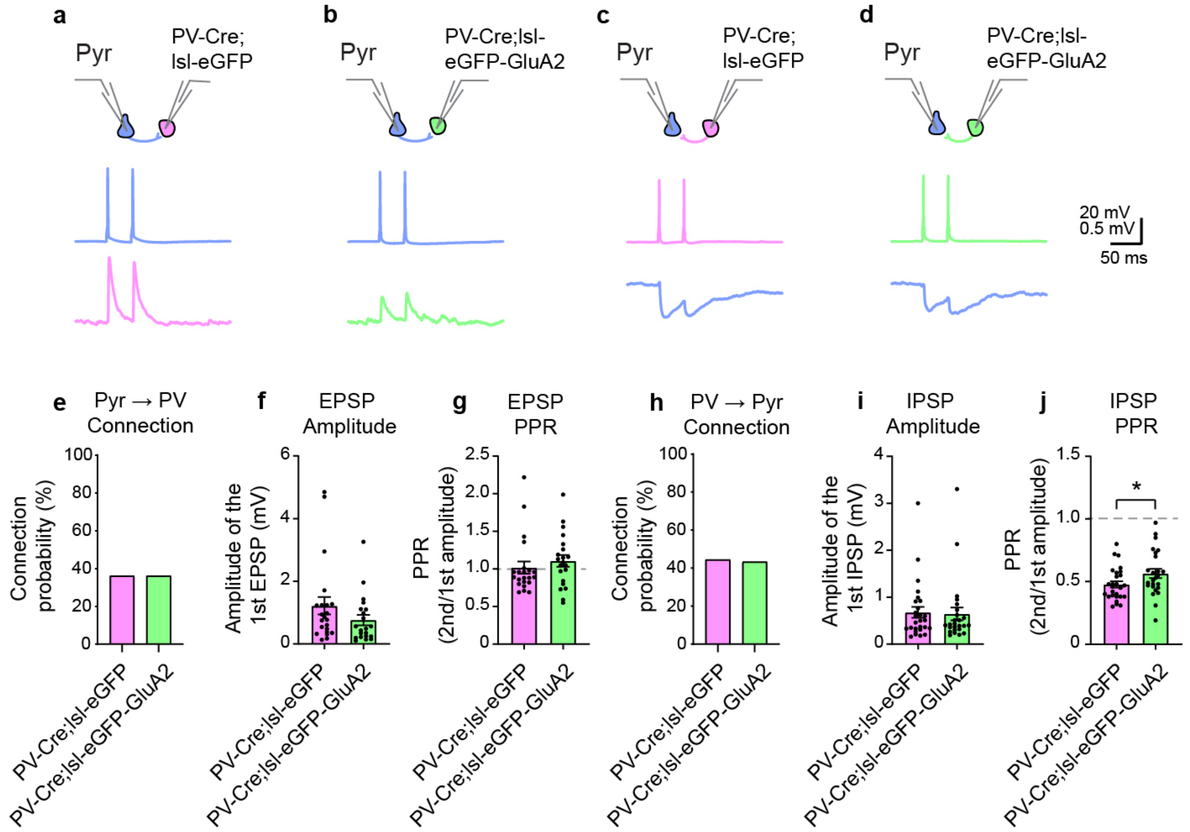


**Extended Data Figure 14. Similar bidirectional connectivity between pyramidal neurons and PV interneurons in PV-Cre;lsl-eGFP and PV-Cre;lsl-eGFP-GluA2 mice. a**, Recording configurations (top), representative action potential traces in presynaptic cells (middle) and voltage response traces in postsynaptic cells (bottom) for Pyr→control PV (**a**), Pyr→GluA2 PV (**b**), control PV→Pyr (**c**), and GluA2 PV→Pyr (**d**) pairs. **e**, The probability of connection for tested Pyr→control PV and Pyr→GluA2 PV connections (Pyr→control PV: 36.7%, n = 22 of 60 tested connections; Pyr→GluA2 PV: 36.8%, n = 21 of 57 tested connections, P = 1, Fisher's exact test). **f**, The amplitudes of the unitary excitatory postsynaptic potentials (uEPSPs) of connected pairs (Pyr→control PV: 1.22 ± 0.30 mV, n = 22 pairs; Pyr→GluA2 PV: 0.76 ± 0.17 mV, n = 21 pairs; P = 0.17702, Mann-Whitney U test). **g**, The paired-pulse ratio (PPR) for connected pairs (Pyr→control PV: 1.02 ± 0.08, n = 22 pairs; Pyr→GluA2 PV: 1.11 ± 0.08, n = 21 pairs; P = 0.13104, Mann-Whitney U test). **h**, The probability of connection for tested control PV→Pyr and GluA2 PV→Pyr connections (PV→Pyr: 45%, n = 27 of 60 tested connections; GluA2 PV→Pyr: 43.9%, n = 25 of 57 tested connections, P = 1, Fisher exact test). **i**, The amplitudes of the unitary inhibitory postsynaptic potentials (uIPSPs) of connected pairs (PV→Pyr: 0.68 ± 0.12 mV, n = 27 pairs; GluA2 PV→Pyr: 0.64 ± 0.14 mV, n = 25 pairs; P = 0.75656, Mann-Whitney U test). **g**, The paired-pulse ratio (PPR) for connected pairs (PV→Pyr: 0.48 ± 0.03, n = 27 pairs; GluA2 PV→Pyr: 0.56 ± 0.04, n = 25 pairs; P = 0.0394, Mann-Whitney U test).

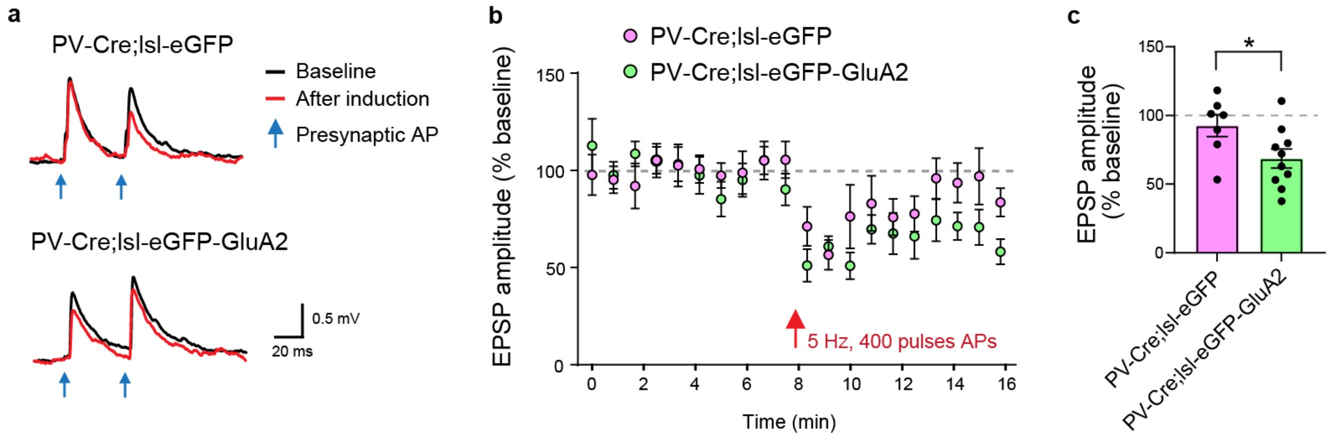


**Extended Data Figure 15. Altered non-Hebbian plasticity in PV interneurons of PV-Cre;lsl-eGFP-GluA2 mice. a**, Representative EPSP traces measured before (black, average of 50 traces) and after (red, average of last 20 traces) anti-Hebbian (AH) plasticity induction (400 presynaptic action potentials at 5 Hz paired with hyperpolarization of postsynaptic PV interneurons to -90 mV). The time points of the presynaptic action potentials for measuring EPSPs are marked with blue arrows. **b**, Normalized EPSP amplitude before and after AH plasticity induction (eGFP, n = 7 pairs from 4 mice; eGFP-GluA2, n = 10 pairs from 9 mice). Red arrow indicates the time point of AH plasticity induction. **c**, A summary graph showing normalized EPSP amplitude of the average of last 20 traces after AH plasticity induction (eGFP, 92.6 ± 8.1%; eGFP-GluA2, 68.6 ± 6.9%, P = 0.03968, *t*-test).

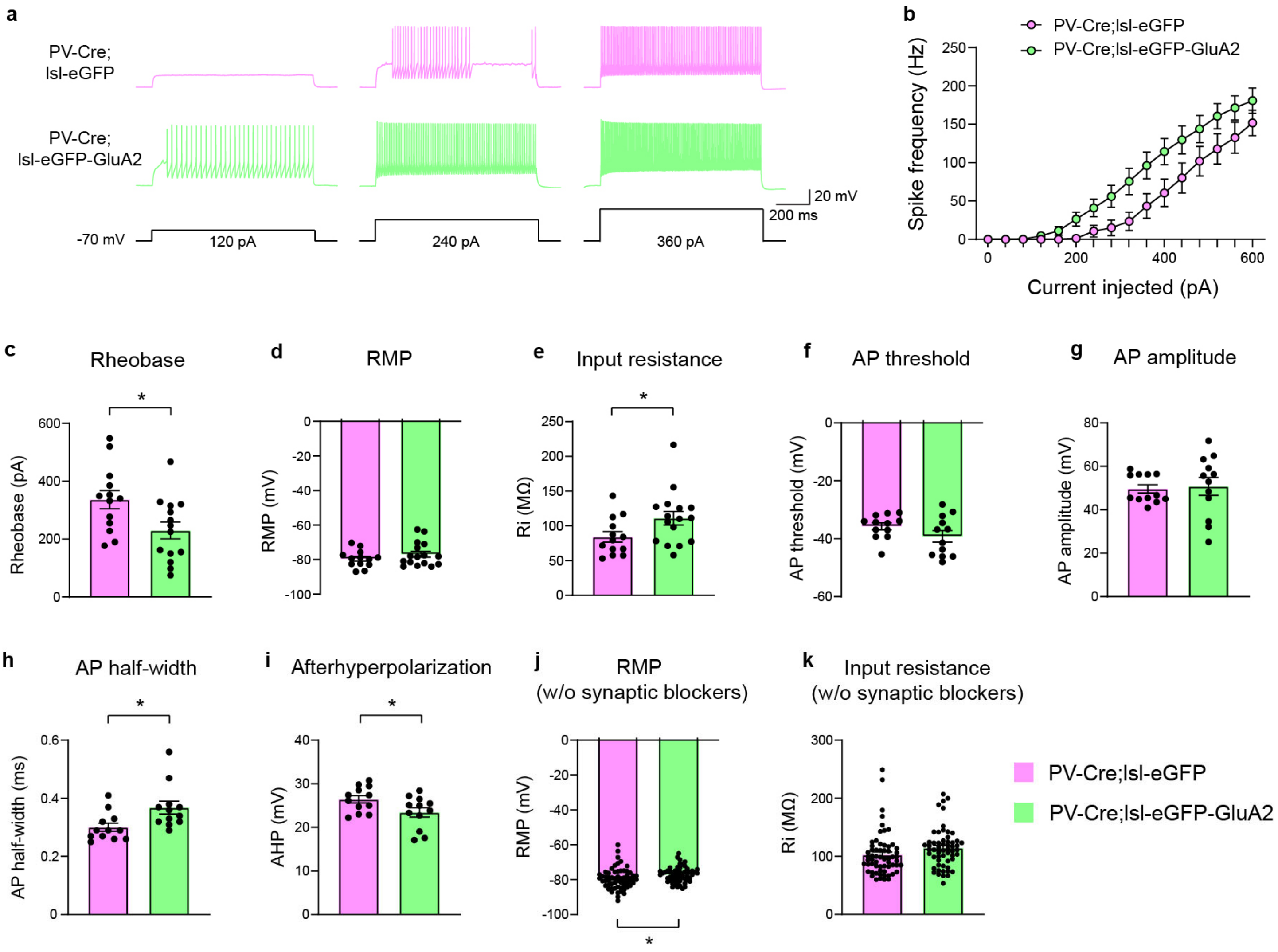


**Extended Data Figure 16. Large increase in intrinsic excitability of PV interneurons in PV-Cre;lsl-eGFP-GluA2 mice.** Representative voltage traces (**a**), and the current–spike frequency relationship (**b**) recorded from control and eGFP-GluA2-expressing PV interneurons (control eGFP: n = 13 cells from 3 mice; eGFP-GluA2: n = 14 cells from 3 mice, P = 0.0025, 2-way ANOVA interaction effect). **c**, The rheobase measured in GluA2-overexpressing PV interneurons was substantially lower than in control PV interneurons (control, 336.31 ± 31.69 pA; GluA2, 230.00 ± 29.35 pA; P = 0.0209). **d**, Resting membrane potential (RMP, control, -79.5 ± 1.37 mV; GluA2, -76.9 ± 1.73 mV; P = 0.4413), in the presence of glutamate and GABA receptor blockers (5 µM NBQX, 5 µM (RS)-CPP, and 10 µM SR95531; applies to panels **a-i**). **e**, Input resistance (Ri, control, 84.3 ± 7.5 MΩ; GluA2, 111.1 ± 9.7 MΩ; P = 0.0453). **f**, Action potential threshold (control, -35.7 ± 1.2 mV; GluA2, -39.2 ± 2.0 mV; P = 0.1501. **g**, Action potential amplitude (control, 49.7 ± 1.9 mV; GluA2, 50.8 ± 4.1 mV; P = 0.509). **h**, Action potential half-width (control, 0.30 ± 0.01 ms; GluA2, 0.37 ± 0.02 ms; P = 0.01). **i**, Afterhyperpolarization (AHP, control, 26.4 ± 0.8 mV; GluA2, 23.4 ± 1.1 mV; P = 0.0387). **j**, Resting membrane potential measured without synaptic glutamate and GABA receptor blockers (RMP, control, -79.5 ± 0.75 mV; GluA2, -77.2 ± 0.58 mV; P = 0.0185). **k**, Input resistance measured without synaptic glutamate and GABA receptor blockers (Ri, control, 102.5 ± 4.7 MΩ; GluA2, 114.3 ± 4.5 MΩ; P = 0.0762). Bars and error bars represent mean ± SEM.

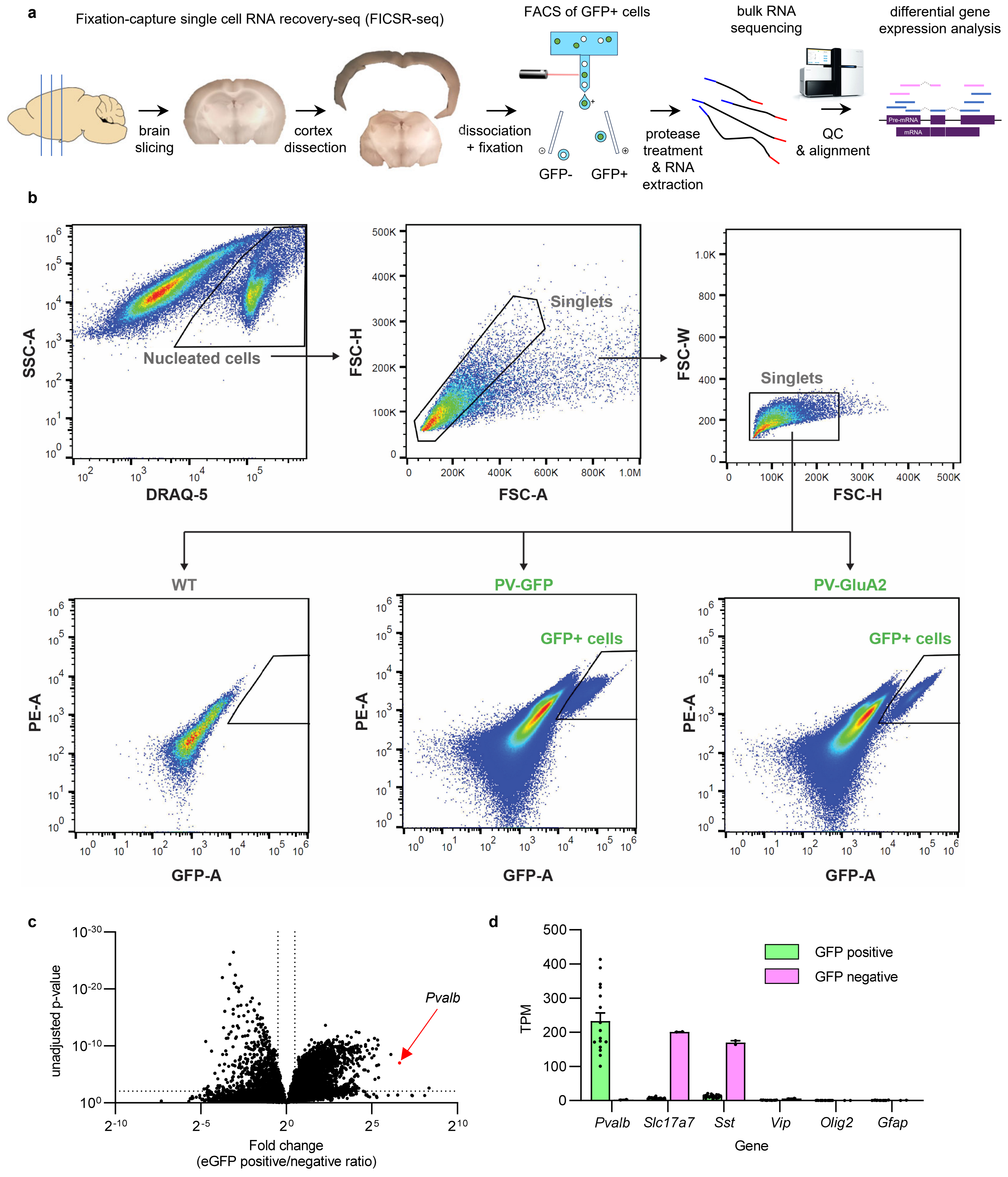


**Extended Data Figure 17 | FACS-assisted bulk RNA-seq of cortical PV interneurons. a,** Overview of the workflow isolating and analyzing mouse cortex PV interneuron mRNA expression. Fixation-capture single cell RNA recovery-seq (FICSR-seq) was used to recover PV interneurons without substantial loss of PV cells during dissociation. After brain slicing, enzymatic dissociation was followed with fixation in 4% PFA and mechanical dissociation. GFP+/DRAQ5+ cells were isolated using FACS and were treated with proteinase K before RNA extraction which removes RNA-binding proteins and increases the yield of intact RNA. The resulting mRNA was sequenced with paired-end Illumina sequencing and analyzed for differential gene expression. **b**, Representative gating diagrams and FACS flowchart. DRAQ-5 was used to sort nuclei-containing cells from debris, and singlets are further sorted into GFP+ and GFP- cells. **c, d**, Bulk RNA-seq reveals ~100-fold enrichment of *Pvalb* mRNA in GFP+ vs GFP- cell samples, validating FACS-based isolation of PV interneuron population. TPM stands for transcripts per million.

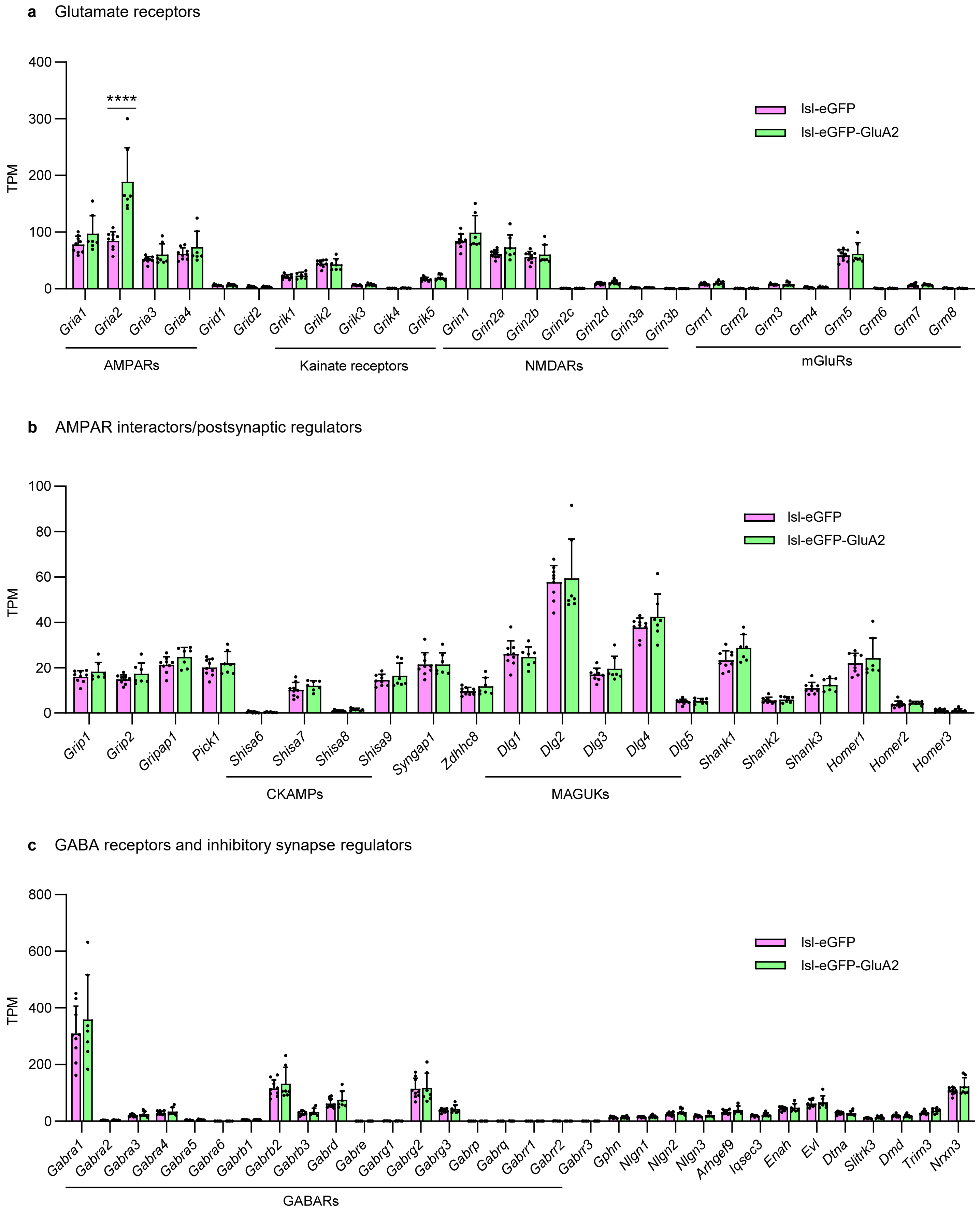


**Extended Data Figure 18 | FACS-assisted bulk RNA-seq of cortical PV interneurons: synaptic gene expression. a,** Gene expression in PV interneurons of each genotype was statistically analyzed with DESeq2 and plotted using TPM (transcripts per million) to visualize expression levels. PV interneurons in PV-Cre;lsl-GFP-GluA2 mice display largely unchanged expression of major glutamate receptor genes compared to PV-Cre;lsl-eGFP control mice, with the exception of *Gria2*, which is overexpressed by roughly 2-fold in these mice (Benjamin-Hochberg adjusted $P < 0.0001$). This is in line with the protein level increases (Fig. 2b) and matches the *Gria2* expression level of typical excitatory neurons. **b**, AMPAR interactor/postsynaptic regulator genes also display unaltered expression.

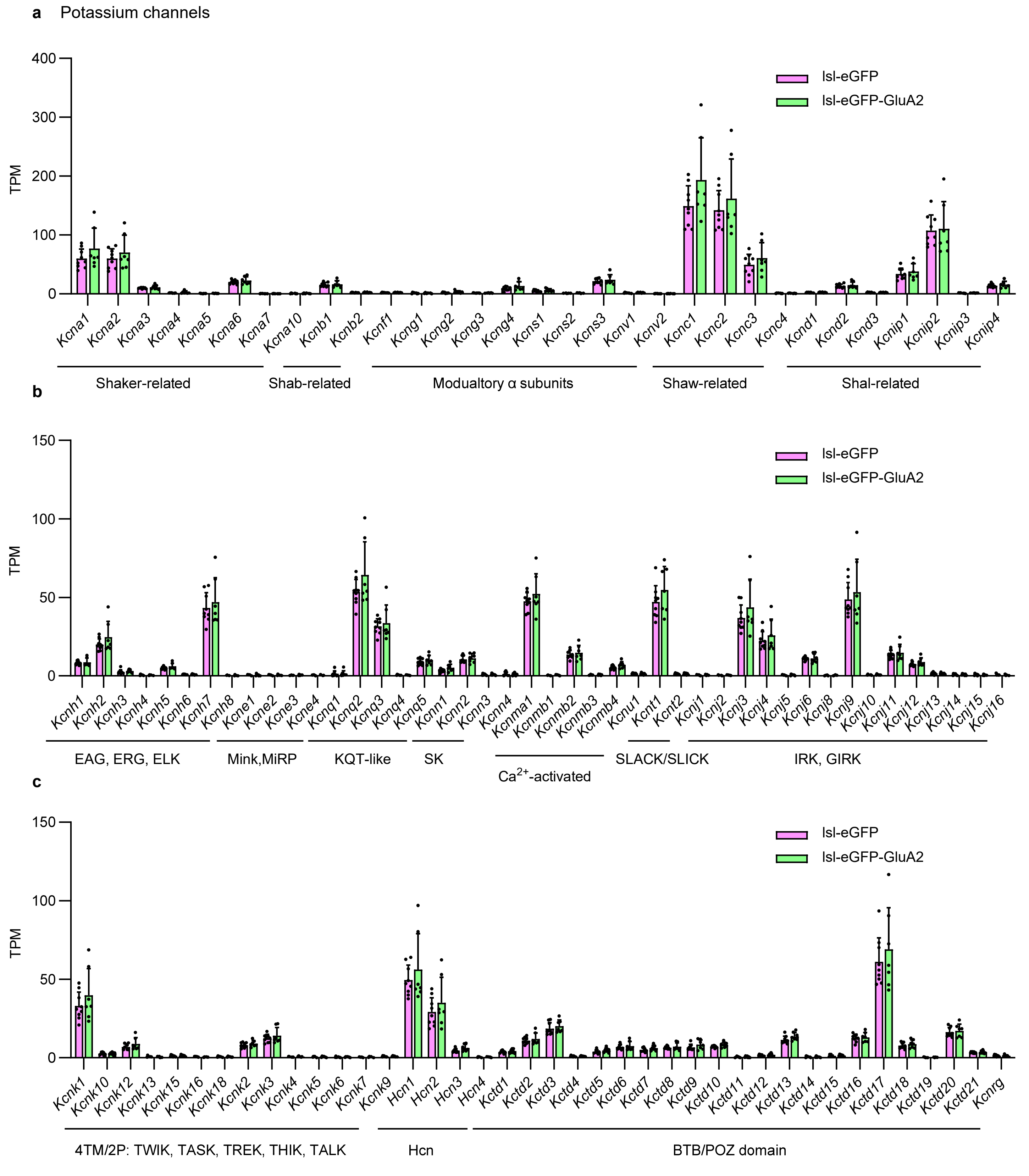


**Extended Data Figure 19 | FACS-assisted bulk RNA-seq of cortical PV interneurons : $K^+$ channel gene expression. a-c,** Gene expression in PV interneurons of each genotype was statistically analyzed with DESeq2 and plotted using TPM (transcripts per million) to visualize expression levels. PV interneurons in PV-Cre;lsl-GFP-GluA2 mice display largely unchanged expression of major $K^+$ channel genes compared to PV-Cre;lsl-eGFP control mice.

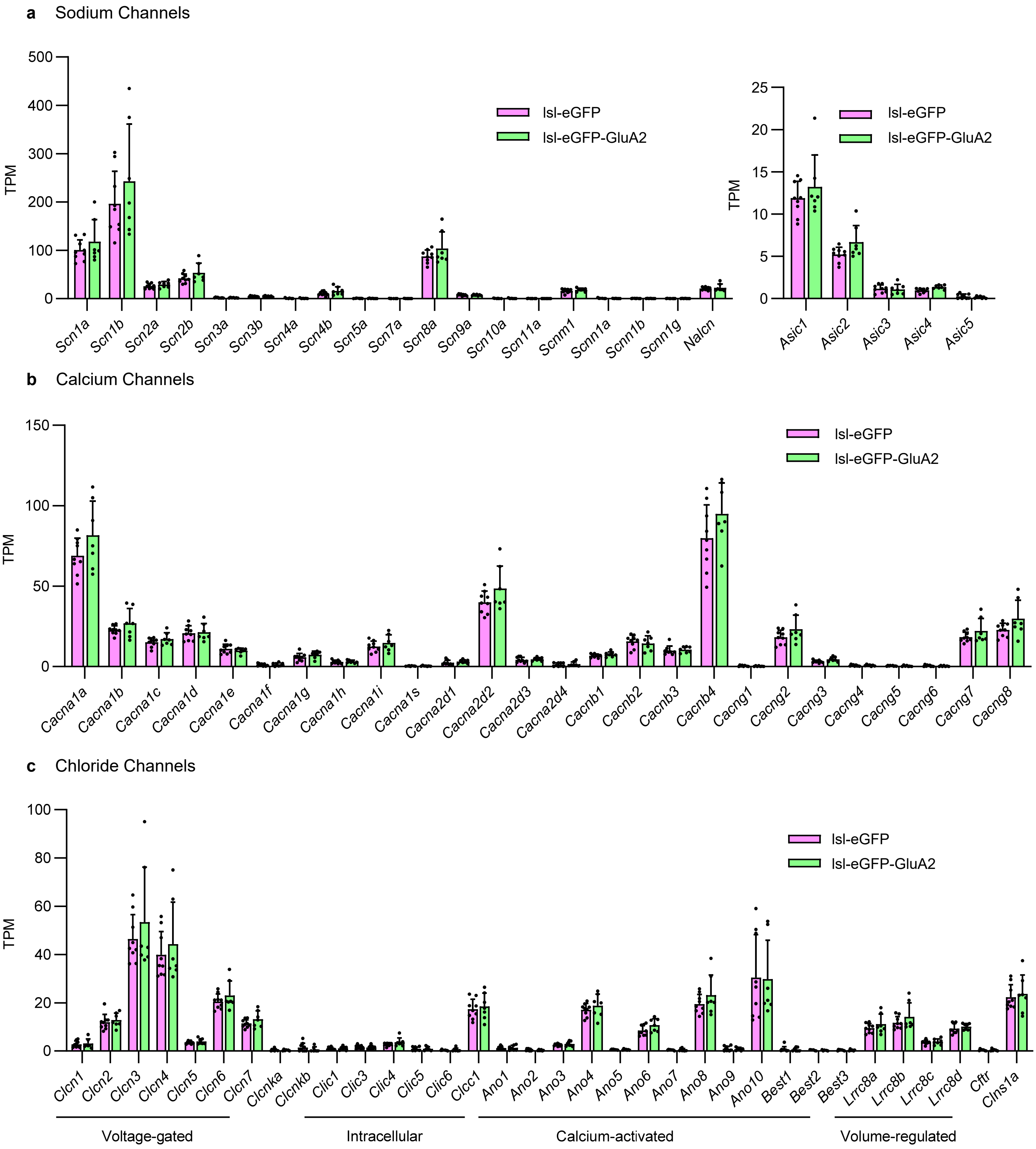


**Extended Data Figure 20 | FACS-assisted bulk RNA-seq of cortical PV interneurons: $Na^+/Ca^{2+}/Cl^-$ channel gene expression. a-c,** Gene expression in PV interneurons of each genotype was statistically analyzed with DESeq2 and plotted using TPM (transcripts per million) to visualize expression levels. PV interneurons in PV-Cre;lsl-GFP-GluA2 mice display largely unchanged expression of major $Na^+$ channel genes (a), $Ca^{2+}$ channel genes (b), and $Cl^-$ channel genes (c), compared to PV-Cre;lsl-eGFP control mice.

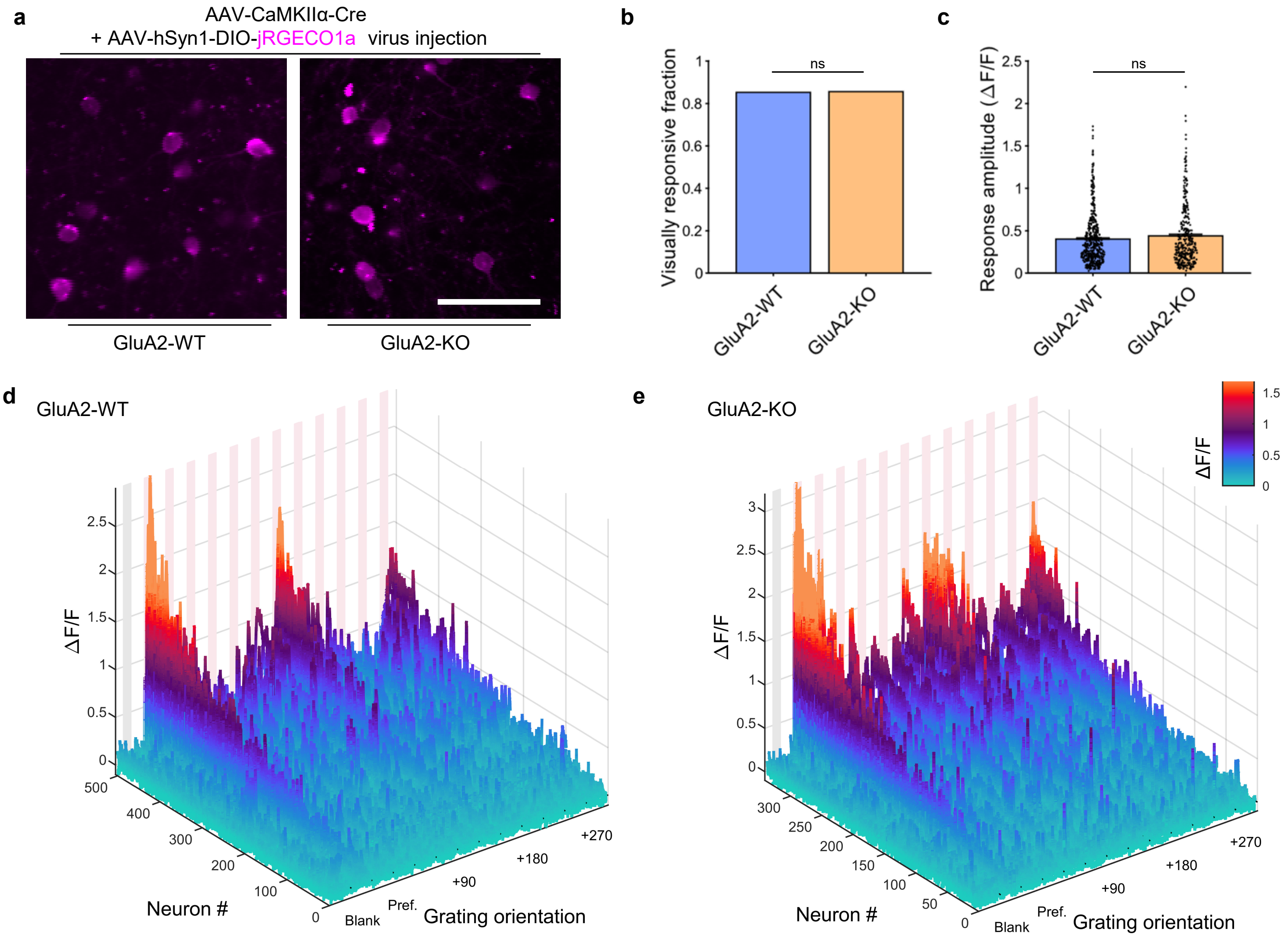


**Extended Data Figure 21 | Visual representation changes in excitatory neurons induced by global GluA2 homozygous knockout. a**, Two-photon micrograph of AAV-infected layer 2/3 excitatory neurons within monocular V1. Scale bar, 50 µm. **b**, Fraction of neurons with statistically significant visual responses in each group (n = 739/500 neurons from n = 3/3 mice, P = 0.9347, Fisher's exact test) . **c**, Average response amplitudes (ΔF/F) of each group was not significantly different (P = 0.4060, Mann-Whitney U-test). **d,e,** Waterfall plots displaying the overall visual response profile of the population of visually responsive neurons with positive preferred stimulus responses in the (**d**) GluA2-WT (+/+), (**e**) GluA2-KO (-/-) groups.

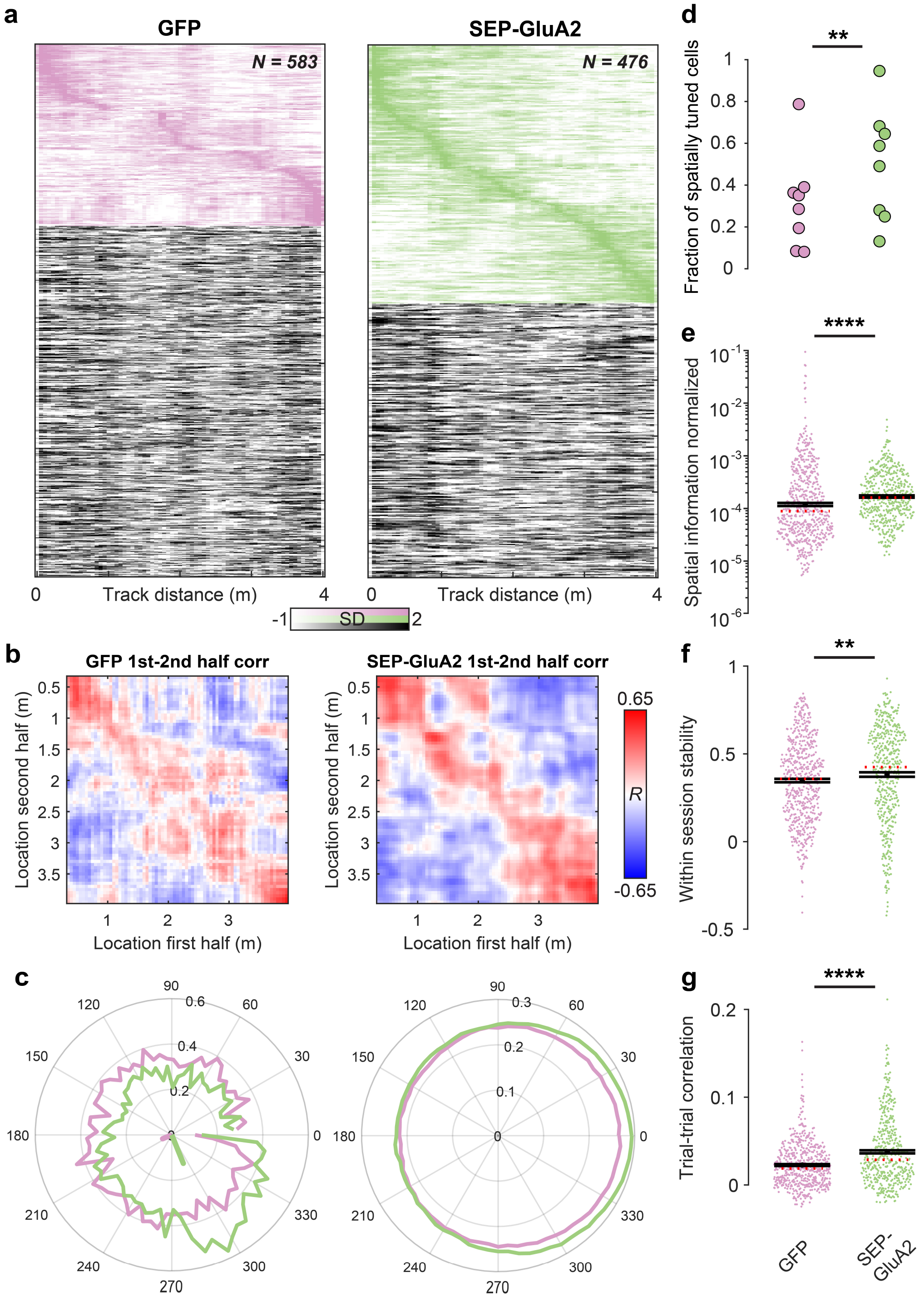


**Extended Data Figure 22 | Increased spatial tuning of hippocampal PV interneurons after removal of CP-AMPARs. a**, Raster plots of $Ca^{2+}$ activity over distance (normalized) for PV interneurons expressing GFP (left) or SEP-GluA2 (right). Activity for cells with significant spatial tuning is shown on top (magenta, green, respectively) and sorted by the peak location of activation. Activity of non-spatially tuned cells is shown below (greyscale). **b**, Average cross correlation of spatial activity profiles of cells between the first and second half of recording sessions (10 runs each). Note the higher and more consistent correlations of spatial tuning in cells expressing GluA2. **c**, *Left,* two representative examples of spatial vector tuning for a GFP (magenta) and SEP-GluA2 expressing (green) PV interneuron. The cell's activity along the track is projected into a circular coordinate system. The orientation and length of the calculated tuning vector in the center correspond to the tuning direction and specificity, respectively. *Right,* Average activity profile for all GFP (magenta) or SEP-GluA2 expressing (green) PV interneurons aligned to their preferred orientation (0°). Note the higher asymmetry and stronger vectorial tuning for GluA2 expressing cells. **d**, Fraction of cells with significant spatial tuning in GFP (magenta) and SEP-GluA2 (green) expressing animals. Dots denote fractions in individual experiments. (n = 8/8 experiments from n = 4/4 mice, P = 0.001, t-test). **e**, Spatial information normalized to activity (see Methods) for GFP (magenta) and SEP-GluA2 (green) expressing cells. Dots denote individual cells. (n = 583/476 cells from n = 4/4 mice, $P = 2.23 \times 10^{-8}$, Wilcoxon rank-sum test) **f**, Same as in (**e**) but for spatial map stability (correlation) between the first and second half of the recording session (P = 0.0026, Wilcoxon rank-sum test). **g**, Same as in (**e**) but for spatial activity map correlations between individual trials of a session ($P = 2.28 \times 10^{-9}$, Wilcoxon rank-sum test). Note the consistently higher stability of spatial representation in PV interneurons expressing GluA2. Black lines in (**e-g**) denote mean ± SEM, and the red dotted line denotes the median. Dots denote values for individual cells.
